# Distinct effects of hypomorphic IFT and dynein-2 skeletal ciliopathy disease alleles on chondrogenic differentiation, ECM composition and wnt signalling in ATDC5 derived cartilage-like organoids

**DOI:** 10.64898/2026.05.21.726337

**Authors:** Anna Klawonn, Himanshu, Zeineb Bakey, Stefan Tholen, Ilona Skatulla, Yong Li, Chiara M. Schröder, Sebastian J. Arnold, Anna Köttgen, Oliver Schilling, Miriam Schmidts

## Abstract

Dysfunction of ciliary intraflagellar transport (IFT) and dynein-2 genes causes severe developmental defects, including skeletal phenotypes characterized by shortened ribs and long bones and polydactyly. Specific gene-phenotype associations suggest individual functions of the different IFT/dynein-2 proteins in development. Since null models disrupt ciliogenesis and hence are not suitable to study individual protein functions, we recreated human hypomorphic disease alleles in IFT-A (IFT43 p.M1V), IFT-B (IFT74 deletion of exon 2), and dynein-2 (WDR60 p.A911V), alongside a WDR60 null model in ATDC5 chondrocyte precursor cells.

Hypomorphic mutants did not show alterations in ciliation efficiency or cilia length but displayed distinct defects in IFT88 localization, indicating impaired intraflagellar transport. Despite altered IFT, Hedgehog signalling responses were variably affected across the different genotypes. Transcriptomic analysis revealed concurrent increases in canonical Wnt signalling and expression of genes related to late skeletal development in WDR60 A911V and IFT74 del ex 2 mutants, but not in IFT43 M1V or WDR60 loss-of-function mutants. These changes were accompanied by alterations in ECM composition. In addition, all hypomorphic mutants showed reduced levels of the non-canonical Wnt ligand WNT5A in ECM proteomic analyses. Interestingly, loss of cilia in WDR60 loss-of-function mutants had only modest effects on chondrogenic differentiation and ECM composition.

Overall, our data provide new evidence of genotype-dependent altered ECM composition as well as dysregulation of canonical and non-canonical Wnt signalling and accelerated chondrocyte differentiation in skeletal ciliopathies. Furthermore, our findings suggest a modulatory rather than essential role of the primary cilium for ATDC5 cell differentiation.

## Introduction

In mammals, the skeleton largely develops through endochondral ossification where a cartilage template forms through coordinated chondrogenesis that is subsequently replaced by mineralized bone^1^. During chondrogenesis, mesenchymal cells condensate and differentiate into extracellular matrix (ECM) secreting chondrocytes through distinct stages^2^. Over time, bone growth occurs at the so-called “growth plates” where chondrocyte proliferation and differentiation is tightly regulated by multiple developmental signalling pathways, including Hedgehog, Wnt, FGFR, BMP, TGF-beta, PTH1R and others^3,4^.

The primary cilium, a microtubule-based cell organelle on almost all cell types, functions as a signalling hub for multiple pathways involved in chondrogenesis^5–7^. Cilium formation and function depend on intraflagellar transport (IFT)^8^, a bi-directional trafficking system along the microtubule based ciliary axoneme. Hereby, anterograde transport from the ciliary base to the tip is powered by kinesin motors^9,10^ and retrograde transport enabled by dynein-2 motors^11^. So-called IFT-A and IFT-B serve as adaptors between the motor and the cargo^12^. While complete loss-of-function of IFT-dynein components results in embryonic lethality in mammals^13–16^, hypomorphic mutations in genes encoding components of the IFT-A, IFT-B, or dynein-2 complexes cause syndromic skeletal phenotypes in humans, collectively termed ciliary chondrodysplasias, characterised by shortened ribs and long bones and frequently accompanied by craniofacial abnormalities and polydactyly^17–22^. Severity of bone shortening and exhibition of extraskeletal features varies considerably depending on which IFT/dynein-2 subunit is defective^22^, suggesting protein specific functions of individual dynein-2 and IFT complex components, that is not yet well understood.

Overall, the role of primary cilia as well as individual IFT-dynein components for skeletal development is still unclear to date. Primary cilia could act as mechanosensors that translate cues from the ECM to developmental signalling^23,24^. While altered cartilage matrix composition and organisation have been described in non-ciliary congenital chondrodysplasias^25,26^, this has not been investigated in ciliary chondrodysplasias.

Likewise, while dysregulated Hedgehog signalling^27–29^ and Wnt signalling^30,31^ are known to affect skeletal development, their role in skeletal ciliopathies, particularly the effects of hypomorphic human disease alleles, remains poorly understood.

Since null alleles in IFTA-, IFTB- or dynein-2 complex genes all severely disrupt ciliogenesis hampering the study of protein specific functions^13,16,28^, we chose to model allele specific effects in ATDC5 chondrocyte precursor cell-derived cartilage-like organoids^32–35^, to study effects on ciliogenesis, IFT88 trafficking, ATDC5 chondrogenic differentiation, cell signalling disturbances as well as potential changes of the ATDC5 cell secreted ECM.

## Results

### Generation of hypomorphic IFT/dynein-2 and Wd60 nonsense alleles in ATDC5 cells

Homozygous WDR60 (dynein-2), IFT43 (IFT-A) and IFT74 (IFT-B) human disease alleles were chosen from reported patients with Short-rib thoracic dysplasia (SRTD)^20,21^, a ciliopathy characterized by severe long-bone shortening and thoracic constriction, and Sensenbrenner syndrome^36^, a ciliopathy with milder skeletal dysplasia and cranioectodermal features. An overview about reported WDR60-, IFT43- and IFT74 human disease alleles is provided in Figure S1/Table S1. Using CRISPR base editing, we obtained three different ATDC5 clones harboring the homozygous *Wdr60* missense allele NM_146039.3 (ENSMUST00000039349.8), c.2732C>T p.Ala911Val equivalent to the human *WDR60* disease allele (NM_018051.5 (ENST00000407559.8), c.2903C>T, p.Ala968Val)^21^, and 3 independent clones harbouring *Ift43* NM_001199843.1 (ENSMUST00000222821.2) c.1A>G corresponding to human *IFT43* p.Met1Val (NM_001102564.3 (ENST00000314067.11) c.1A>G)^36^. Targeting the first coding exon of *Ift74* (NM_026319.3, ENSMUST00000030311.11) (exon 2), using CRISPR resulted in two IFT74 mutant clones derived from wild-type ATDC5 cells and three additional clones generated in iAPEX-tagged^37^ ATDC5 cells (Figure 1, Figure S2) harbouring biallelic exon 2 frameshift alleles, predicted to result in either nonsense mediated decay of RNA or production of a truncated protein lacking exon 2, where the next in-frame exons are used, the first of which locates to the start of exon 3. Western blot analysis indeed confirmed loss of full-length IFT74 and the presence of truncated IFT74 proteins corresponding to using downstream in-frame ATGs in all mutant clones (Figure 1 F), replicating effects of a recurrent human disease allele (*IFT74* g.26959922_26962977delinsTTATTATACTC)^20^.

**Figure 1:**
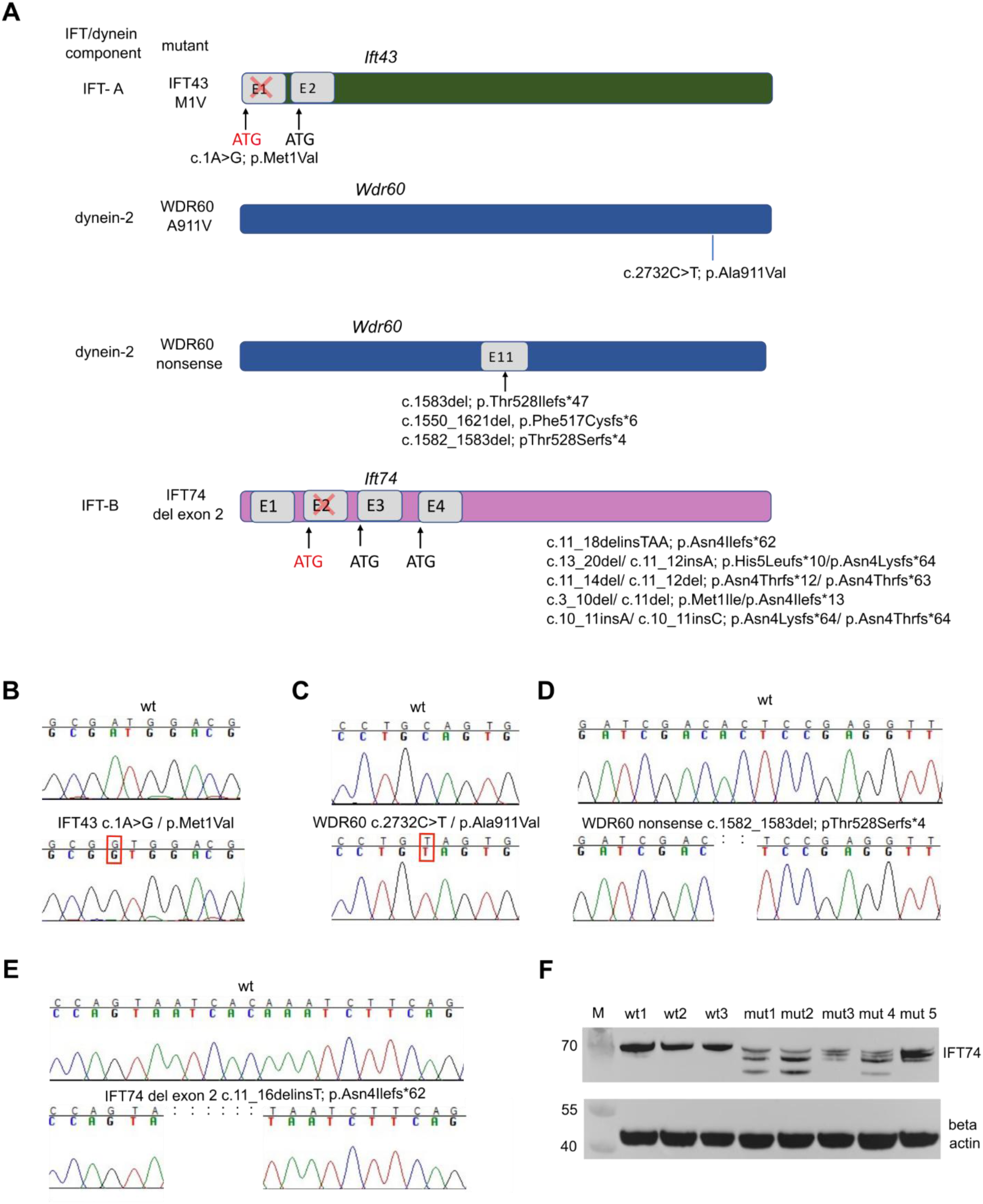
Creation of *Ift43*, *Wdr60* and *Ift74* human disease alleles and *Wdr60* nonsense alleles. (A) Alleles created in this study using CRISPR or CRISPR base editing. (B) Homozygous Ift43 c.1A>G base change leads to translation start at the second in frame ATG. (C) Homozygous *Wdr60* c.2732C>T base change resembles human disease allele. (D) Representative chromatogram of the *Wdr60* nonsense mutation resulting in a frameshift and premature stop codon. (E) Representative chromatogram of *Ift74* mutant leading to translation start at the next in frame ATG. Additional chromatograms from independent IFT74 del ex 2 and WDR60 nonsense clones are shown in Figure S2 and S3. (F) Western blot analysis of three wild-type and five *Ift74* del ex 2 mutant clones used in this study, confirming loss of the original full-length IFT74 protein in mutant clones. Bands of lower molecular weight in mutant samples indicate alternative translation initiation from downstream ATGs.

Finally, to generate a cell line with severe ciliogenesis defects for comparison, we used CRISPR gene editing to obtain *Wdr60* clones carrying biallelic out of frame indels (Figure 1 D, Figure S3, Table S2). Attempts to validate a potential knockout at the protein level using a commercially available antibody was unsuccessful due to no specific bands present in western blot (data not shown).

### Differential effects of WDR60, IFT74, and IFT43 alleles on ciliogenesis and Hedgehog signalling

To assess potential effects of disease alleles on cilia length and ciliation efficiency, all mutant cell lines were investigated using immunofluorescence analyses in comparison to control cells (Figure 2 A, Figure S4). In line with normal ciliation observed in IFT43 M1V patient fibroblasts^36^ and IFT74 del ex 2 mouse fibroblasts^20^, the hypomorphic WDR60 A911V, IFT43 M1V and IFT74 del ex 2 cell lines showed no differences in ciliation efficiency (Figure 2 B-D) or cilia length (Figure 2 F-H) compared to controls. In contrast, *Wdr60* nonsense cells only extended a very short stumpy ciliary axoneme (Figure 2 A, E), in line with previous studies describing cells deficient for dynein-2 components DYNC2H1^16^ and WDR34^38^ extending very short, bulged cilia. Interestingly however, in contrast to our findings, previous reports describe largely unaffected cilia formation and length in WDR60 null hTERT-RPE1 cells^39,40^ and IMCD-3 cells^41^.

**Figure 2:**
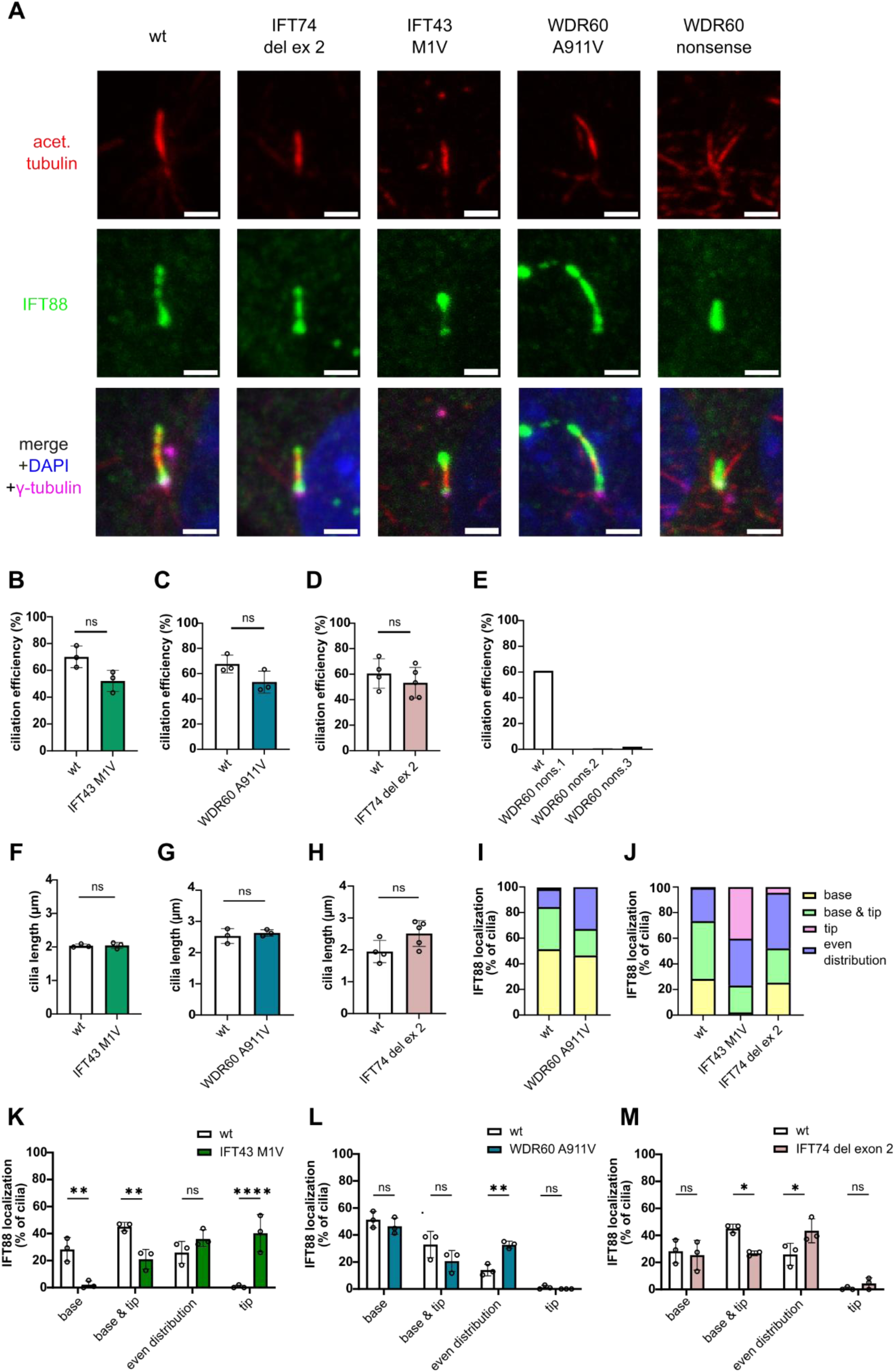
Hypomorphic WDR60, IFT43 and IFT74 mutants display no changes in cilia length or ciliation efficiency, but altered IFT88 localization, while WDR60 nonsense cells fail to extend a ciliary axoneme. (A) Representative immunofluorescence images of cilia of each genotype. Ciliation was induced by serum starvation. Images were taken using a Zeiss LSM 880 Observer confocal microscope. Scale bars: 2 µm. The following analyses of ciliation efficiency, cilia length and IFT88 localization were performed on independent non-confocal images acquired with a Leica Thunder imager under identical experimental conditions. (B-D) Ciliation efficiencies of wild type and hypomorphic mutant clones are similar. For each genotype hundred nuclei of at least three different clones were analysed. Statistical significance was determined using an unpaired two-tailed t-test (n= 3-5). (E) WDR60 nonsense cells do not extend a ciliary axoneme. (F-H) Cilia length of wt and hypomorphic mutant clones is similar. For each genotype 100 cilia of at least three different clones were measured. Data points represent mean cilia length of one clone. Statistical significance was determined using an unpaired two-tailed t-test (n= 3-5). (I-M) Analysis of IFT88 localization in hypomorphic mutant and wild type cells reveals altered IFT88 localization in all hypomorphic mutants compared to wild type. For each genotype 100 cilia of at least three different clones were analysed and assigned to category (I, J). (K, L, M) Comparison of IFT88 localization in hypomorphic mutants vs. wt. Statistical significance was determined using two-way ANOVA with Sidak’s multiple comparisons test (n= 3, \**p* < 0.05, ** *p* < 0.01, *** *p* < 0.001).

To investigate potential intraflagellar transport defects, we examined ciliary IFT88 localization (Figure 2 A, I-M). All mutant lines presented altered IFT88 distribution compared to wildtype cells. WDR60 A911V and IFT74 del ex 2 mutants showed a higher proportion of cilia with uniform IFT88 distribution along the axoneme compared to controls, with a corresponding reduction in the punctate localization pattern, characterized by IFT88 accumulation at the base and tip (Figure 2 I, J, L, M). In contrast, IFT43 M1V cells showed prominent IFT88 accumulation at the ciliary tip and reduced localization at the base (Figure 2 J, K). IFT88 localization was not assessed in WDR60 nonsense cells due the very stumpy appearance of cilia.

Hedgehog signalling is essential for skeletal embryonic development including limb patterning^42^, and disturbances in this pathway can result in polydactyly and shortened bones^43,44^. Hedgehog signalling is mediated through the primary cilium^5^, where in absence of a Hedgehog ligand, full-length Gli proteins are processed into repressor forms^45^ such as Gli3 repressor, preventing target gene expression. Upon pathway activation this processing is stopped resulting in accumulation of full-length Gli3 (Gli3fl) activator^46^. Notably, the balance between Gli3 repressor and Gli3fl expression is crucial for further transcriptional outcomes^47^. IFT contributes to Hedgehog pathway regulation by modulating both its repression and activation^18,48^. Both hypomorphic and null mutations have been reported to result in increased Gli3fl levels together with reduced Gli3 repressor formation^19,27,29,48–50^, probably mediated by reduced processing of Gli3fl to Gli3 repressor^18,27,51^. However, elevated basal Gli3fl levels do not necessarily translate into enhanced Hedgehog pathway activation as shown by in vivo studies using IFT/dynein-2 knockout mouse models indicating that despite increased basal Gli3fl expression, expression of Hedgehog target genes like Gli1 is not induced^16,38^. Moreover, most models show an attenuated or absent response to pathway stimulation. In contrast to IFT/dynein null cells, which mostly show no to very little response to Hedgehog pathway activation^52,53^, hypomorphic cells hereby retain a reduced but detectable response^21,50,54,55^.

We quantified Gli3 repressor / Gli3fl ratios in mutant and wild type ATDC5 clones before and after Hedgehog pathway activation using Smoothened Agonist (SAG)^56^ (Figure 3). WDR60 nonsense cells did not respond to SAG in line with other IFT/dynein-2 null mutants previously reported^52,53^. Interestingly, WDR60 nonsense cells presented with a lower Gli3 repressor / Gli3fl ratio, reflecting concurrently increased Gli3fl levels and reduced Gli3 repressor levels compared to controls both at basal state and after SAG addition, despite not reaching statistical significance (Figure 3 A, Figure S5 G, H, Figure S6 D, E). Likewise, IFT74 del ex 2 mutants showed a lower Gli3 repressor / Glifl ratio before addition of SAG compared to controls (Figure 3 D) with significantly lower Gli3 repressor expression as well as increased Gli3fl expression compared to controls before SAG treatment (Figure S5 E, F). In contrast to WDR60 nonsense and IFT74 mutants, WDR60 A911V, IFT43 M1V responded to SAG treatment with a significant reduction of Gli3 repressor to Gli3fl ratios similar to what we observed in controls (Figure 3). For WDR60 nonsense cells, in line with increased Gli3fl to repressor ratios a basal state, we further observed increased expression of the Hedgehog target gene Gli1^57^ in transcriptomic analyses from day 0 to day 14 (Figure S7 A, Suppl. File S1).

**Figure 3:**
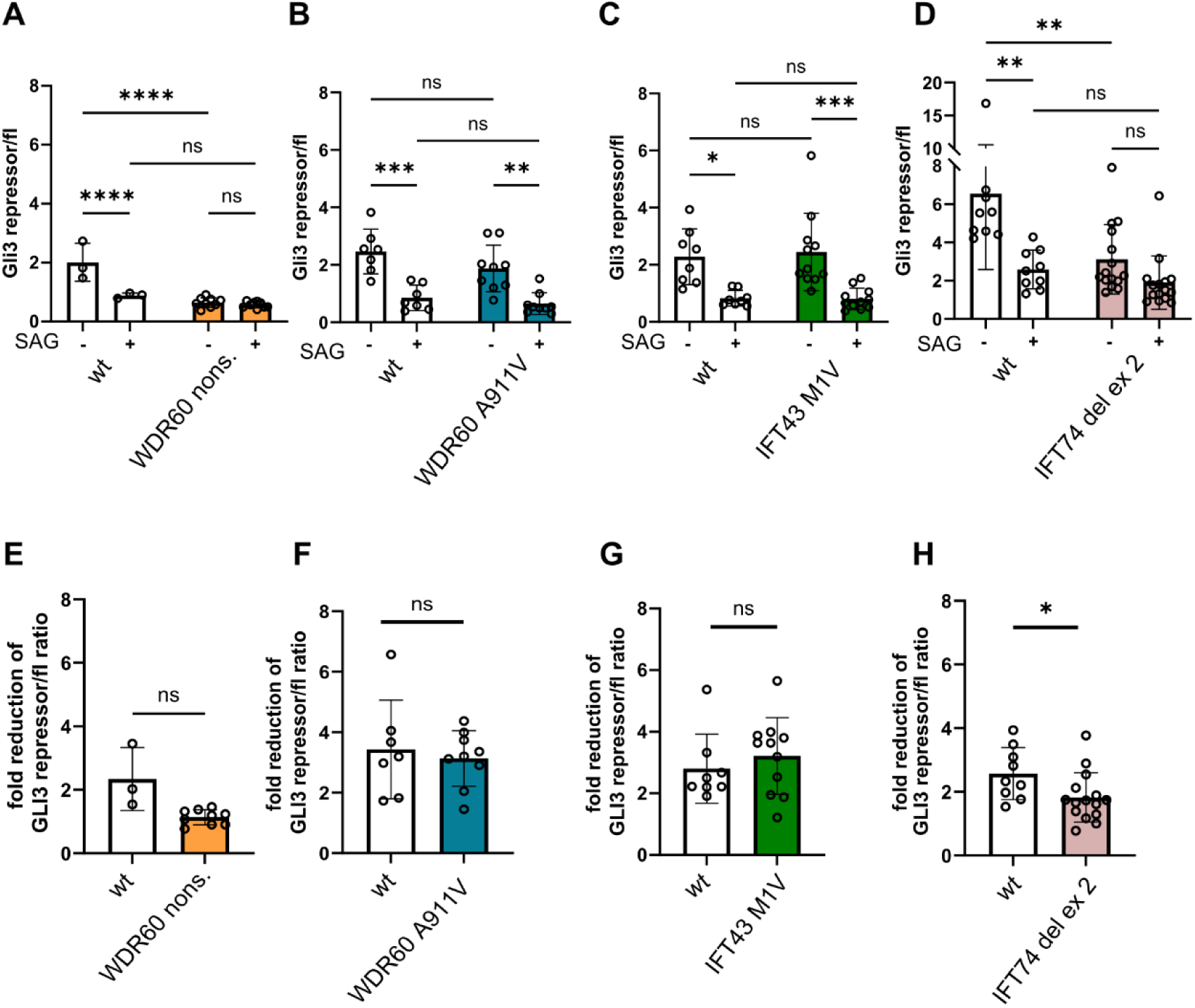
IFT74 del ex 2 mutants show impaired Hedgehog signalling, while WDR60 nonsense cells fail to respond to SAG. (A-D) Quantification of Gli3 full-length (Gli3fl) and Gli3 repressor expression by Western Blot and densitometry. The Gli3 repressor/ Gli3fl ratio before and after SAG treatment was calculated for each sample and pooled across biological replicates containing multiple wild-type and mutant samples per experiment. Bars represent mean ± SD from at least three independent experiments. Statistical significance was determined using two-way ANOVA with Tukeýs multiple comparisons test (n= 3-15, \**p* < 0.05, ** *p* < 0.01, *** *p* < 0.001). (E-H) Fold reduction of the Gli3 repressor/Gli3fl ratio in response to SAG treatment. In WDR60 nonsense clones (E), no statistically significant difference was detected, likely due to the limited sample size of wild-type clones. However, a lack of SAG response in WDR60 nonsense cells is consistently observed across independent experiments (see also Fig. S5 G, H and Fig. S6 D). IFT74 del ex 2 mutants show an impaired response compared to wild-type controls (H). Statistical significance was determined using unpaired t-test (n=3-15, \**p* < 0.05).

### Effects of IFT/dynein-2 dysfunction on ATDC5 chondrogenic differentiation

We next investigated potential shared molecular mechanisms underlying skeletal ciliopathy phenotypes caused by three IFT/dynein-2 disease alleles by analysing gene expression and ECM composition in differentiated mutant and control ATDC5 chondrocyte precursor cells. ATDC5 cells, derived from AT805 mouse teratocarcinoma cells^32^, represent a suitable, established in vitro model of chondrogenesis, progressing through proliferative and hypertrophic stages and secreting cartilage-like ECM upon differentiation, although the transitions between these stages are not sharply defined^33,34,58^. Mutant and wildtype ATDC5 lines were differentiated into cartilage-like organoids as previously described^59^, and RNA and ECM samples were collected at multiple time points during differentiation. We first focussed on differentially expressed genes and proteins across all mutants versus controls (Table 1 and 2, Figure S8, S9, Suppl. File 2 and 3) to explore common effects. To restrict the proteomic analysis to relevant ECM components, only proteins annotated to GO:0031012 (extracellular matrix) were considered.

**Table 1:**
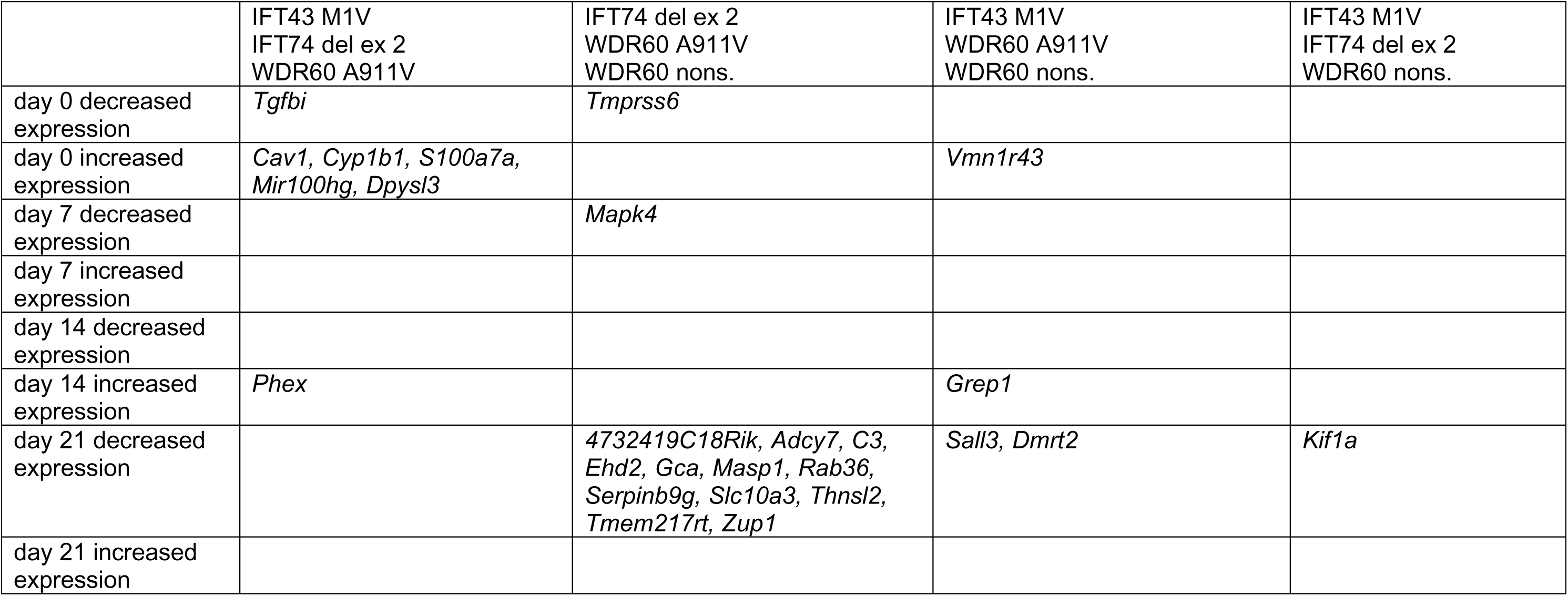
Differentially expressed genes consistently regulated in mutants vs. wt.

All mutant and wild type clones showed comparable proliferation (Figure S10), produced proteoglycan- and collagen-rich ECM (Figure S11) and expressed typical chondrogenic markers during differentiation (Figure S12-S14). To identify shared differentially expressed genes and proteins across mutants versus controls, differential expression analyses were first performed separately for each mutant versus controls at day 0, day 7, day 14, and day 21 in the transcriptomic and proteomic datasets independently. The resulting sets of differentially expressed genes and proteins were then compared across mutants at each time point. Overall, these analyses revealed little overlap for individual genes and proteins between all mutant clones versus controls in both transcriptomic and ECM proteomic datasets (Table 1 and 2). No genes or proteins were shared as differentially expressed in proteomic as well as transcriptomic analyses across all mutant lines relative to controls at any given time point, except for ANG2 protein, which was reduced in all mutants compared to controls at day 14 (Table 2).

**Table 2:** Differentially expressed proteins consistently regulated in mutants vs. wt.

|  | IFT43 M1V<br>IFT74 del ex 2<br>WDR60 A911V | IFT74 del ex 2<br>WDR60 A911V<br>WDR60 nons. | IFT43 M1V<br>WDR60 A911V<br>WDR60 nons. | IFT43 M1V<br>IFT74 del ex 2<br>WDR60 nons. | IFT43 M1V<br>IFT74 del ex 2<br>WDR60 A911V<br>WDR60 nons. |
| --- | --- | --- | --- | --- | --- |
| day 7 decreased expression | ADAMTSL4, FMOD, **WNT5A** |  | ALB, SERPINE1 |  |  |
| day 7 increased expression |  |  |  |  |  |
| day 14 decreased expression | **WNT5A** | EMILIN1 | IGFBP7, TNFRSF11B |  | ANG2 |
| day 14 increased expression |  |  |  |  |  |
| day 21 decreased expression |  |  | COL9A1, SERPINE1, TNFRSF11B | IGFBP7 |  |
| day 21 increased expression |  |  | CST3 |  |  |

Looking at transcriptomics datasets separately from proteomic datasets, *Phex*, a phosphate-regulating endopeptidase implicated in bone mineralization and mutated in X-linked hypophosphatemic rickets^60^, was the only gene with consistently elevated expression levels across all hypomorphic mutants at day 14. On day 0, *Tgbfi*, a positive regulator of bone size and ECM homeostasis^61^, showed lower gene expression across all hypomorphic mutants (Table 1).

The ECM proteomic analysis revealed reduced protein expression in all hypomorphic mutants at day 7 for ADAMTSL4 (Disintegrin and metalloproteinase with thrombospondin motifs 4), a secreted glycoprotein with roles in microfibril assembly^62^, and FMOD (Fibromodulin) a small leucine rich proteoglycan fibromodulin regulating collagen fibril diameter^63,64^ (Table 2). In addition, WDR60 nonsense, IFT43 M1V and WDR60 A911V mutants showed reduced expression of osteoprotegerin (TNFRSF11B), a RANKL decoy receptor regulating bone density^65^ at day 14 and day 21.

Interestingly, the non-canonical Wnt ligand WNT5A was the only protein annotated in the GO term “extracellular matrix” (GO:0031012) consistently reduced across all hypomorphic mutants versus controls on day 7 and 14 (Table 2, Figure 4, Suppl. File 3). In WDR60 nonsense mutants, no differential expression of Wnt5a was found, neither in transcriptomic, nor proteomic analyses (Suppl. File 4 and 5). This could indicate differential effects of hypomorphic disease alleles versus complete loss of cilia. Interestingly, in contrast to proteomics analysis, transcriptome analysis did not reveal significant differences in WNT5A expression between mutants and controls (Figure S15-S18, Suppl. File 5) with the exception of IFT43 M1V mutants, where WNT5A expression was reduced at RNA level at day 0 (Figure 5 E), but not at later stages (Figure S15). Wnt5a is a key regulator of planar cell polarity (PCP), crucial for skeletal development^66,67^, especially for maintaining columnar arrangement in the growth plate^68–70^. It exerts stage-dependent effects on chondrogenesis, promoting early differentiation while inhibiting hypertrophy^31,71,72^. In humans, Wnt5a loss of function causes Robinow syndrome, a short-limbed dwarfism resembling ciliopathy phenotypes^66,73^.

**Figure 4:**
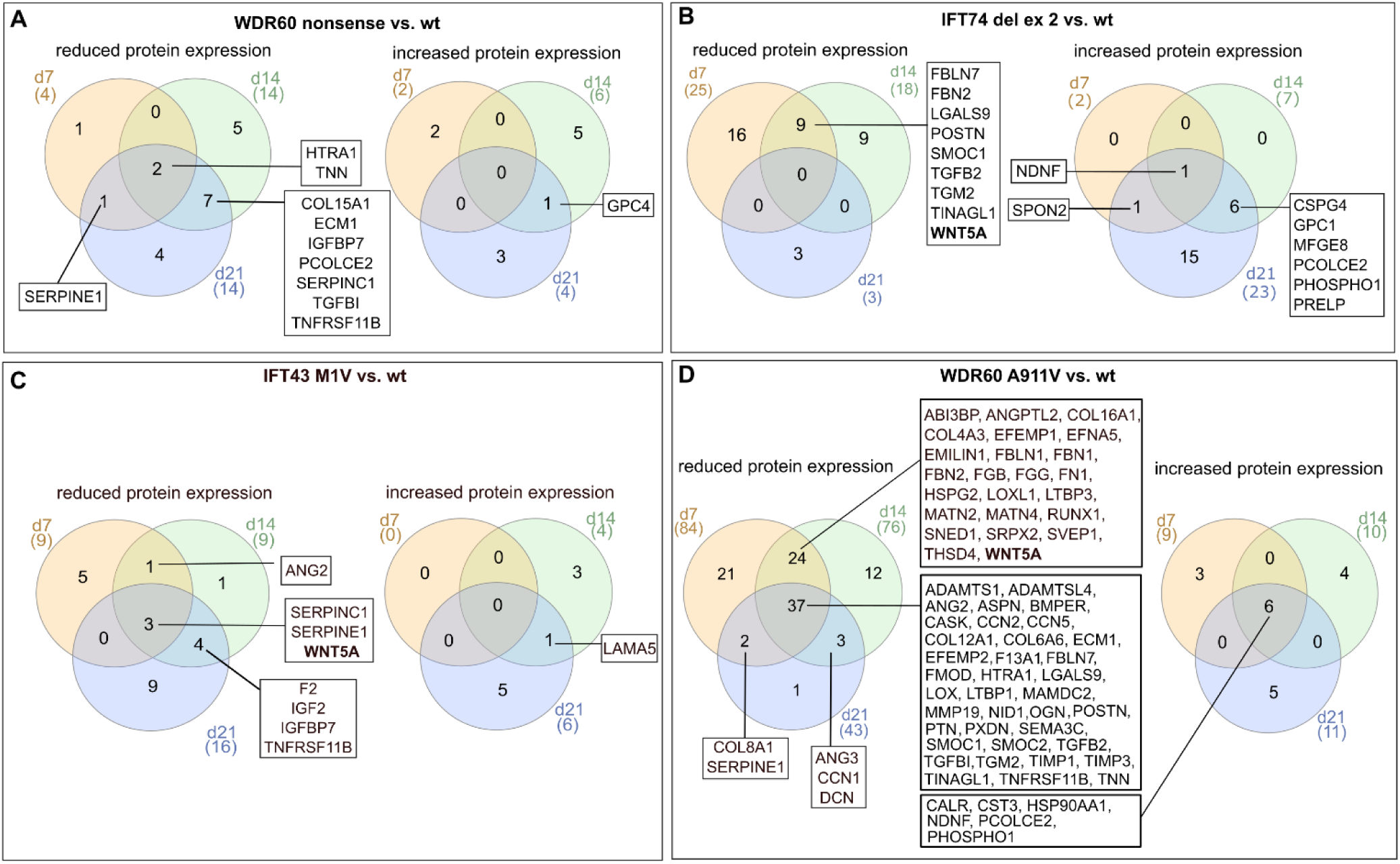
ECM Proteomics reveals decreased expression of WNT5A in all hypomorphic mutants. (A-D) Venn Diagrams illustrating ECM proteins with increased or decreased expression in WDR60 nonsense (A), IFT74 del ex 2 (B), IFT43 M1V (C) and WDR60 A911V (D) mutant clones compared to wild type at indicated time points. Only proteins annotated in GO:0031012 extracellular matrix were considered. WDR60 A911V mutants exhibit the most pronounced changes in ECM composition, whereas IFT74 del ex 2 mutants show moderate alterations. In contrast, WDR60 nonsense and IFT43 M1V mutants display only minor changes. Notably, all hypomorphic mutants show decreased WNT5A expression at day 7 and day 14. Complete lists of intersections are provided in Suppl. File S3. See also Suppl. File S4.

**Figure 5:**
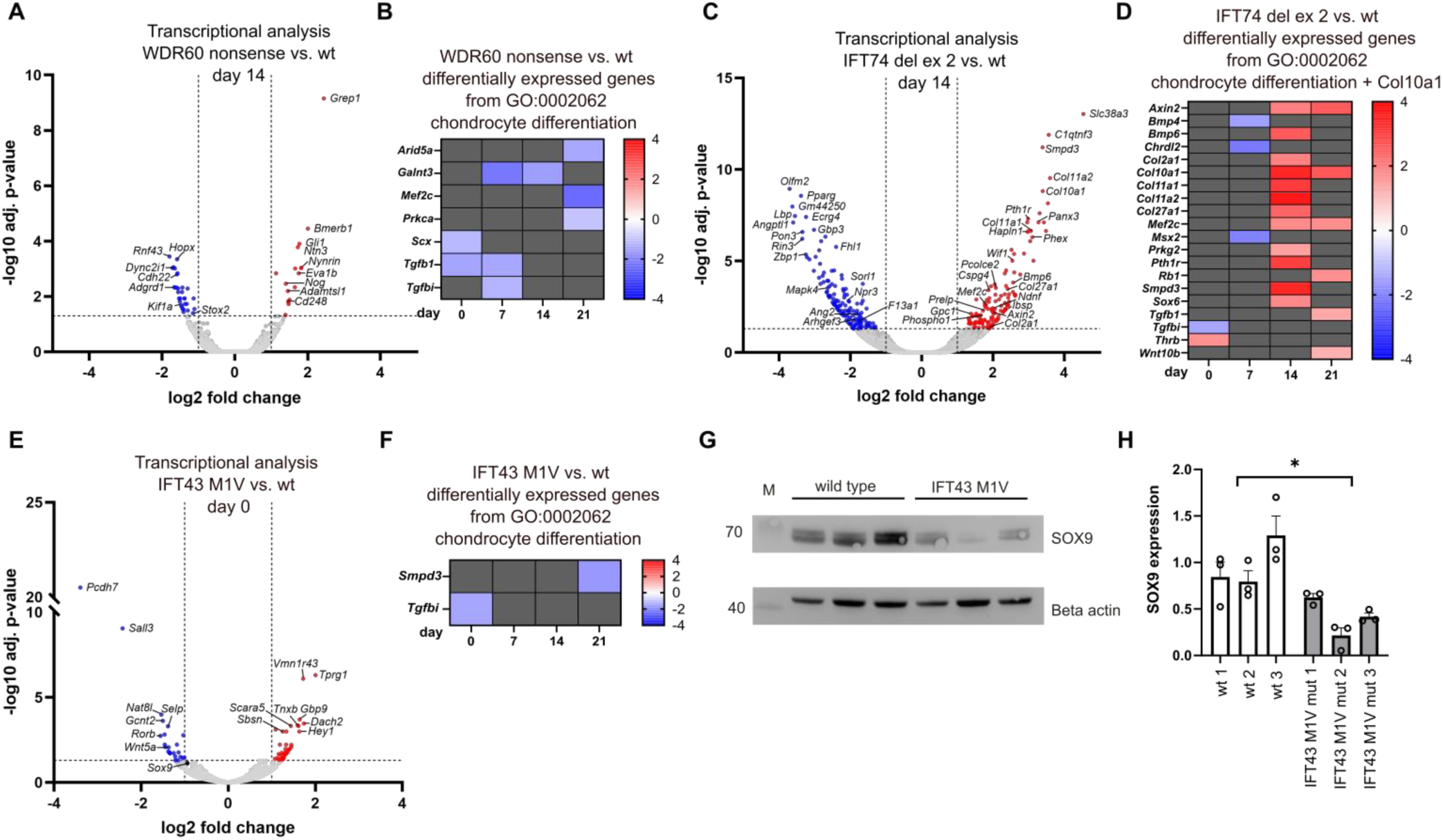
Limited effects on chondrocyte differentiation in IFT43 M1V and WDR60 nonsense mutants in contrast to increased transcriptional changes in IFT74 del ex 2 mutants and reduced SOX9 expression in IFT43 M1V. (A, C, E) Volcano plots showing differentially expressed genes in WDR60 nonsense (A), IFT74 del ex 2 (C), and IFT43 M1V (E) mutant cells compared to wild-type controls at the indicated time points. (B, D, F) Heatmaps of differentially expressed genes associated with GO:0002062 (chondrocyte differentiation) in WDR60 nonsense (B), IFT74 del ex 2 (D), and IFT43 M1V (F) mutant cells compared to wild type. WDR60 nonsense and IFT43 M1V samples show only few significant changes, whereas IFT74 del ex 2 mutants display a higher number of differentially expressed genes, increasing after day 14. In IFT43 M1V cells, only two genes (Smpd3 and Tgfbi) were significantly altered. (G) Representative Western blot analysis of SOX9 with β-actin as loading control in IFT43 M1V mutants and wild type cells at day 0. (H) Densitometric quantification shows significantly reduced SOX9 expression in IFT43 M1V mutants (Student’s t-test; mean ± SEM; p < 0.05; n = 3). In all volcano plots, genes were color-coded based on log2 fold change thresholds of ±1 and an adjusted p-value ≤ 0.05. See also Figure S15, S16, S18.

### Shared differential expression of skeletal development genes, Wnt signalling pathway genes and changed ECM composition between WDR60 A911V and IFT74 del ex 2 mutants

As few individual genes were consistently differentially expressed across mutants, we performed GO analysis of genes with increased or decreased expression for each mutant vs. control at each time point to identify shared functional patterns (Suppl. File 6). IFT43 M1V and WDR60 nonsense mutants generally showed few differentially expressed genes and correspondingly few or no significant GO terms, whereas IFT74 del ex 2 and WDR60 A911V mutants exhibited increased expression of genes related to Wnt signalling and skeletal development (Figures 5-7, S16, S17, Suppl. File 6).

Particularly at day 14, IFT74 del ex 2 and WDR60 A911V mutants shared numerous genes with increased expression compared to control, linked to later stages of endochondral bone formation, with GO terms predominantly associated with “bone mineralization” and “endochondral ossification” (Figure S19, Table S3). These included the hypertrophy-inducing transcription factor Mef2c^74^, mineralization enzymes Alpl and Phospho1^75,76^ as well as *Panx3, Smpd3* and *Pth1r* which have crucial roles in endochondral bone formation^77–79^. Notably, ATP/Ca^2+^ channel PANX3 and sphingomyelin phosphodiesterase SMPD3 are specifically implicated in chondrocyte hypertrophy (Figure 5, 6, S19). As a regulator of the Golgi-mediated secretion of ECM components, SMPD3 deficiency causes chondrodysplasia^80,81^. In contrast, genes with reduced expression in WDR60 A911V and IFT74 del ex 2 mutants at day 14 were primarily associated with stress response pathways (Table S4).

**Figure 6:**
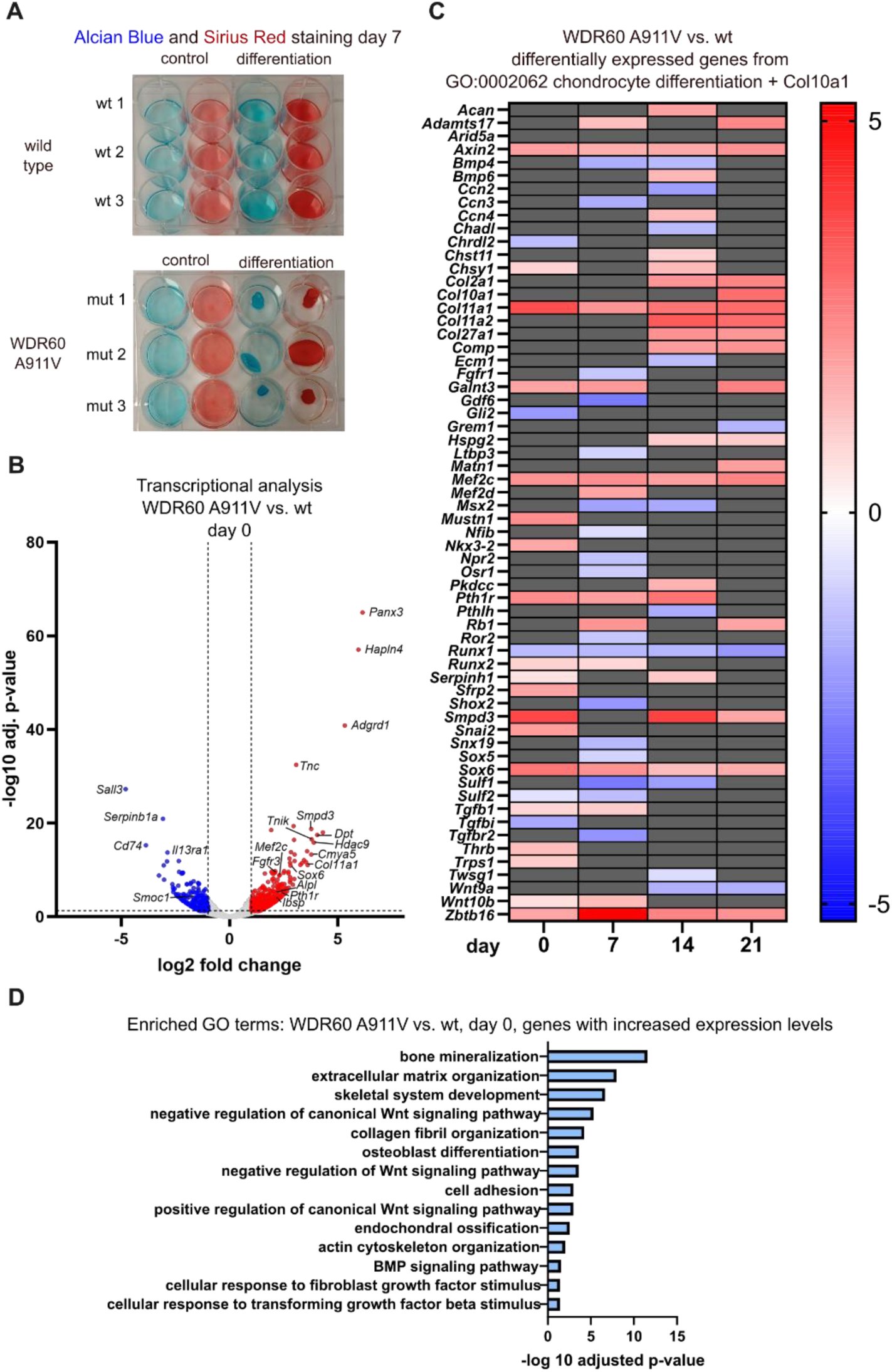
WDR60 A911V mutants exhibit early transcriptional changes in chondrocyte differentiation genes. (A) Representative photographs of each three different ATDC5 wild type and WDR60 A911V mutant clones at day 7 under control and differentiation condition after Alcian Blue and Sirius Red staining. (B) Volcano plot illustrating differential gene expression in WDR60 A911V mutants vs. wt at day 0. Genes were color-coded based on log2 fold change thresholds of ±1 and an adjusted p-value ≤ 0.05. (C) Heatmap of significantly differentially expressed genes associated with GO:0002062 (chondrocyte differentiation) and Col10a1 in WDR60 A911V mutants compared to wild type. A high number of differentially expressed genes is observed, indicating altered regulation of chondrocyte differentiation pathways. (D) GO enrichment analysis of genes with increased expression in WDR60 A911V mutants compared to wild type at day 0. Enriched terms include processes related to bone mineralization and skeletal development, indicating early transcriptional changes prior to the onset of differentiation. See also Figure S17 and Suppl. File 6.

GO analysis further revealed enhanced expression of genes associated with canonical Wnt signalling in IFT74 del ex 2 and WDR60 A911V compared to controls (Figure 7). Wnt signaling plays a key role in skeletal development, with β-catenin-dependent (canonical) and independent (non-canonical) pathways exerting opposing effects on chondrogenesis: canonical signalling promotes chondrocyte maturation and hypertrophy^30,31^, whereas non-canonical ligands like WNT5A inhibit hypertrophy^31,71^. We found increased gene expression of the canonical Wnt activator TNIK^82^, Wnt positive regulator BAMBI^83^ and negative feedback regulators AXIN2 and WIF1^84,85^ in WDR60 A911V and in IFT74 del ex 2 mutants at day 14 compared to controls (Figure 7).

**Figure 7:**
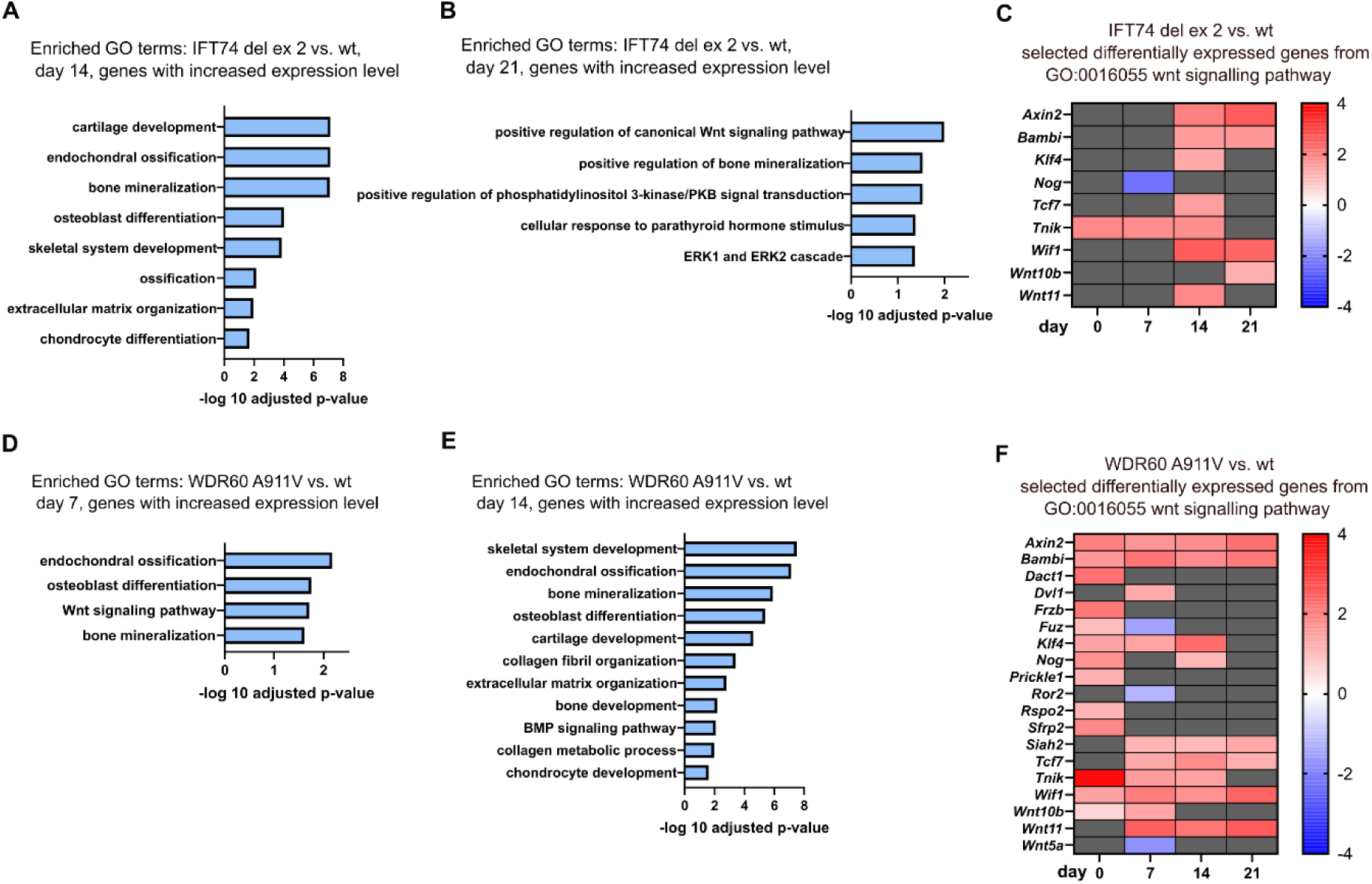
Concurrent canonical Wnt signalling and late skeletal differentiation signatures in WDR60 A911V and IFT74 del ex 2 mutants. (A, B) GO enrichment analysis of genes with increased expression in IFT74 del ex 2 mutants compared to wild type controls at day 14 (A) and day 21 (B). Enriched terms include processes related to positive regulation of canonical Wnt signalling, ossification and bone mineralization. (C) Selected differentially expressed genes associated with GO:0016055 (Wnt signalling pathway) in IFT74 del ex 2 mutant cells compared to wild type. (D, E) GO enrichment analysis of genes with increased expression in WDR60 A911V mutant cells compared to wild type controls at day 7 (D) and day 14 (E). Enriched terms include Wnt signalling as well as processes related to ossification and bone mineralization. (F) Selected differentially expressed genes associated with GO:0016055 (Wnt signalling pathway) in WDR60 A911V mutant cells compared to wild type. Details on GO analysis are provided in Suppl. File 6.

In addition, WDR60 A911V and IFT74 del ex 2 mutant cells also shared reduced expression of several ECM proteins (Figure S19), including microfibril components FBN1, FBN2, EMILIN1 and LTBP1, which also regulate TGF-β signalling^86–89^, as well as *Ccn2* encoding connective tissue growth factor (CTGF). Interestingly, knockout mouse models for *Ccn2* exhibit growth plate abnormalities and impaired ossification manifesting as a developmental chondrodysplasia phenotype^90^. While we overall observed limited overlap between transcriptomic and proteomic findings in mutants compared to controls, PHOSPHO1, a key regulator of bone mineralization^76,91^, showed increased expression at both gene and protein levels in WDR60 A911V and IFT74 del ex 2 mutants at day 14 (Figure S19). Differential gene expression and ECM proteomic analyses for timepoints other than day 14 are shown in Suppl. File 4 and 5.

Altogether, this suggests disturbances of chondrogenic differentiation, altered Wnt signalling and changed composition of the secreted ECM in IFT74 del ex 2 and WDR60 A911V hypomorphic mutants.

### IFT74 del ex 2 mutants show early ECM alterations followed by increased late-stage chondrogenic differentiation marker gene expression

While dynein-2, IFT-A, and IFT-B SRTD patients all exhibit shortened ribs and long bones, phenotype severity differs, with dynein-2 patients showing higher cardiorespiratory mortality^22^. Surviving dynein-2 and IFT patients appear to “grow out” of the skeletal phenotype, with reduced bone shortening over time^22^, IFT74 del ex 2 deletion patients in contrast present with worsening growth failure over time^20^. Together, this may point towards protein-specific functions of individual dynein-2 and IFT complex components. To dissect those in the context of chondrogenic differentiation and ECM composition, we next analysed mutants separately. Detailed mutant specific differential gene or protein expression including volcano plots, overlap analysis and GO analysis are provided in Figures 4, S7, S15-S18, S20-S23 and Suppl. Files 1, 6 and 7.

Overall, IFT74 del ex 2 mutants displayed a marked increase in differentially expressed genes compared to controls associated with the GO term “chondrocyte development*”* (GO:0002062) from day 14 onwards, whereas alterations in the ECM composition compared to controls were already apparent at day 7 (Figure 5 D, Figure S22 A, C, E). Notably, at day 14, early chondrocyte differentiation marker *Col2a1*^92^ and hypertrophy marker *Col10a1*^93^, as well as *Col11a1* and *Col11a2*, were among the most significantly differentially expressed genes (Figure 5 C). Differentially expressed genes for all time points are displayed in Figure S16. ECM proteomics analyses did not reveal many proteins consistently changed in IFT74 del ex 2 mutants versus controls at all timepoints. Instead, distinct ECM proteins were differentially expressed at each time point (Figure 4, Figure S22).

Notably, at day 7 the ECM of IFT74 del ex 2 mutants was deficient for several ECM components compared to control, involved in regulation of ECM architecture including TGM2 and POSTN^94–96^ as well as proteins regulating TGF-beta availability^97^ (e.g. FBN2, TGFB2, SMOC1, details in Suppl file 4 and 7)^87,98,99^. These remained reduced at day 14, while simultaneously higher expression of GPC1 (Glypican-1) and MFGE8, both glycoproteins that have been shown to modulate various signalling pathways like canonical Wnt and TGF-beta^100,101^, and PHOSPHO1 was observed. By day 21, few proteins showed reduced expression, whereas COL2A1, COL10A1, COL11A1, and COL11A2 were elevated, consistent with transcriptomic changes observed at day 14 (Figure 5 C, D).

Similarly, only a few genes showed consistent changes across multiple time points (Figure S20). Among the four genes with consistent lower expression levels in mutant versus wildtypes (*Sorl1*, *Nos1*, *Ifih1*, and *Arhgef3*), two have known links to skeletal biology but opposing effects on bone mass/density: *Nos1* (neuronal nitric oxide synthase) is a positive regulator of bone mass^102^, probably due to disturbed osteoclast/osteoblast balance^103^, while *Arhgef3*, a guanine nucleotide exchange factor for RhoA and RhoB, was associated with osteoporosis in humans^104,105^, although *Arghef3* mutant mice did not display remarkable skeletal phenotypes^106^.

### Increased early expression of late skeletal development marker genes and persistent ECM alterations in WDR60 A911V mutants

We next dissected effects of hypomorphic WDR60 loss of function in the context of the severe skeletal phenotype observed in human WDR60 related SRTD and Jeune asphyxiating thoracic dysplasia (JATD) cases^21,107^. In line with more pronounced early-age bone shortening in dynein-2 compared to IFT patients, in WDR60 A911V mutant clones, we observed more pronounced transcriptional and ECM proteomic alterations compared to wild type than for IFT74 del ex 2 or IFT43 M1V clones, accompanied by detachment of mutant cells from the cell culture well surface at around day 7 of differentiation, which was not observed for other mutants or wild type clones (Figure 6 A, Figure S11). Transcriptional analysis revealed that many genes associated with the GO term “chondrocyte differentiation” (GO:0002062) were differentially expressed compared to wild type at all time points, including lower and higher expressed gene subsets (Figure 6 C). Genes with consistently elevated expression included key chondrocyte transcription factors *Sox6*^108^ and *Mef2c*^74^ as well as late stage regulators *Smpd3* and *Zbtb16*^109,110^. Genes with reduced expression at ≥ 3 time points were mostly stress response genes, whereas those with increased expression at ≥ 3 time points were enriched for factors involved in bone mineralization, ossification, and Wnt signalling (Figure S24, Table S5 + S6).

Notably, genes with increased expression in WDR60 A911V mutants were enriched for GO terms related to bone mineralization and Wnt signalling regulation already at day 0 (Figure 6). From day 0 onward, these mutants showed significant induction of key osteogenic markers *Alpl* and *Ibsp*^75,111^ (Figure 6 B, S17, S21).

The extensive transcriptional changes in WDR60 A611V mutants versus controls were accompanied by stable ECM composition alterations with numerous proteins showing consistently increased or reduced expression (Figure 4 D, S22 B, D, F). Proteins with reduced expression include regulators of TGF-beta/BMP signalling pathways and their availability in the ECM (ASPN, BMPER, TGFB2, LTBP1^112–114^), structural components (COL12A1, FMOD, OGN, POSTN), ECM enzymes (ADAMTS1, HTRA1, LOX, TIMP3) as well as SMOC1 and SMOC2, known regulators of bone formation ^115^. For detailed gene/protein lists see Supplement Files 1, 4, 5 and 7.

### IFT43 M1V mutants show minimal changes in transcriptomic and proteomic analyses but reduced SOX9 expression on day 0

Across all time points, IFT43 M1V mutants displayed few genes with altered expression, with the largest number at day 0 (Figure S15, S7 C, D). Although not statistically significant in transcriptomics (p = 0.07, log2fc = -0.94 at day 0), Sox9 was selected for validation due to its key role in chondrocyte differentiation^116,117^ (Figure 5 E). Western blot confirmed lower SOX9 protein levels in mutants at day 0 (Figure 5 G, H), with levels comparable to wild type at later stages. In line with reduced SOX9 expression *Dmrt2*, a Sox9-inducible transcription factor in pre-hypertrophic chondrocytes^118^ was also reduced at two time points, similar to findings in WDR60 A911V mutants (Figure S21).

*Pcdh7* (Protocadherin 7) was the only gene with significant consistent lower expression in *Ift43* mutant versus control cells across all timepoints (Figure S7 D). *Pcdh7* knockout mouse studies demonstrated its role in regulation of osteoclast differentiation via RhoA and Rac1^119^. Apart from *Tgfbi* and *Smpd3*, reduced at day 0 and 21 respectively, no other chondrocyte differentiation genes (GO:0002062) were differentially expressed (Figure 5 F).

ECM proteomics analyses revealed reduced expression of bone development regulators IGF2, IGFBP7 and osteoprotegerin (TNFRSF11B)^65,120,121^ was observed in later stages of differentiation (Figure 4 C). Additional details of proteomic analyses are displayed in Figure S23.

### Complete loss of WDR60 causes only minor transcriptional and proteomic alterations during the ATDC5 differentiation process

Dynein-2, the motor complex for retrograde IFT^11^, is essential for vertebrate development. Loss of dynein-2 components has been shown to decrease dynein-2 complex stability^18,122^, to cause severely shortened cilia^16,38^ (Figure 2) and results in mid-gestational lethality in mice^38,123^. Consistent with this, with the exception of the dynein-2 light chain TCTEX1D2^122^, no human individuals harbouring biallelic complete loss-of-function alleles have been reported, likely due to embryonic lethality. Hence, dynein-2 or IFT knockout models do not reflect the human disease state, but are of interest to investigate the effect of cilia loss on cell differentiation and ECM secretion. We therefore investigated the impact of biallelic *WDR60* loss of function in ATDC5 cartilage-like organoids. As expected from nonsense-mediated decay of the out-of-frame indels, WDR60 (*Dync2i1*) expression was significantly reduced (Figure S25), though differences were no longer significant by day 21, likely due to read misalignment (Figure S26). Surprisingly, despite lack of cilia in WDR60 nonsense cells in these mutants (Figure 2 A, E), transcriptomic and ECM proteomic changes were modest. Few genes annotated to GO:0002062 (chondrocyte differentiation) showed transient changes, but none were consistently altered across time points (Figure 5 B).

Similar to the surprisingly little affected gene expression, ECM proteome analyses revealed consistently reduced expression of only two proteins, serine protease HTRA1 and glycoprotein Tenascin-W (TNN), throughout differentiation. Aside from this, differential protein expression was limited to a few individual proteins at each time point (Figure 4 A, S23 B, D, F).

## Discussion

Hypomorphic human disease alleles for IFT-A, IFT-B and dynein-2 components cause qualitatively similar chondrodysplasia phenotypes with shortening of long bones and ribs, but with genotype depending severity. Phenotypic severity is greatest for dynein-2 dysfunction, while IFT-A patients often present with additional renal, retinal or craniofacial features^22^. Bone shortening generally becomes less pronounced with age in dynein-2 and IFT patients^22^, except for IFT74 patients, where growth deficit increases over time^20^. This indicates generally shared mechanisms underlying the skeletal phenotype resulting from IFT-A, IFT-B and dynein-2 dysfunction but also individual protein functions. Biallelic loss-of-function of any IFT-A, IFT-B, or dynein-2 component severely disrupts cilia, so null models are unsuitable for studying human disease alleles or individual protein functions. We therefore generated ATDC5 models carrying hypomorphic alleles for IFT-A, IFT-B, and dynein-2 components, alongside a WDR60 complete loss-of-function model.

### Effects of IFT74, IFT43 and WDR60 dysfunction on ciliogenesis, IFT and Hedgehog signalling

Effects of hypomorphic human IFT dynein disease alleles on ciliation efficiency and cilia length have remained unclear and somewhat controversial. Some studies report reduced ciliation and shorter cilia in hypomorphic mouse models or fibroblasts from ciliopathy patients^36,107,124,125^, whereas others indicate minimal effects on cilia length or variable phenotype severity^20,50,54,126^. These discrepancies may reflect comparisons across patients and mouse models where genetic background effects exceed effects of individual IFT or dynein-2 gene mutations. In our study, hypomorphic mutants displayed similar ciliation and cilia length compared to wild type, suggesting that these hypomorphic alleles do not disrupt cilia structure in this cellular context. While several studies report cilia formation in presumed *Wdr60* null cell lines^39–41^, in line with previous studies investigating complete loss of function of other dynein-2 complex components such as DYNC2H1^16^ or WDR34^38^, we did not observe cilia in WDR60 nonsense cells. As previous observations were made in distinct cellular models, cell line specific differences may contribute to the observed discrepancy.

Dynein-2 and IFT-A human disease alleles have been reported to cause IFT protein accumulation at the ciliary tip in patient fibroblasts^18,36,126,127^, underlining their role for retrograde IFT. Consistently, we observed IFT88 accumulation at distal cilia parts for IFT43 M1V cells. In contrast, IFT74 del ex 2 and WDR60 A911V mutants displayed a uniform distributed of IFT88 along the axoneme. For IFT74, this is in line with previous findings in the corresponding mouse model^20^ and the slight shift toward a more uniform IFT88 distribution along the axoneme in these cells may suggest a decrease in IFT velocity. For WDR60 A911V mutants, we would have expected a more pronounced accumulation at the distal ciliary end^18,126^.

In ciliary chondrodysplasia mouse models and patient fibroblasts, disturbed Hedgehog signalling is frequently reported^18,19,128,129^. The decreased Gli3 repressor/Gli3fl ratio together with the lack of responsiveness to SAG in WDR60 nonsense and IFT74 cells are consistent with published observations of Hedgehog disturbances in mouse^38,123^ and cellular models^39,40^ lacking dynein-2 components. Among the few genes differentially expressed across multiple time points in WDR60 nonsense cells versus controls, increased *Gli1* expression was the most notable. Our findings align with previous reports about patient fibroblasts with hypomorphic IFT74 mutations^130^ and chondrocytes from patients carrying IFT81 (IFT-B) mutations^55^, but contrast previous reports about reduced Hedgehog signalling and lower Gli1 expression in IFT/dynein-2 knockout mouse models^16,28,38,131^. However the observations are in accordance with results described in the articular cartilage of Col2aCre;Ift88fl/fl mutant mice^129^.

Despite significantly impaired retrograde IFT, unexpectedly no impaired response to SAG stimulation, nor significant *Gli1* expression changes at basal state were found in IFT43 M1V mutants, in contrast to previous studies using hypomorphic mouse models for *Ift144* and *Ift140* (IFT-A) reporting Hedgehog signalling defects and Gli3 misexpression in developing limbs^13,50,132^. Surprisingly, we neither observed an impaired Hedgehog response in WDR60 A911V mutant cells in contrast to previous results from WDR60 mutant patient cells^107^ and experiments from our laboratory in IMCD3 cells carrying the same mutation^21^. This could point towards cell/tissue specific effects. Transcriptomics analyses did not reveal significant expression differences for Hedgehog pathway components or direct target genes at basal state, with the exception of increased Gli1 expression in WDR60 null cells. Overall, the different findings of in vitro and vivo studies may result from presence of Hedgehog ligand in vivo while in our system, we did not use any Hedgehog pathway activation. Future studies could include transcriptional analyses after Hedgehog pathway activation.

### Effects on chondrogenic differentiation and ECM composition

Previous mouse model studies^28,128,133,134^ and studies on growth plates from SRTD patients^17,55,135^ suggest shortening and disorganization of the growth plate, affecting both proliferative and hypertrophic zones as well as loss of the typical columnar organization of chondrocytes. In addition, impaired mineralization and attenuated hypertrophic differentiation have been reported^128,133,136^, while some studies also describe accelerated hypertrophy accompanied by ossification defects^28,134,137^. Cilia act as mechanosensors oriented into the ECM^24^, decorated with integrins^138^ and interact with ECM components^24^. They transduce mechanical cues into developmental signalling responses^23,139^, for example by recruiting Pth1 receptors^140^. Accordingly, ECM alterations may influence ciliary signalling, while ciliary dysfunction may in turn affect ECM composition and structure. Although ECM changes are well described in non-ciliary chondrodysplasias^25,26^, this mechanism remains poorly explored in ciliary chondrodysplasia.

Our data suggests accelerated hypertrophy and premature ossification, reflected by elevated *Col10a1* and enrichment of late chondrogenic, osteogenic, and mineralization markers, including PHOSPHO1 at both transcript and protein levels in WDR60 A911V and IFT74 del ex 2 mutants. Increased *Col2a1* expression at day 14 likely reflects incomplete transition between proliferative and hypertrophic states rather than failure to initiate hypertrophy, consistent with the disorganized zonal architecture described in IFT mutant models^128,133,134^ and patients^17,55,135^.

Consistent with a previous ATDC5 study showing Sox6- and Mef2c-dependent induction of Smpd3^141^. *Mef2c* expression was consistently increased in WDR60 A911V mutants and elevated at later stages in IFT74 del ex 2 mutants. As *Mef2c* overexpression promotes hypertrophy and ossification in vivo^74^ and its expression is reduced upon Wnt inhibition^142,143^, the coincident increase in Wnt signalling and Mef2c may potentially explain the tendency toward accelerated chondrocyte hypertrophy.

WDR60 A911V mutants further did not follow the expected multistep differentiation program, instead showing early expression of late-stage skeletal markers and detachment from the culture surface. However, proliferation rates at early time points and initial collagen or proteoglycan deposition were comparable to controls (Figure S10 E, F, S27), and fibronectin staining did not reveal structural ECM abnormalities (Figure S28).

The overall small number of differentially expressed chondrocyte differentiation markers and ECM proteins, suggests that the IFT43 M1V mutation has only limited influence on the differentiation process and on ECM composition. Overall, these findings align with the comparatively mild skeletal phenotype described in patients carrying IFT43 M1V mutations^36^.

Interestingly, despite loss of cilia and aberrant Hedgehog signalling, WDR60 nonsense mutants showed only minor transcriptional and ECM changes compared to wild type (Figure 5 A, B, Figure S18, S23). This raises the question if the cilium as a structure is required for the ATDC5 cell chondrogenic differentiation program. Tao et al.^144^ previously reported impaired ATDC5 differentiation upon chloral hydrate-mediated cilia removal every second day, in the light of our results, these effects may partly reflect the compound’s toxicity^145^.

Previous studies have shown that mutations in *Ift80* and *Ift88* impair collagen X and proteoglycan expression^125,128^. Further, studies in IFT80 hypomorph mice reported reduced endocytotic uptake of ECM proteases at the ciliary pocket^146^. Here, we provide the first characterization of ciliary chondrodysplasia ECM using the ATDC5 cartilage-like organoids, performing proteomics analyses at several differentiation timepoints and correlating these results with transcriptomics analyses. Overlap between datasets was limited, with PHOSPHO1 being one of the few factors increased at both gene and protein levels in WDR60 A911V and IFT74 del ex 2 mutants. This is consistent with previous chondrocyte multi-omics studies reporting modest concordance^147–149^, and likely reflects post-transcriptional regulation and differences in protein turnover. Overall, reduced expression of ECM-remodeling enzymes and structural components suggests impaired matrix integrity. Notably, ECM alterations in IFT74 del ex 2 mutants preceded corresponding transcriptional changes, indicating early effects on protein secretion or stability rather than transcriptional regulation.

### Effects of hypomorphic disease alleles and WDR60 complete loss of function on Wnt signalling

Wnt signalling plays a crucial role in skeletal development^150^, preventing hypertrophy early in chondrogenesis but driving hypertrophy and ossification in later maturation stages^150–152^. Interestingly, alongside markers of accelerated hypertrophy and ossification we observed increased expression of canonical Wnt signalling components in WDR60 A911V and IFT74 mutants after day 14. Consistent with our observations, Chang and Serra reported accelerated hypertrophy together with elevated canonical Wnt signalling in a Col2aCre;Ift88fl/fl mouse model^131^. Two more studies observed increased canonical Wnt signalling after IFT80 knock-out in a cell model^153^ and in Col2α1; IFT80f/f mutant mice^128^. As WDR60 nonsense mutants showed no Wnt signalling disturbances and cilia may not be required for Wnt signalling^154^, the observed increase in canonical Wnt signalling in WDR60 A911V and IFT74 del ex 2 mutants could be independent of the primary cilium as a structure.

ECM proteomics revealed reduced WNTA5A expression in all hypomorphic mutants at days 7 and 14. Non-canonical Wnt5a signalling, acting in part via the PCP pathway, regulates cytoskeletal organization and cell polarity^155^, including in chondrocytes^68,69,72^. Reduced Wnt5a expression may further contribute to accelerated hypertrophy in WDR60 A911V and IFT74 del ex 2 mutants, as Wnt5a restrains hypertrophic differentiation^71^ and inhibits canonical Wnt^156^. The discrepancy between reduced WNT5A protein and unchanged mRNA levels suggests post-transcriptional regulation, such as altered secretion or protein stability.

## Conclusion

Overall, our results indicate that in our ATDC5 in-vitro model of chondrogenesis, WDR60 A911V and IFT74 del ex 2 disease alleles accelerated chondrocyte hypertrophy, coinciding with Wnt signalling changes, whereas the IFT43 M1V mutation had only little impact. Importantly, we showed for the first time that hypomorphic mutations in IFT/dynein-2 genes alter ECM composition during chondrogenesis. In this context, reduced expression of WNT5A in hypomorphic IFT and dynein-2 mutants highlights Wnt5a and the PCP pathway as candidates for follow- up studies. Furthermore, our data suggest that primary cilium function is dispensable for ATDC5 differentiation per se, which proceeds despite impaired Hedgehog signalling. Nonetheless, the pronounced effects of hypomorphic disease alleles on specific aspects of differentiation indicate that affected proteins have at least a modulatory role for chondrocyte differentiation.

## Methods

Additional methodological details are provided in the supplemental data.

### Cell culture

ATDC5 cells were purchased from ECACC. Cells were maintained Dulbecco’s modified eagle medium (DMEM, 32430100, Gibco) and Ham’s F-12 (11765054, Gibco) in a ratio of 1:1, supplemented with 10 % fetal calf serum (FCS, F7524, Sigma), 1% Sodium Pyruvate (S8636, Sigma), 100 U/ml penicillin and 100 µg/ml streptomycin (P-0781, Sigma) at 37°C and 5% CO_2_. In differentiation medium FCS was reduced to 5% and it was supplemented with 10µg/ml insulin (I9278, Sigma), 30nM sodium selenite (S5261, Sigma), 50µg/ml ascorbic acid (A4403, Sigma) and 10µg/ml transferrin (90190, Sigma).

### Generation of CRISPR/Cas9 mutant ATDC5 cell lines

Mutant lines were created using CRISPR or CRISPR base editing to introduce point mutations (WDR60 A911V and IFT43 M1V) or frameshift mutations (IFT74 del ex 2 and WDR60 nonsense). Correct editing was confirmed by sequencing. Detailed protocols are provided in the Supplementary Methods.

### Immunofluorescence

Primary cilia were visualized by immunofluorescence staining of acetylated tubulin, gamma tubulin and IFT88. Cilia length, ciliation efficiency and IFT88 accumulation were quantified from fluorescent images using ImageJ.

### Proteoglycan and collagen staining

Cells were fixed with methanol and stained with 0.1% Alcian Blue (A5268, Sigma) in 0.1M HCl or Sirius Red solution (13422, Morphisto), for 2 hours at room temperature. Following incubation, excess dye was removed and images were taken.

### Western blot analysis

Cells were lysed in RIPA buffer supplemented with protease inhibitors. Protein concentration was determined by Bradford assay before SDS-PAGE. Nitrocellulose membranes were probed with primary and secondary antibodies as detailed in the Supplementary Methods, and signals were detected by enhanced chemiluminescence. Band intensities were quantified by densitometry using ImageJ.

### SAG treatment and Hedgehog pathway analysis

Confluent cells were kept under low serum conditions (0.5% FCS) and treated with Smoothened agonist SAG for 24h to stimulate Hedgehog signalling, followed by western blot analysis.

### Transcriptomic analysis

Total RNA was isolated from ATDC5 cells and library preparation and sequencing performed by Novogene (Cambridge, UK). Reads were processed and aligned to the mouse genome (mm39), and gene expression was quantified using Galaxy platform (https://usegalaxy.org). Differential expression analysis was conducted using DESeq2.

### ECM Proteomic analysis

After decellularisation, lysis of remaining ECM and preparation for mass spectrometry analysis was performed as described previously^157^. Peptides were analysed by LC-MS/MS, and protein identification and quantification were performed using standard pipelines.

### Statistical analysis

GraphPad Prism 10 software was used for data visualization and statistical analysis. Data are presented as mean ± SD. Statistical analyses were performed as indicated in the figure legends. Two-group comparisons were conducted using unpaired two-tailed Student’s t-test. Two-way ANOVA followed by Sidak’s or Tukeýs multiple comparisons test was used to perform pairwise comparisons of each mutant to the wild-type control, as indicated in the figure legends. For transcriptomic and proteomic data, p-values were adjusted for multiple testing using the Benjamini–Hochberg method, and adjusted p-values (FDR) ≤ 0.05 were considered significant (*p ≤ 0.05, **p ≤ 0.01, ***p ≤ 0.001).

## Data availability statement

All RNA sequencing data generated in this study have been deposited in the Gene Expression Omnibus under accession number GSE326420. Mass spectrometry raw data have been deposited at the ProteomeXchange Consortium (http://proteomecentral.proteomexchange.org) under the accession number PXD076232. Furthermore, all mass spectrometry proteomics datasets used and/or analysed during this study are available online at the MassIVE repository (http://massive.ucsd.edu/; dataset identifier: MSV000101257)

## Acknowledgement

We thank Christoph Schell and Maximilian Wess for helpful discussion of fibronectin immunofluorescence. The Proteomic Platform – Core Facility (ProtCF) was supported by the Medical Faculty of the University of Freiburg to Prof. Dr. Oliver Schilling (2021/A3-Sch and 2023/A3-Sch). We acknowledge the support of the Freiburg Galaxy Team, funded by the German Federal Ministry of Education and Research <u>BMFTR</u> grant 031 A538A <u>de.NBI</u>-RBC and the Ministry of Science, Research and the Arts Baden-Württemberg (MWK) within the framework of LIBIS/de.NBI Freiburg. MS acknowledges funding from the Deutsche Forschungsgemeinschaft (DFG) Cilia FOR 5547 Cilia Dynamics (project 503306912) and CRC 1597 SmallData-project number 499552394. MS, AK and SA acknowledge funding from the Deutsche Forschungsgemeinschaft (DFG), Cluster of Excellence, CIBBS – Centre for Integrative Biological Signalling Studies (EXC-2189, project 390939984) and MS, AK and OS acknowledge funding from the Deutsche Forschungsgemeinschaft (DFG), SFB1543 NEPHGEN (project 431984000).

## CRediT authorship contribution statement

**Anna Klawonn:** Writing – review & editing, Writing – original draft, Investigation, Formal analysis, Visualization. **Himanshu:** Investigation, Formal analysis. **Zeineb Bakey:** Investigation. **Stefan Tholen:** Formal analysis, Investigation. **Ilona Skatulla:** Investigation. **Yong Li:** Formal analysis, Visualization. **Chiara M. Schröder:** Formal analysis. **Sebastian J. Arnold:** Resources, Funding acquisition. **Anna Köttgen:** Resources, Funding Acquisition. **Oliver Schilling:** Resources, Funding acquisition. **Miriam Schmidts:** Writing – review & editing, Writing – original draft, Resources, Project administration, Funding acquisition, Conceptualization. All authors read and approved of the current manuscript.

