## Supplementary material information for "Distinct effects of hypomorphic IFT and dynein-2 skeletal ciliopathy disease alleles on chondrogenic differentiation, ECM composition and wnt signalling in ATDC5 derived cartilage-like organoids"

Supplementary Files

Supplementary File 1: Comparison of genes with higher or lower expression levels for each mutant vs. wild type at all time points. Gene lists of Venn Diagrams displayed in Figures S18-S20.

Supplementary File 2: Comparison of genes with higher or lower expression levels for each mutant vs. wild types at matched time points. Gene lists of intersections of Venn Diagrams displayed in Figure S8.

Supplementary File 3: Comparison of proteins with higher or lower expression levels for each mutant vs. wild types at matched time points. Protein lists of intersections of Venn Diagrams displayed in Figure S9.

Supplementary File 4: Differentially expressed proteins in ECM Proteomic analyses for each mutant vs. wild type at different time points

Supplementary File 5: Differentially expressed genes in transcriptomic analyses for each mutant vs. wild type at different time points

Supplementary File 6: GO analyses for each mutant vs. wild type at each time point. Genes with increased and decreased expression levels compared to wild type were analysed separately.

Supplementary File 7: Comparison of proteins with higher or lower expression levels for each mutant vs. wild type at all time points. Gene lists of Venn Diagrams displayed in Figure 4.
