## supplementary figures for "Distinct effects of hypomorphic IFT and dynein-2 skeletal ciliopathy disease alleles on chondrogenic differentiation, ECM composition and wnt signalling in ATDC5 derived cartilage-like organoids"

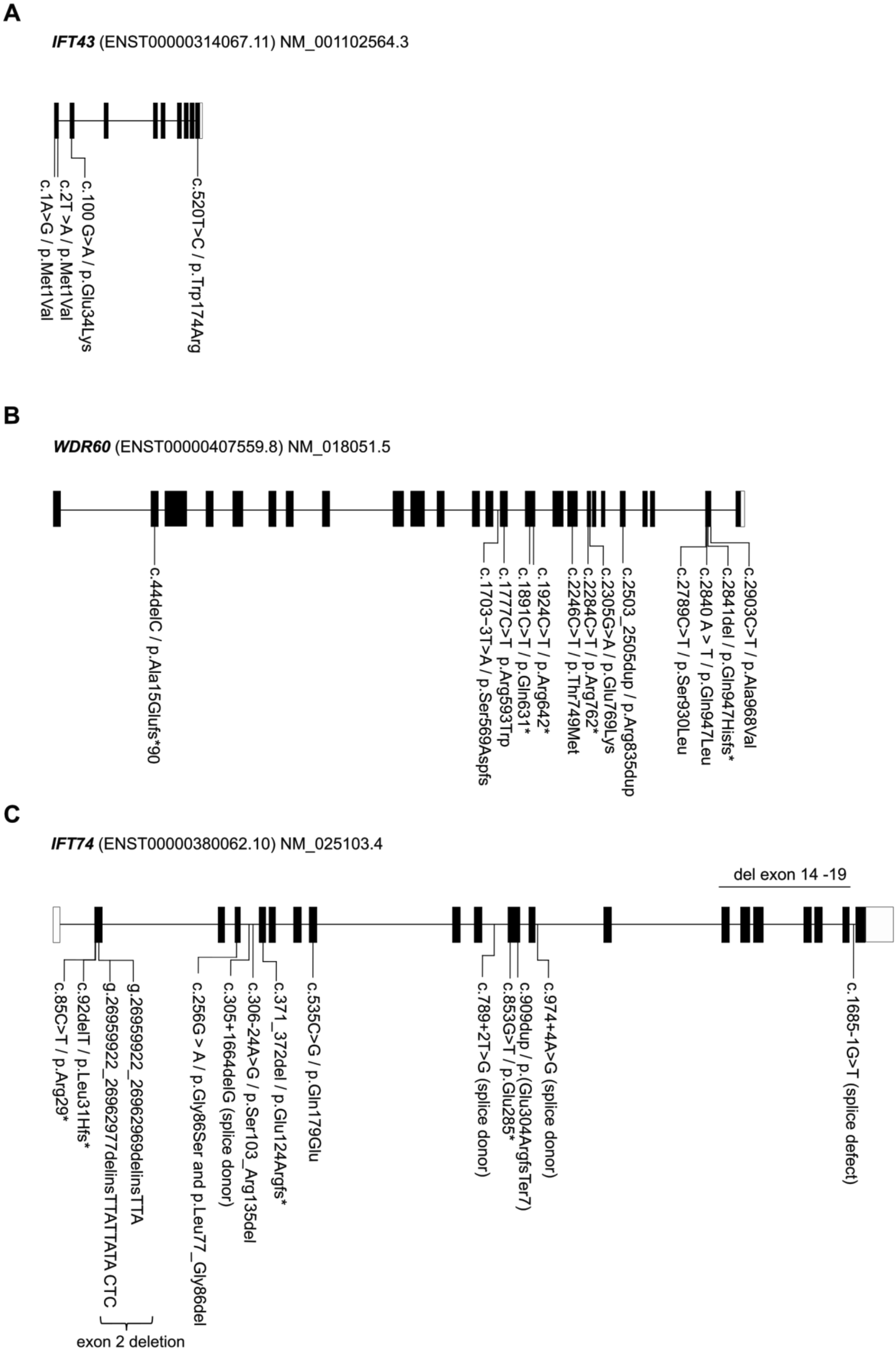
A
IFT43 (ENST00000314067.11) NM_001102564.3
c.1A>G / p.Met1Val
c.2T >A / p.Met1Val
c.100 G>A / p.Glu34Lys
c.520T>C / p.Trp174Arg
B
WDR60 (ENST00000407559.8) NM_018051.5
c.44delC / p.Ala15Glufs*90
c.1703-3T>A / p.Ser569Aspfs
c.1777C>T p.Arg593Trp
c.1891C>T / p.Gln631*
c.1924C>T / p.Arg642*
c.2246C>T / p.Thr749Met
c.2284C>T / p.Arg762*
c.2305G>A / p.Glu769Lys
c.2503_2505dup / p.Arg835dup
c.2789C>T / p.Ser930Leu
c.2840 A > T / p.Gln947Leu
c.2841del / p.Gln947Hisfs*
c.2903C>T / p.Ala968Val
C
IFT74 (ENST00000380062.10) NM_025103.4
del exon 14 -19
c.85C>T / p.Arg29*
c.92delT / p.Leu31Hfs*
g.26959922_26962977delinsTTATTATA CTC
g.26959922_26962969delinsTTA
exon 2 deletion
c.256G > A / p.Gly86Ser and p.Leu77_Gly86del
c.305+1664delG (splice donor)
c.306-24A>G / p.Ser103_Arg135del
c.371_372del / p.Glu124Argfs*
c.535C>G / p.Gln179Glu
c.789+2T>G (splice donor)
c.853G>T / p.Glu285*
c.909dup / p.(Glu304ArgfsTer7)
c.974+4A>G (splice donor)
c.1685-1G>T (splice defect)

Supplementary Figure 1:

Diagram of (A) *IFT43*, (B) *WDR60* and (C) *IFT74* gene structure. Black boxes denote coding exons; white boxes denote 5’and 3’ non-coding exons. Introns are shown as horizontal lines. Disease-associated variants reported in PubMed are highlighted. References are listed in Table S1.

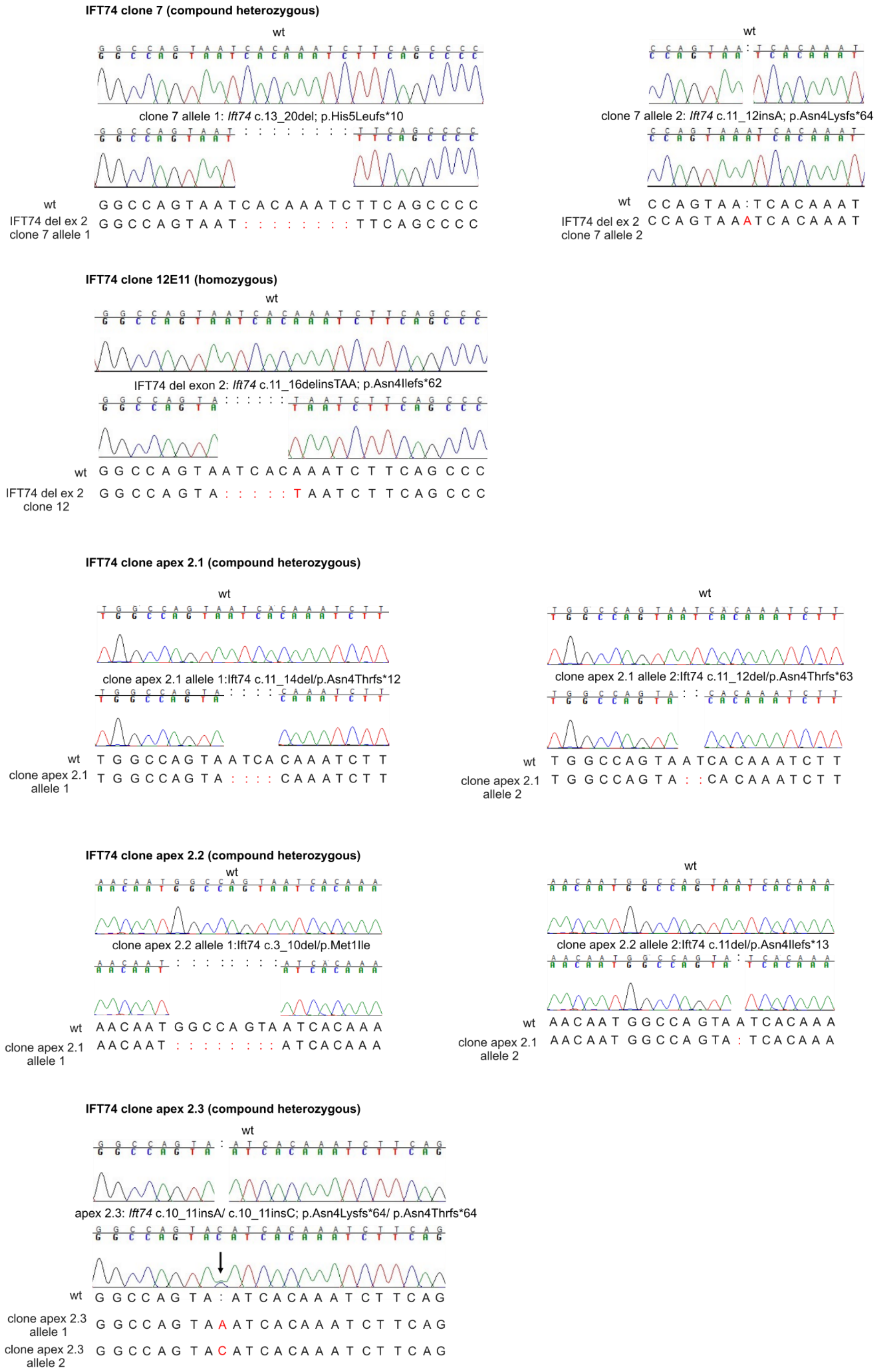
IFT74 clone 7 (compound heterozygous)
wt
clone 7 allele 1: Ift74 c.13_20del; p.His5Leufs*10
wt
IFT74 del ex 2
clone 7 allele 1
G G C C A G T A A T C A C A A A T C T T C A G C C C C
G G C C A G T A A T : : : : : : : : T T C A G C C C C
wt
clone 7 allele 2: Ift74 c.11_12insA; p.Asn4Lysfs*64
wt
IFT74 del ex 2
clone 7 allele 2
C C A G T A A : T C A C A A A T
C C A G T A A A T C A C A A A T
IFT74 clone 12E11 (homozygous)
wt
IFT74 del exon 2: Ift74 c.11_16delinsTAA; p.Asn4Ilefs*62
wt
IFT74 del ex 2
clone 12
G G C C A G T A A T C A C A A A T C T T C A G C C C
G G C C A G T A : : : : : T A A T C T T C A G C C C
IFT74 clone apex 2.1 (compound heterozygous)
wt
clone apex 2.1 allele 1:Ift74 c.11_14del/p.Asn4Thrfs*12
wt
clone apex 2.1
allele 1
T G G C C A G T A A T C A C A A A T C T T
T G G C C A G T A : : : : C A A A T C T T
wt
clone apex 2.1 allele 2:Ift74 c.11_12del/p.Asn4Thrfs*63
wt
clone apex 2.1
allele 2
T G G C C A G T A A T C A C A A A T C T T
T G G C C A G T A : : C A C A A A T C T T
IFT74 clone apex 2.2 (compound heterozygous)
wt
clone apex 2.2 allele 1:Ift74 c.3_10del/p.Met1Ile
wt
clone apex 2.1
allele 1
A A C A A T G G C C A G T A A T C A C A A A
A A C A A T : : : : : : : : A T C A C A A A
wt
clone apex 2.2 allele 2:Ift74 c.11del/p.Asn4Ilefs*13
wt
clone apex 2.1
allele 2
A A C A A T G G C C A G T A A T C A C A A A
A A C A A T G G C C A G T A : T C A C A A A
IFT74 clone apex 2.3 (compound heterozygous)
wt
apex 2.3: Ift74 c.10_11insA/ c.10_11insC; p.Asn4Lysfs*64/ p.Asn4Thrfs*64
wt
clone apex 2.3
allele 1
clone apex 2.3
allele 2
G G C C A G T A : A T C A C A A A T C T T C A G
G G C C A G T A A A T C A C A A A T C T T C A G
G G C C A G T A C A T C A C A A A T C T T C A G

Supplementary Figure 2:

Representative Sanger sequencing chromatograms and description of IFT74 del ex 2 mutant clones. Mutations were confirmed by Sanger sequencing. Heterozygous and homozygous genotypes are indicated, with one or two mutated alleles shown, respectively. Red text indicates sequence changes relative to the wild-type allele.

**WDR60 clone 14 (homozygous)**

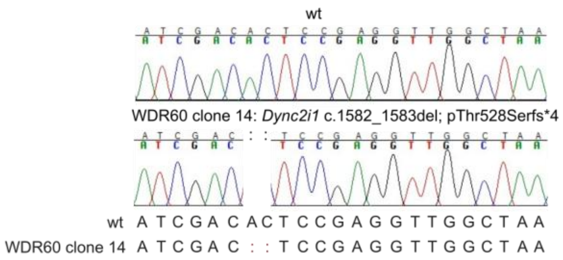


**WDR60 clone 20 (homozygous)**

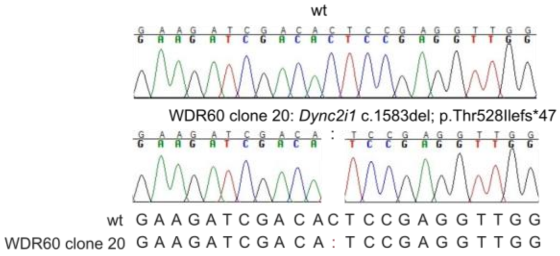


**WDR60 clone 7 (homozygous)**

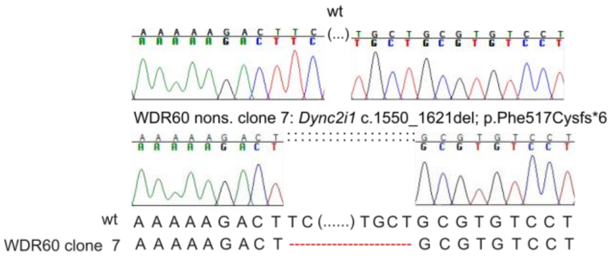


Supplementary Figure 3:

Representative Sanger sequencing chromatograms and description of WDR60 nonsense clones. Mutations were confirmed by Sanger sequencing. Red text indicates sequence changes relative to the wild-type allele.

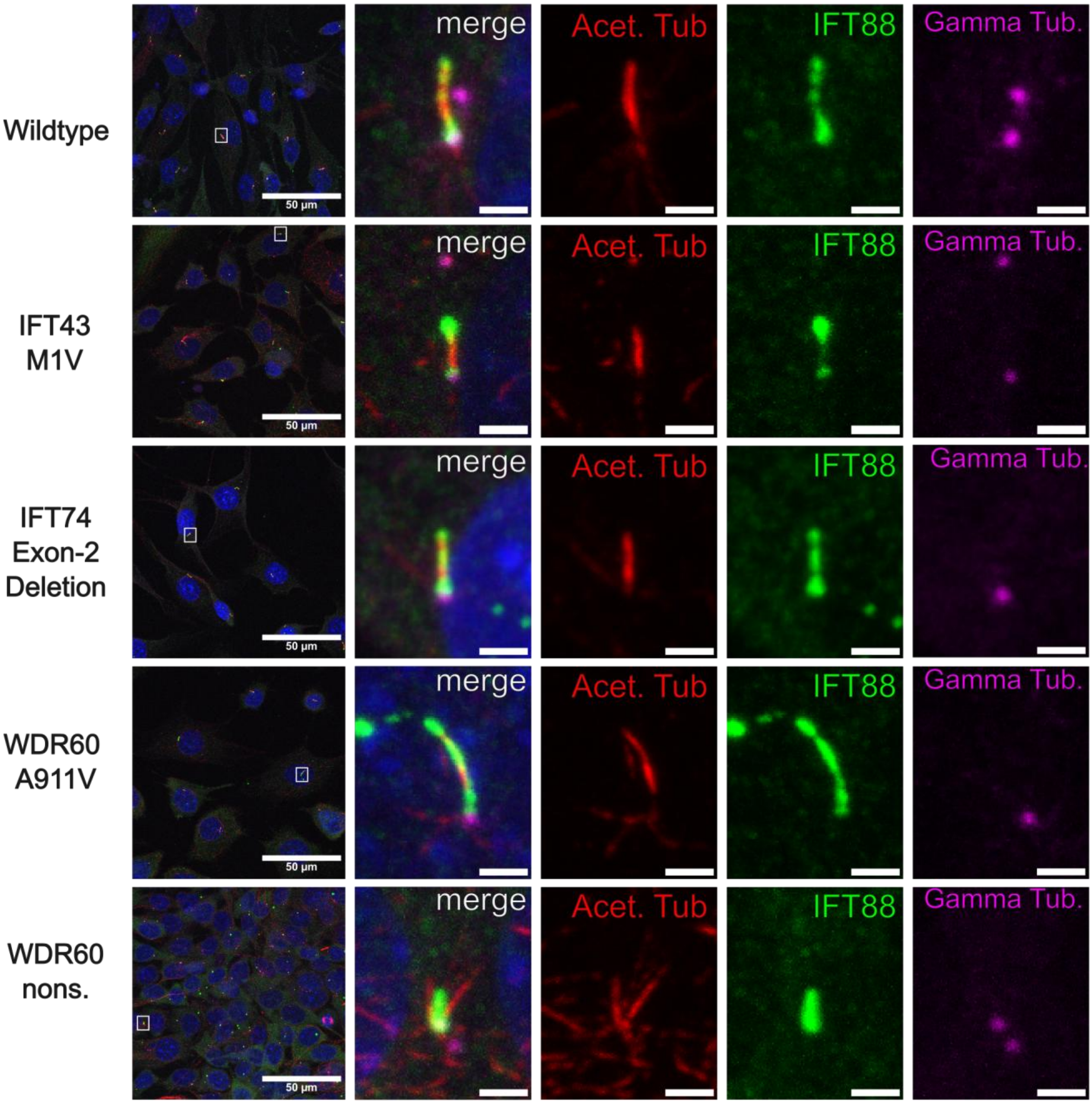


Supplementary Figure 4:

Immunofluorescence analysis of wild-type, IFT43 M1V, IFT74 del ex 2, WDR60 A911V, and WDR60 nonsense clones. Images were taken with Zeiss LSM 880 Observer confocal microscope. Representative overview images, individual channels for acetylated tubulin, IFT88, and gamma tubulin, and corresponding merged images with DAPI nuclear staining are shown. Scale bars 50 µm (left) and 2 µm.

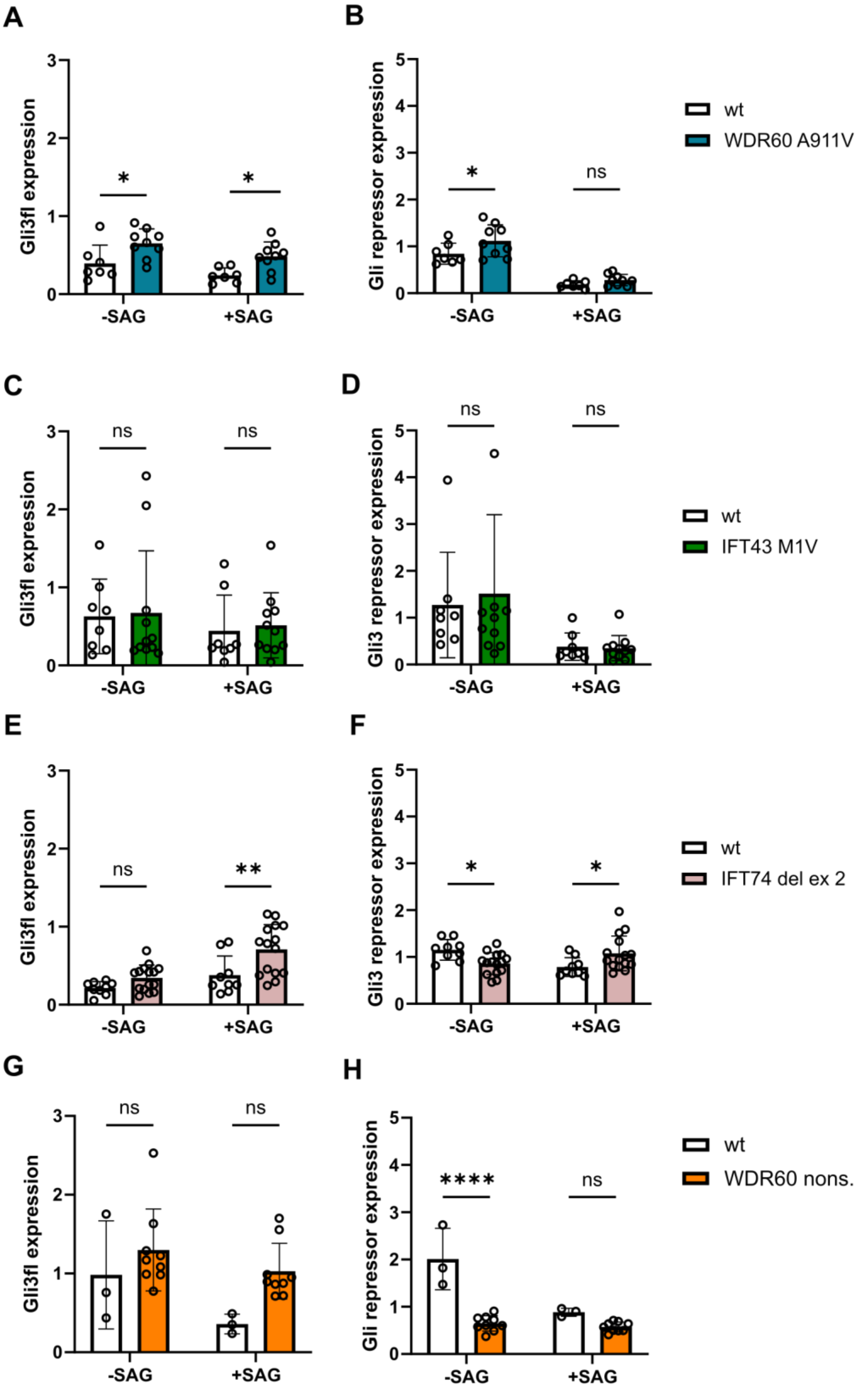


Supplementary Figure 5:

Densiometric quantification of Gli3 full length (A, C, E, G) and Gli3 repressor (B, D, F, H) expression, normalized to beta actin (loading control) for WDR60 A911V, IFT43 M1V, IFT74 del ex 2 and WDR60 nonsense mutants before and after SAG treatment. Expression was calculated for each sample and pooled across biological replicates containing multiple wild-type and mutant samples per experiment. Bars represent mean ± SD from at least three independent experiments. Statistical significance was determined using two-way ANOVA with Sidak´s multiple comparisons test to compare mutant to wild type samples (n= 3-15, *p < 0.05, ** p < 0.01, *** p < 0.001, **** p < 0.0001)

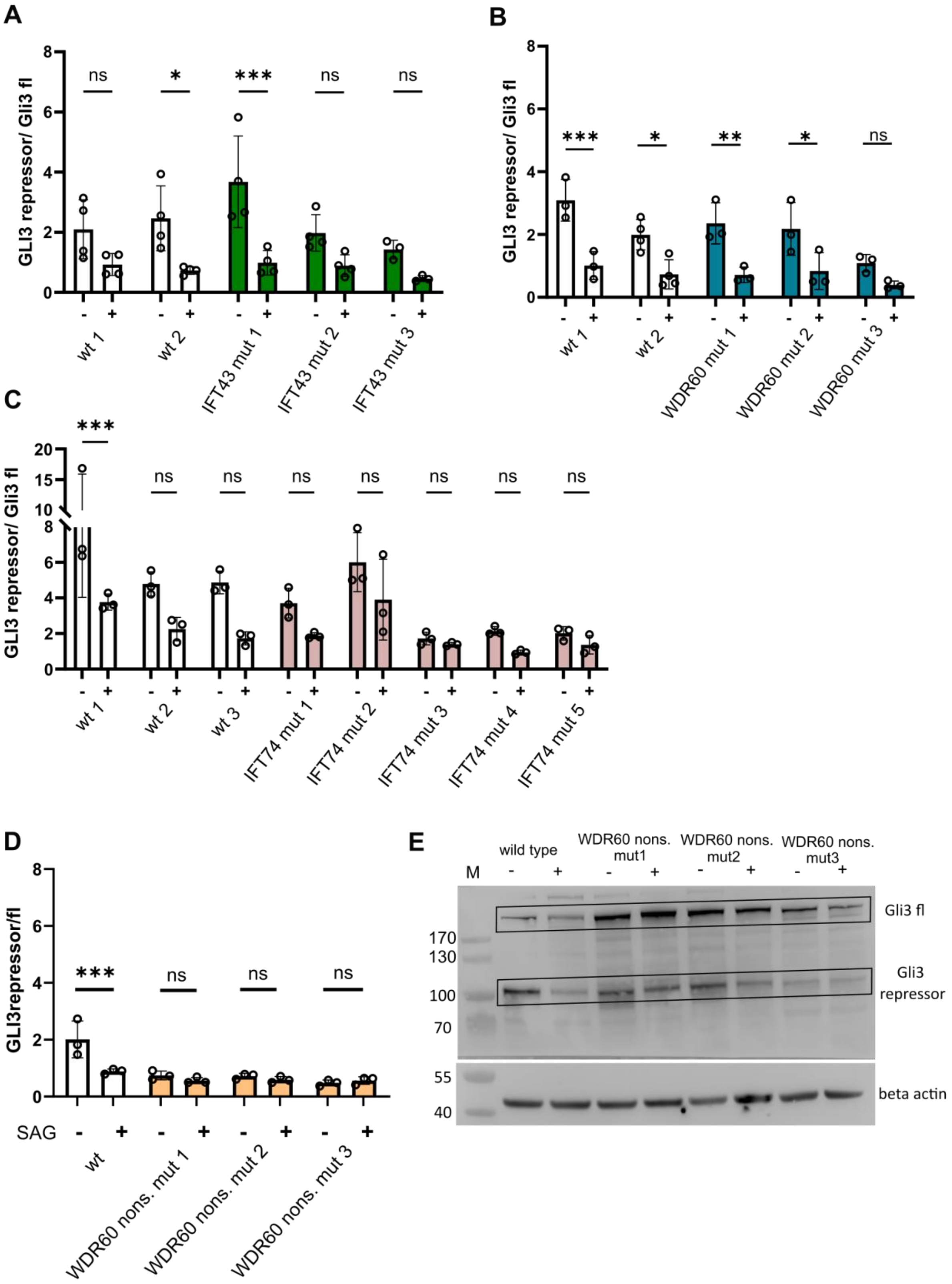

A
GLI3 repressor/ Gli3 fl
ns
*
***
ns
ns
wt 1
wt 2
IFT43 mut 1
IFT43 mut 2
IFT43 mut 3
B
GLI3 repressor/ Gli3 fl
***
*
**
*
ns
wt 1
wt 2
WDR60 mut 1
WDR60 mut 2
WDR60 mut 3
C
GLI3 repressor/ Gli3 fl
***
ns
wt 1
wt 2
wt 3
IFT74 mut 1
IFT74 mut 2
IFT74 mut 3
IFT74 mut 4
IFT74 mut 5
D
GLI3repressor/fl
***
ns
SAG
wt
WDR60 nons. mut 1
WDR60 nons. mut 2
WDR60 nons. mut 3
E
wild type
WDR60 nons. mut1
WDR60 nons. mut2
WDR60 nons. mut3
M
170
130
100
70
55
40
Gli3 fl
Gli3 repressor
beta actin

Supplementary Figure 6:

(A-D) Gli3 repressor/ Glifl ratios before and after SAG treatment for each clone used in the study, shown in a pooled analysis in Figure 3. Band intensities of Gli3 full-length (Gli3fl) and Gli3 repressor were quantified by densitometry. The Gli3 repressor/ Gli3 fl ratio before and after SAG treatment was calculated for each sample. Bars represent mean ± SD from at least three independent experiments. Statistical significance was determined using two-way ANOVA with Sidak´s multiple comparisons test to compare the Gli3 repressor/ Gli3 fl ratio before and after SAG treatment (n= 3-4, *$p < 0.05$, ** $p < 0.01$, *** $p < 0.001$). (E) Representative Western blot of Gli fl and Gli repressor before and after SAG activation in wild type and WDR60 nonsense samples.

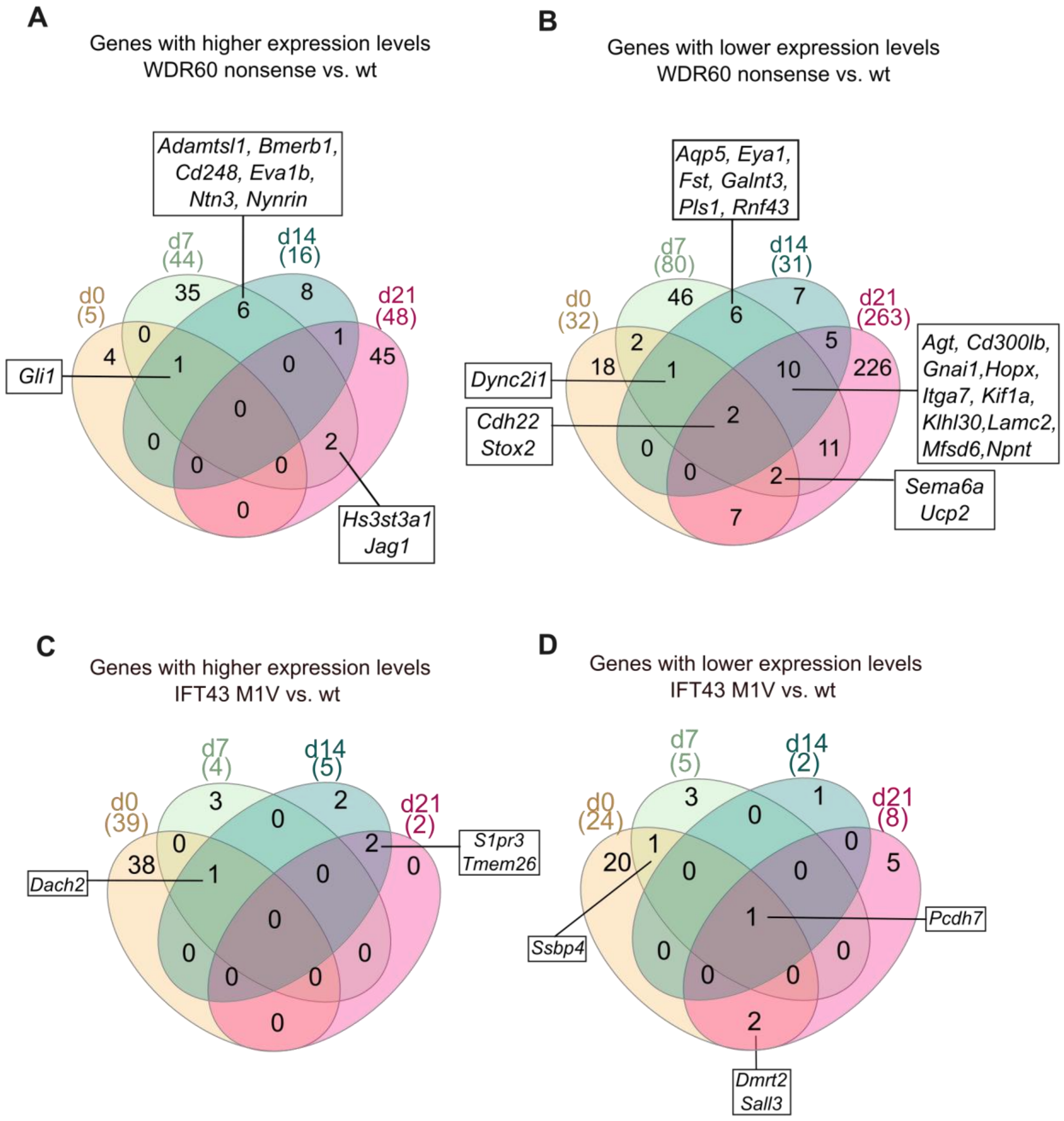


Supplementary Figure 7:

Venn diagrams illustrating overlap of genes with increased (A, C) or decreased (B, D) expression levels in WDR60 nonsense (A, B) and IFT43 M1V (C, D) mutants versus wilt type in transcriptomic analysis across different time points. All genes from intersections are listed in Suppl. File S1.

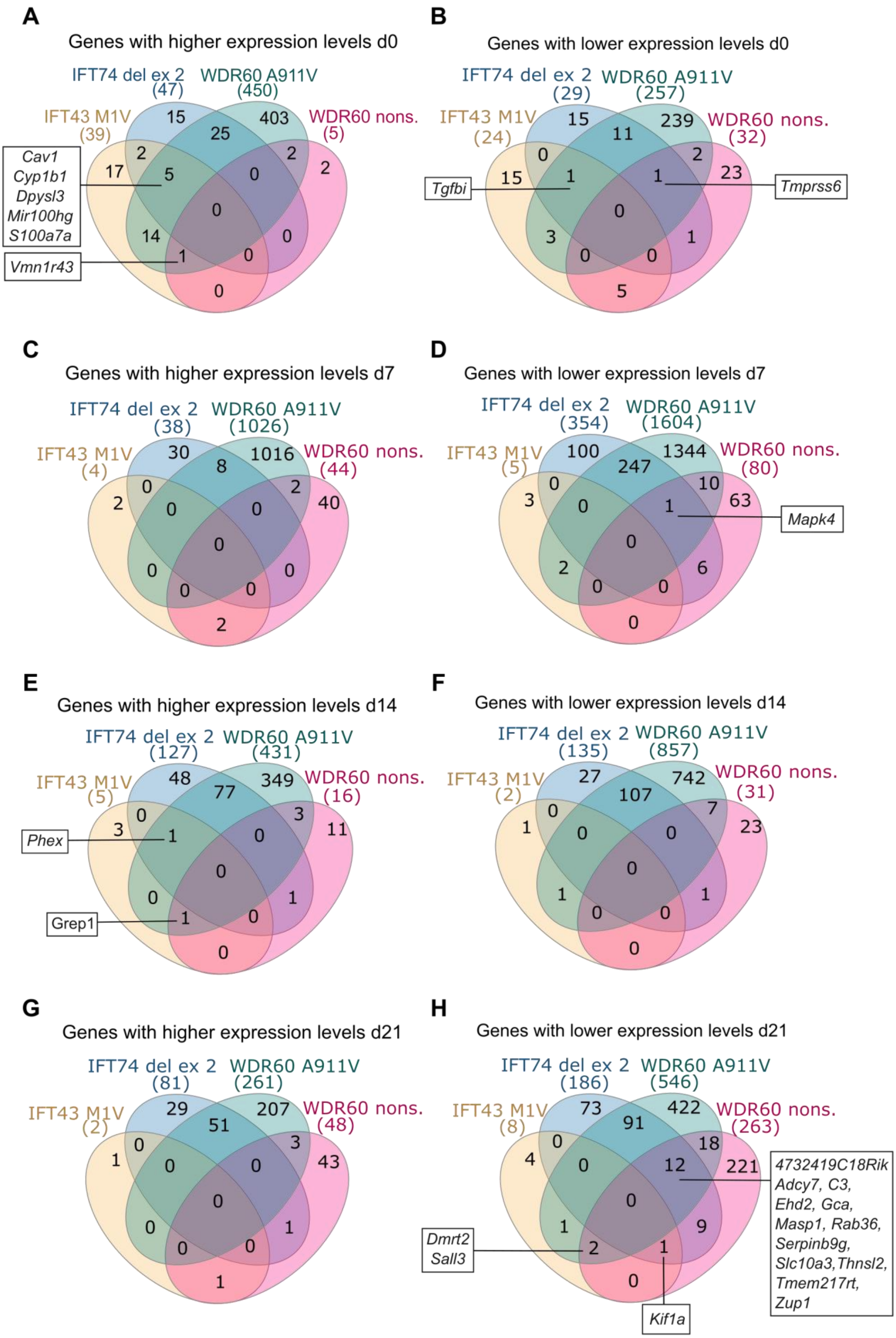
A
Genes with higher expression levels d0
IFT74 del ex 2 (47)
WDR60 A911V (450)
IFT43 M1V (39)
WDR60 nons. (5)
15
403
25
2
2
17
5
0
2
0
14
0
1
0
0
Cav1
Cyp1b1
Dpysl3
Mir100hg
S100a7a
Vmn1r43
B
Genes with lower expression levels d0
IFT74 del ex 2 (29)
WDR60 A911V (257)
IFT43 M1V (24)
WDR60 nons. (32)
15
239
11
0
2
23
15
1
1
0
3
1
0
0
5
Tgfbi
Tmprss6
C
Genes with higher expression levels d7
IFT74 del ex 2 (38)
WDR60 A911V (1026)
IFT43 M1V (4)
WDR60 nons. (44)
30
1016
8
0
2
2
0
0
40
0
0
0
0
0
2
D
Genes with lower expression levels d7
IFT74 del ex 2 (354)
WDR60 A911V (1604)
IFT43 M1V (5)
WDR60 nons. (80)
100
1344
247
0
10
3
0
1
63
0
2
6
0
0
0
Mapk4
E
Genes with higher expression levels d14
IFT74 del ex 2 (127)
WDR60 A911V (431)
IFT43 M1V (5)
WDR60 nons. (16)
48
349
77
0
3
3
1
0
11
0
0
1
1
0
0
Phex
Grep1
F
Genes with lower expression levels d14
IFT74 del ex 2 (135)
WDR60 A911V (857)
IFT43 M1V (2)
WDR60 nons. (31)
27
742
107
0
7
1
0
0
23
0
1
1
0
0
0
G
Genes with higher expression levels d21
IFT74 del ex 2 (81)
WDR60 A911V (261)
IFT43 M1V (2)
WDR60 nons. (48)
29
207
51
0
3
1
0
0
43
0
0
1
0
0
1
H
Genes with lower expression levels d21
IFT74 del ex 2 (186)
WDR60 A911V (546)
IFT43 M1V (8)
WDR60 nons. (263)
73
422
91
0
18
4
0
12
221
0
1
9
2
1
0
4732419C18Rik
Adcy7, C3,
Ehd2, Gca,
Masp1, Rab36,
Serpinb9g,
Slc10a3, Thnsl2,
Tmem217rt,
Zup1
Dmrt2
Sall3
Kif1a

Supplementary Figure 8:

(A-H) Venn diagrams illustrating overlap of genes with significantly higher expression levels (A, C, E, G, $p \leq 0.05$, $\log2fc \geq 1$) or lower expression levels (B, D, F, H, $p \leq 0.05$, $\log2fc \leq -1$) in mutants compared to wild type cells at indicated time points. Full lists of genes in intersections are provided in Suppl. File S1.

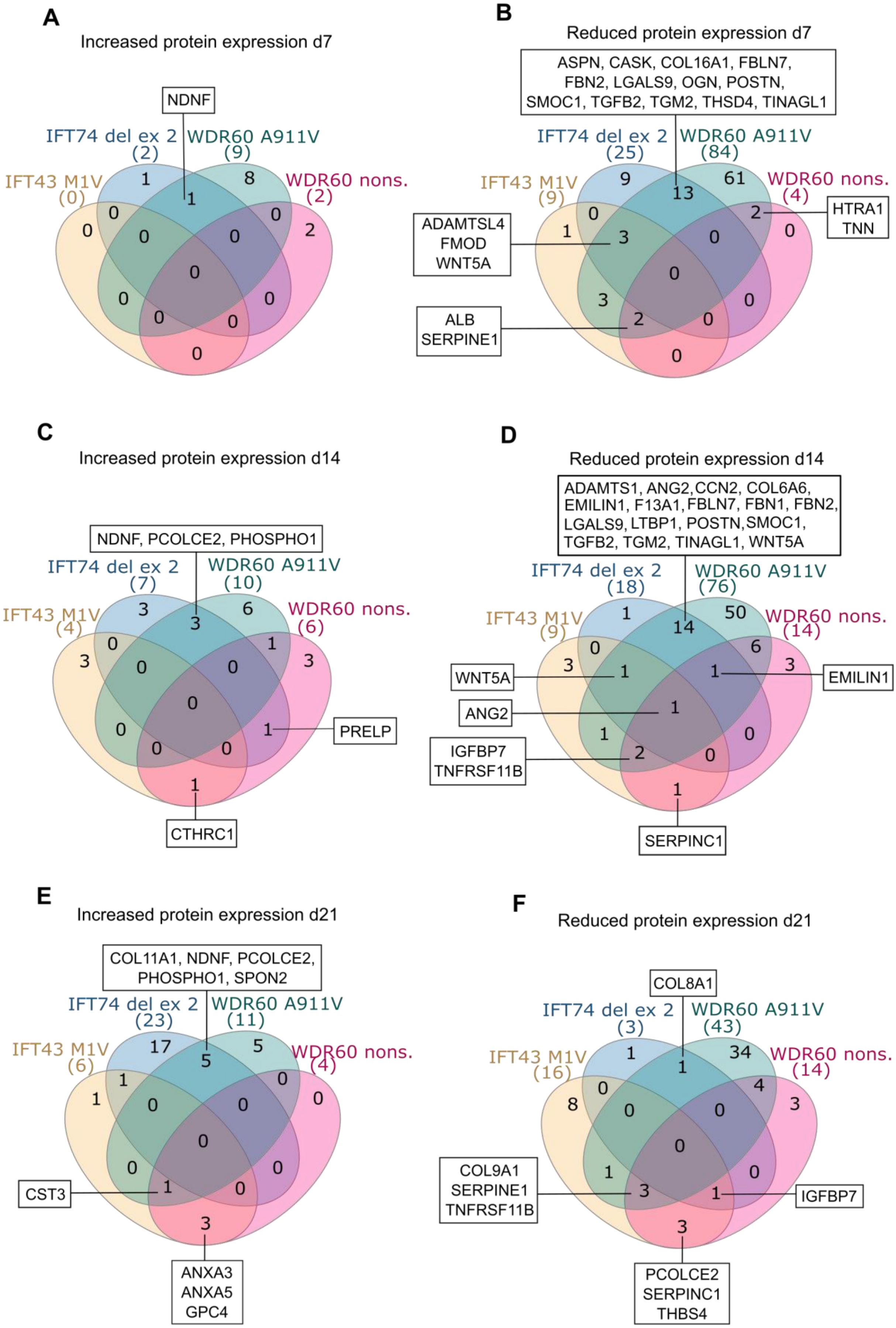

A
Increased protein expression d7
NDNF
IFT74 del ex 2 (2)
WDR60 A911V (9)
IFT43 M1V (0)
WDR60 nons. (2)
1 8 1 0 0 0 2 0 0 0 0 0 0 0 0
B
Reduced protein expression d7
ASPN, CASK, COL16A1, FBLN7, FBN2, LGALS9, OGN, POSTN, SMOC1, TGFB2, TGM2, THSD4, TINAGL1
IFT74 del ex 2 (25)
WDR60 A911V (84)
IFT43 M1V (9)
WDR60 nons. (4)
9 61 13 2 0 1 3 0 0 3 2 0 0 0
ADAMTSL4
FMOD
WNT5A
HTRA1
TNN
ALB
SERPINE1
C
Increased protein expression d14
NDNF, PCOLCE2, PHOSPHO1
IFT74 del ex 2 (7)
WDR60 A911V (10)
IFT43 M1V (4)
WDR60 nons. (6)
3 6 3 0 1 3 0 0 3 0 0 1 0 0 1
PRELP
CTHRC1
D
Reduced protein expression d14
ADAMTS1, ANG2, CCN2, COL6A6, EMILIN1, F13A1, FBLN7, FBN1, FBN2, LGALS9, LTBP1, POSTN, SMOC1, TGFB2, TGM2, TINAGL1, WNT5A
IFT74 del ex 2 (18)
WDR60 A911V (76)
IFT43 M1V (9)
WDR60 nons. (14)
1 50 14 0 6 3 1 1 3 1 1 0 2 0 1
WNT5A
EMILIN1
ANG2
IGFBP7
TNFRSF11B
SERPINC1
E
Increased protein expression d21
COL11A1, NDNF, PCOLCE2, PHOSPHO1, SPON2
IFT74 del ex 2 (23)
WDR60 A911V (11)
IFT43 M1V (6)
WDR60 nons. (4)
17 5 5 1 0 1 0 0 0 0 0 1 0 0 3
CST3
ANXA3
ANXA5
GPC4
F
Reduced protein expression d21
COL8A1
IFT74 del ex 2 (3)
WDR60 A911V (43)
IFT43 M1V (16)
WDR60 nons. (14)
1 34 1 0 4 8 0 0 3 0 1 0 3 1 3
COL9A1
SERPINE1
TNFRSF11B
IGFBP7
PCOLCE2
SERPINC1
THBS4

Supplementary Figure 9:

(A-F) Venn diagrams illustrating overlap of proteins with significantly higher expression levels (A, C, E, G, $p \leq 0.05$, log2fc ≥ 1) or lower expression levels (B, D, F, H, $p \leq 0.05$, log2fc ≤ -1) in mutants compared to wild type ECM samples at indicated time points. Full list of proteins in intersections are provided in Suppl. File S2.

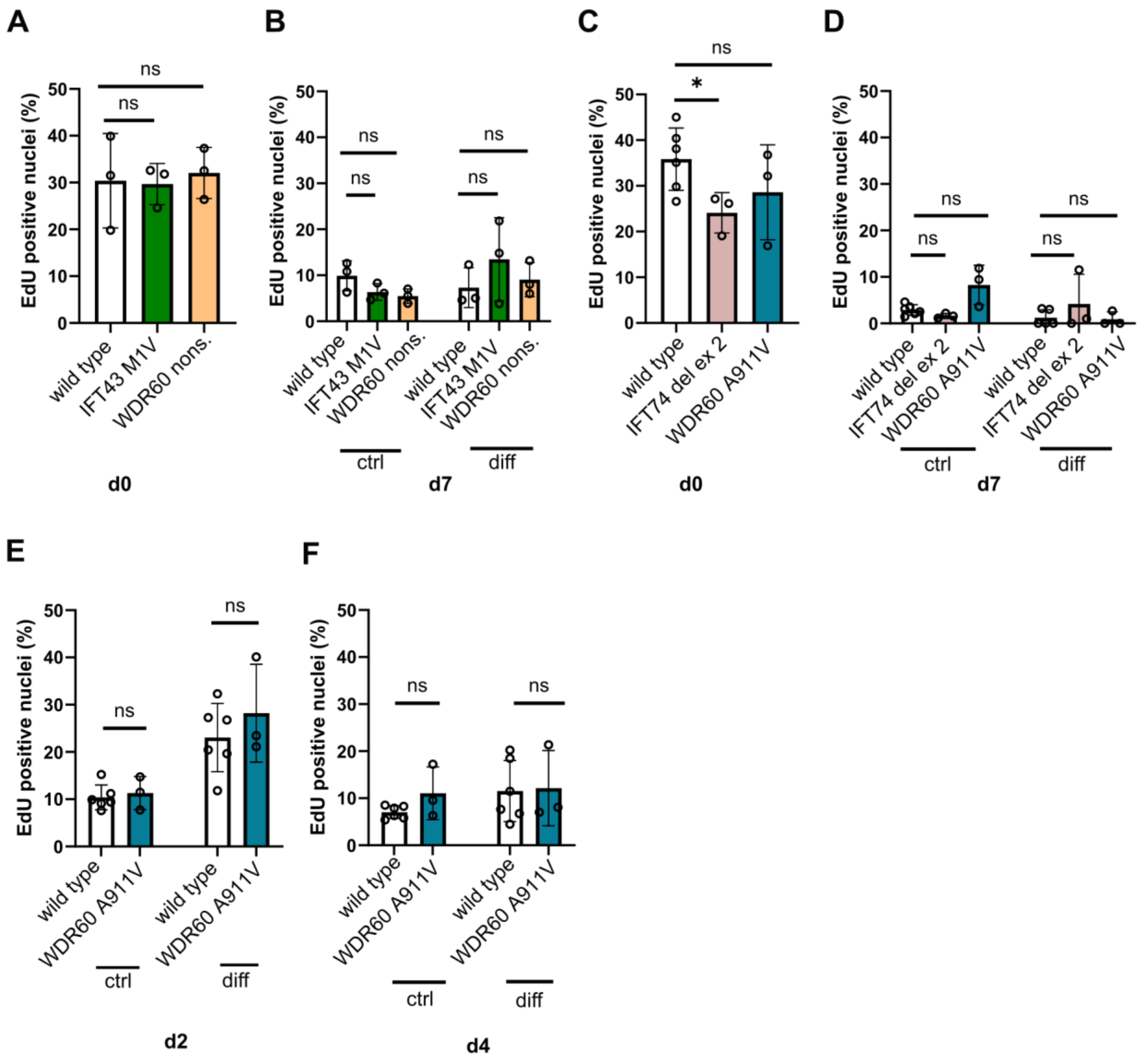


Supplementary Figure 10:

(A-D) Proportion of EdU positive nuclei of all nuclei was determined at day 0 (A, C) and day 7 (B, D) after 2 h EdU incubation. For each genotype three independent clones were analyzed. Images were taken at random locations on the slide. For each genotype, at least three images comprising a minimum of 100 nuclei in total were analyzed. (E-F) EdU positive nuclei in WDR60 A911V mutants and wt samples at day 2 (E) and day 4 (F). Statistical significance was determined using unpaired t-test (n=3-6, *p < 0.05).

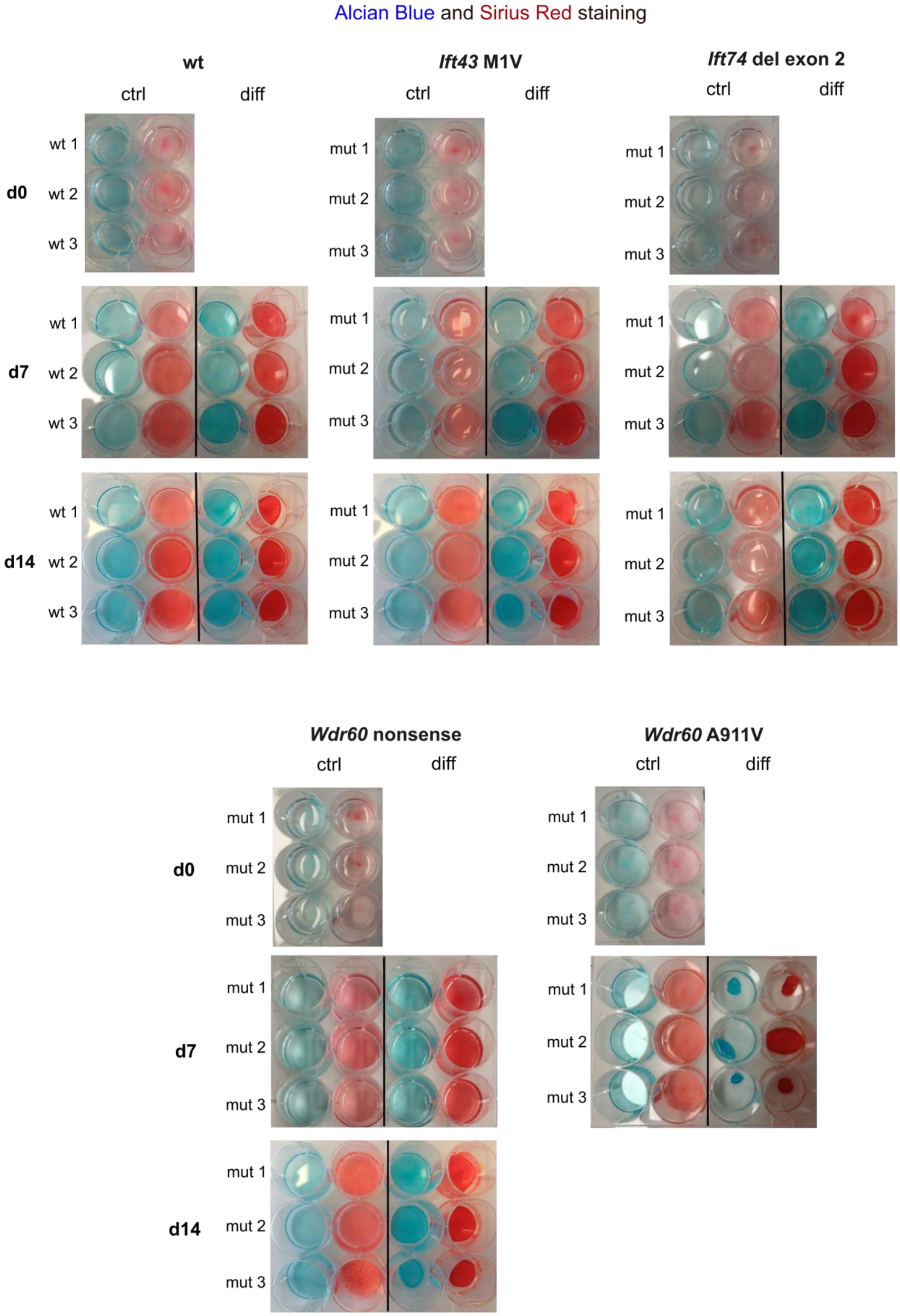

Alcian Blue and Sirius Red staining
wt
Ift43 M1V
Ift74 del exon 2
ctrl
diff
d0
d7
d14
wt 1
wt 2
wt 3
mut 1
mut 2
mut 3
Wdr60 nonsense
Wdr60 A911V

Supplementary Figure 11:

Alcian Blue and Sirius Red staining at day 0, 7 and 14 of three different clones per genotype. Cells were kept under control conditions (5% FCS medium) and differentiation conditions (differentiation medium) for the respective time periods. For WDR60 A911V clones, no staining later than day 7 could be performed due to detachment of differentiating cells.

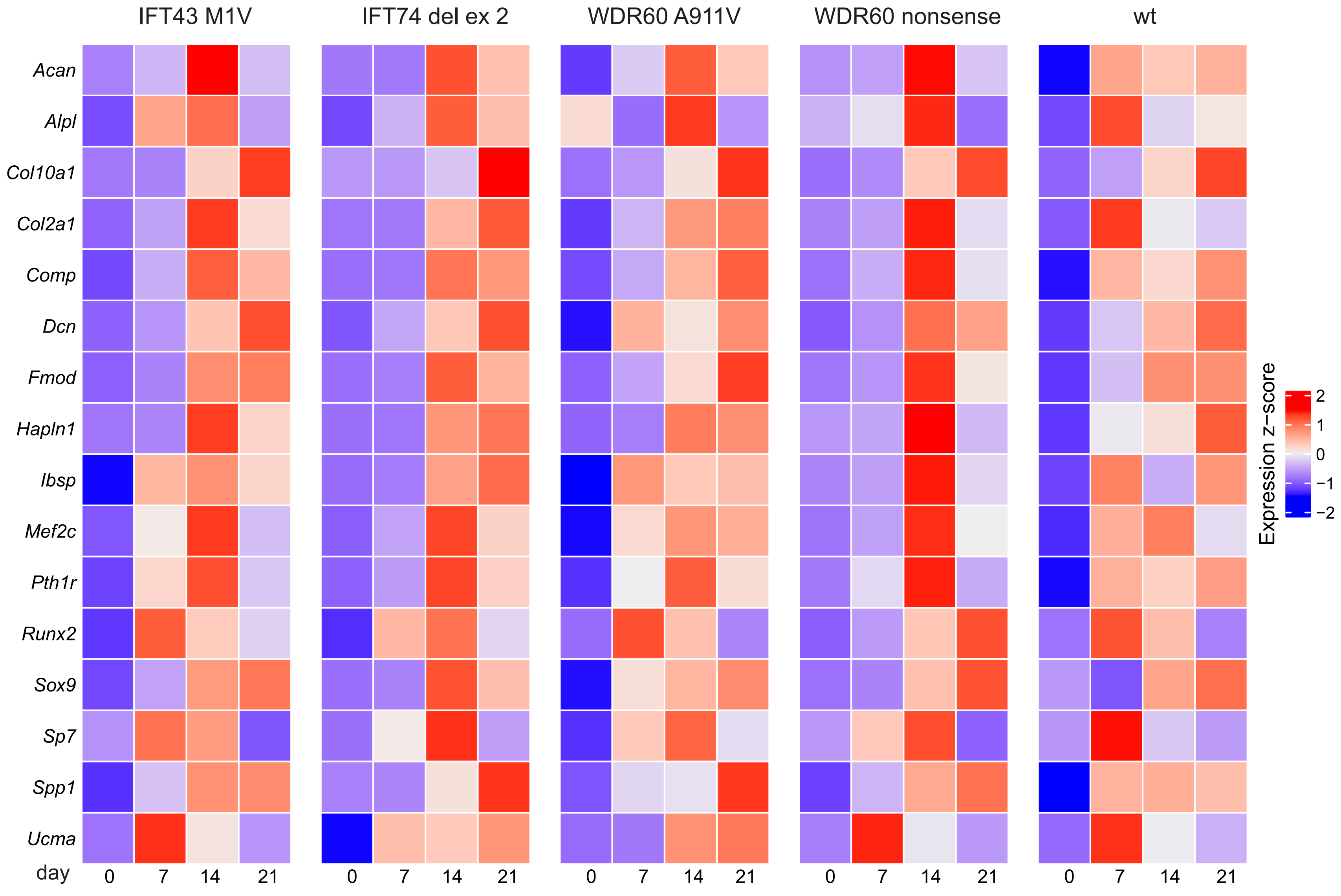
IFT43 M1V
IFT74 del ex 2
WDR60 A911V
WDR60 nonsense
wt
Acan
Alpl
Col10a1
Col2a1
Comp
Dcn
Fmod
Hapln1
Ibsp
Mef2c
Pth1r
Runx2
Sox9
Sp7
Spp1
Ucma
day
0
7
14
21
Expression z−score
2
1
0
−1
−2

Supplementary Figure 12:

Heatmap showing the expression dynamics of selected chondrocyte marker genes in IFT43 M1V, IFT74 del ex 2, WDR60 A911V, WDR60 nonsense, and wild-type cells across the differentiation time course. DESeq2-normalized counts were used to calculate z-scores for each gene across the time course. Z-scores reflect relative expression changes within each gene and do not allow direct comparison of expression levels between genotypes.

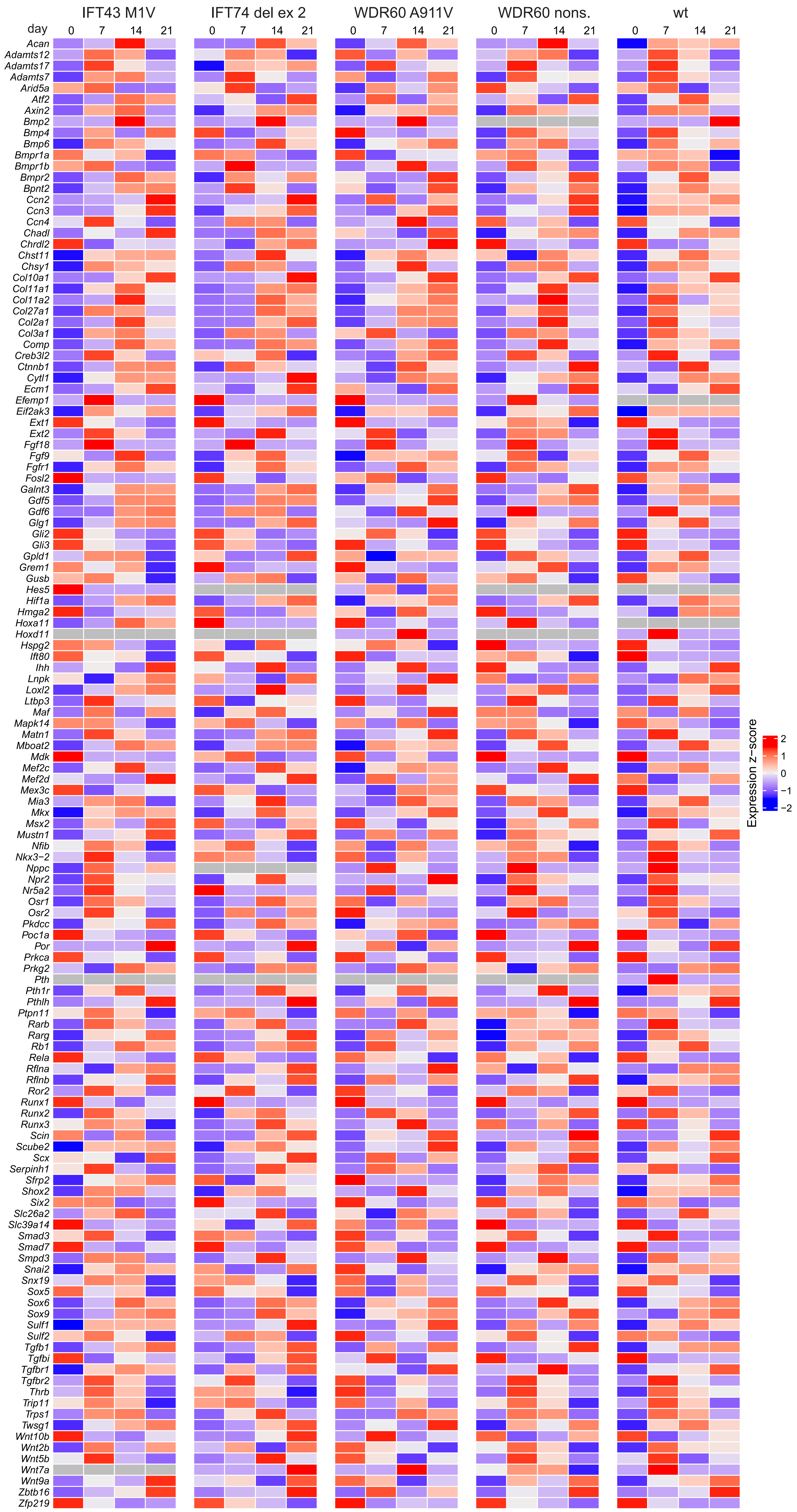
IFT43 M1V
IFT74 del ex 2
WDR60 A911V
WDR60 nons.
wt
day
0
7
14
21
Acan
Adamts12
Adamts17
Adamts7
Arid5a
Atf2
Axin2
Bmp2
Bmp4
Bmp6
Bmpr1a
Bmpr1b
Bmpr2
Bpnt2
Ccn2
Ccn3
Ccn4
Chadl
Chrdl2
Chst11
Chsy1
Col10a1
Col11a1
Col11a2
Col27a1
Col2a1
Col3a1
Comp
Creb3l2
Ctnnb1
Cytl1
Ecm1
Efemp1
Eif2ak3
Ext1
Ext2
Fgf18
Fgf9
Fgfr1
Fosl2
Galnt3
Gdf5
Gdf6
Glg1
Gli2
Gli3
Gpld1
Grem1
Gusb
Hes5
Hif1a
Hmga2
Hoxa11
Hoxd11
Hspg2
Ift80
Ihh
Lnpk
Loxl2
Ltbp3
Maf
Mapk14
Matn1
Mboat2
Mdk
Mef2c
Mef2d
Mex3c
Mia3
Mkx
Msx2
Mustn1
Nfib
Nkx3−2
Nppc
Npr2
Nr5a2
Osr1
Osr2
Pkdcc
Poc1a
Por
Prkca
Prkg2
Pth
Pth1r
Pthlh
Ptpn11
Rarb
Rarg
Rb1
Rela
Rflna
Rflnb
Ror2
Runx1
Runx2
Runx3
Scin
Scube2
Scx
Serpinh1
Sfrp2
Shox2
Six2
Slc26a2
Slc39a14
Smad3
Smad7
Smpd3
Snai2
Snx19
Sox5
Sox6
Sox9
Sulf1
Sulf2
Tgfb1
Tgfbi
Tgfbr1
Tgfbr2
Thrb
Trip11
Trps1
Twsg1
Wnt10b
Wnt2b
Wnt5b
Wnt7a
Wnt9a
Zbtb16
Zfp219
Expression z−score
2
1
0
−1
−2

Supplementary Figure 13:

Heatmap displaying the expression of genes annotated in GO:0002062 “chondrocyte differentiation” + Col10a1 in IFT43 M1V, IFT74 del ex 2, WDR60 A911V, WDR60 nonsense and wild type cells over the course of the differentiation process in alphabetical order. DESeq2-normalized counts were used to calculate z-scores for each gene across the time course. Z-scores reflect relative expression changes within each gene and do not allow direct comparison of expression levels between genotypes. Grey areas indicate no detectable expression of the gene at the respective time point.

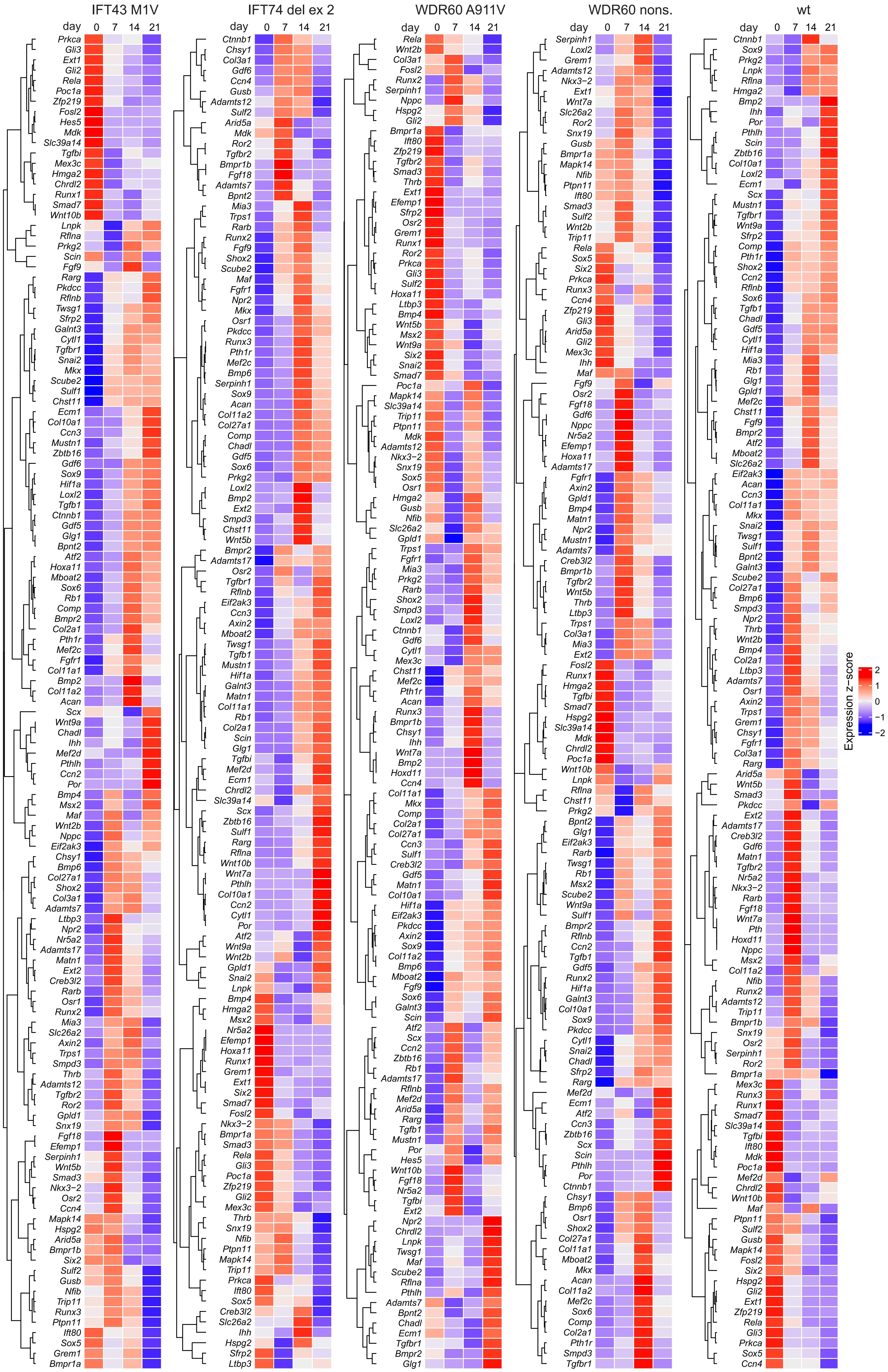
IFT43 M1V
IFT74 del ex 2
WDR60 A911V
WDR60 nons.
wt
day 0 7 14 21
Expression z-score
2
1
0
−1
−2

Supplementary Figure 14:

Clustered heatmap displaying the expression of genes annotated in GO:0002062 “chondrocyte differentiation” + Col10a1 in IFT43 M1V, IFT74 del ex 2, WDR60 A911V, WDR60 nonsense and wild type cells over the course of the differentiation process. DESeq2-normalized counts were used to calculate z-scores for each gene across the time course. Z-scores reflect relative expression changes within each gene and do not allow direct comparison of expression levels between genotypes. Grey areas indicate no detectable expression of the gene at the respective time point.

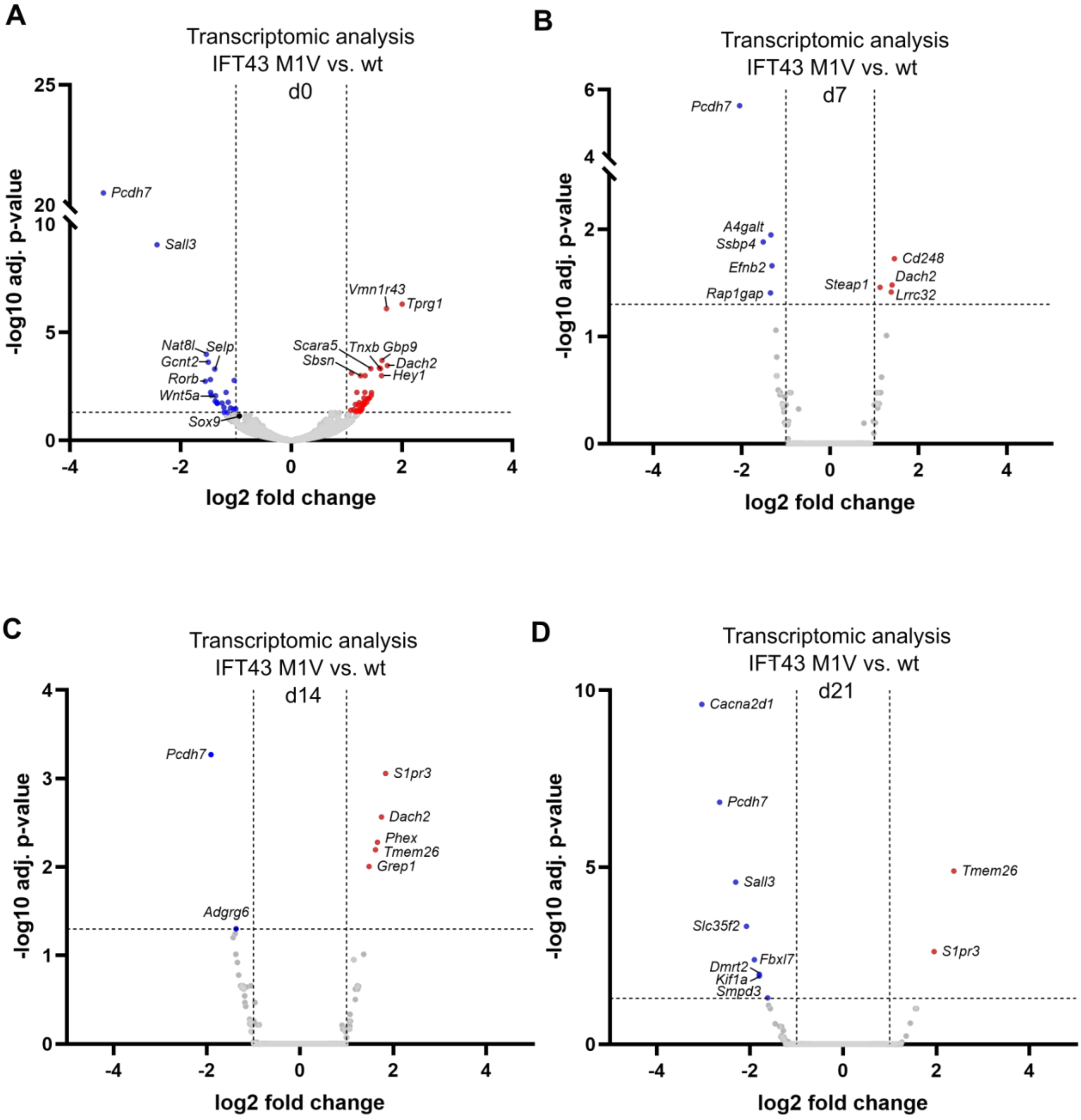


Supplementary Figure 15:

Volcano plots illustrating transcriptomic analysis at day 0 (A), day 7 (B), day 14 (C) and day 21 (D) for IFT43 mutant clones compared to wild type. Genes were color-coded according to log2fold change thresholds of ±1 and adjusted p-value ≤ 0.05. Parts of this figure are also shown in Figure 5.

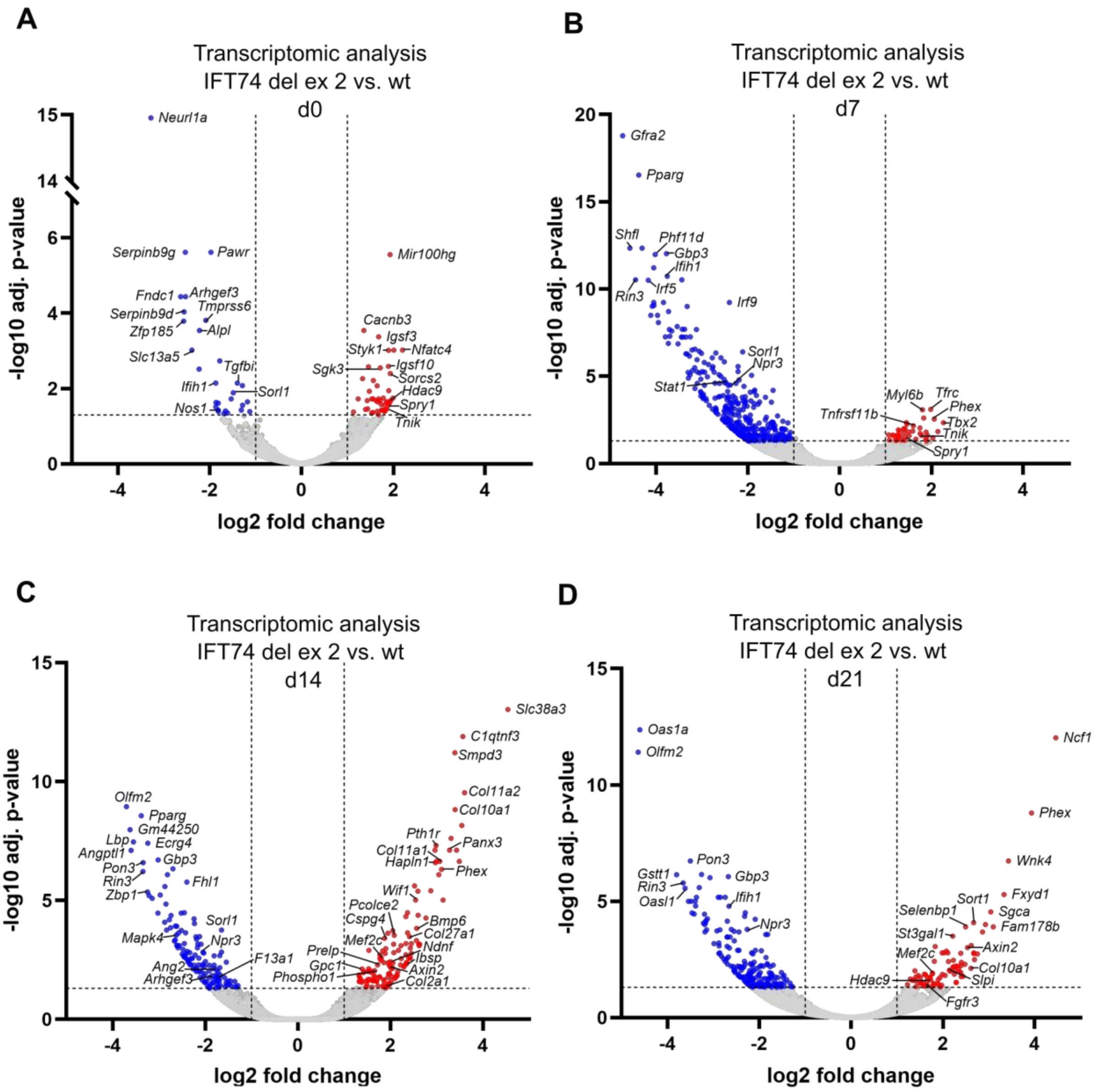


Supplementary Figure 16:

Volcano plots illustrating transcriptomic analysis at day 0 (A), day 7 (B), day 14 (C) and day 21 (D) for IFT74 del ex 2 mutant clones compared to wild type. Genes were color-coded according to log2fold change thresholds of ±1 and adjusted p-value ≤ 0.05. Parts of this figure are shown in Figure 5.

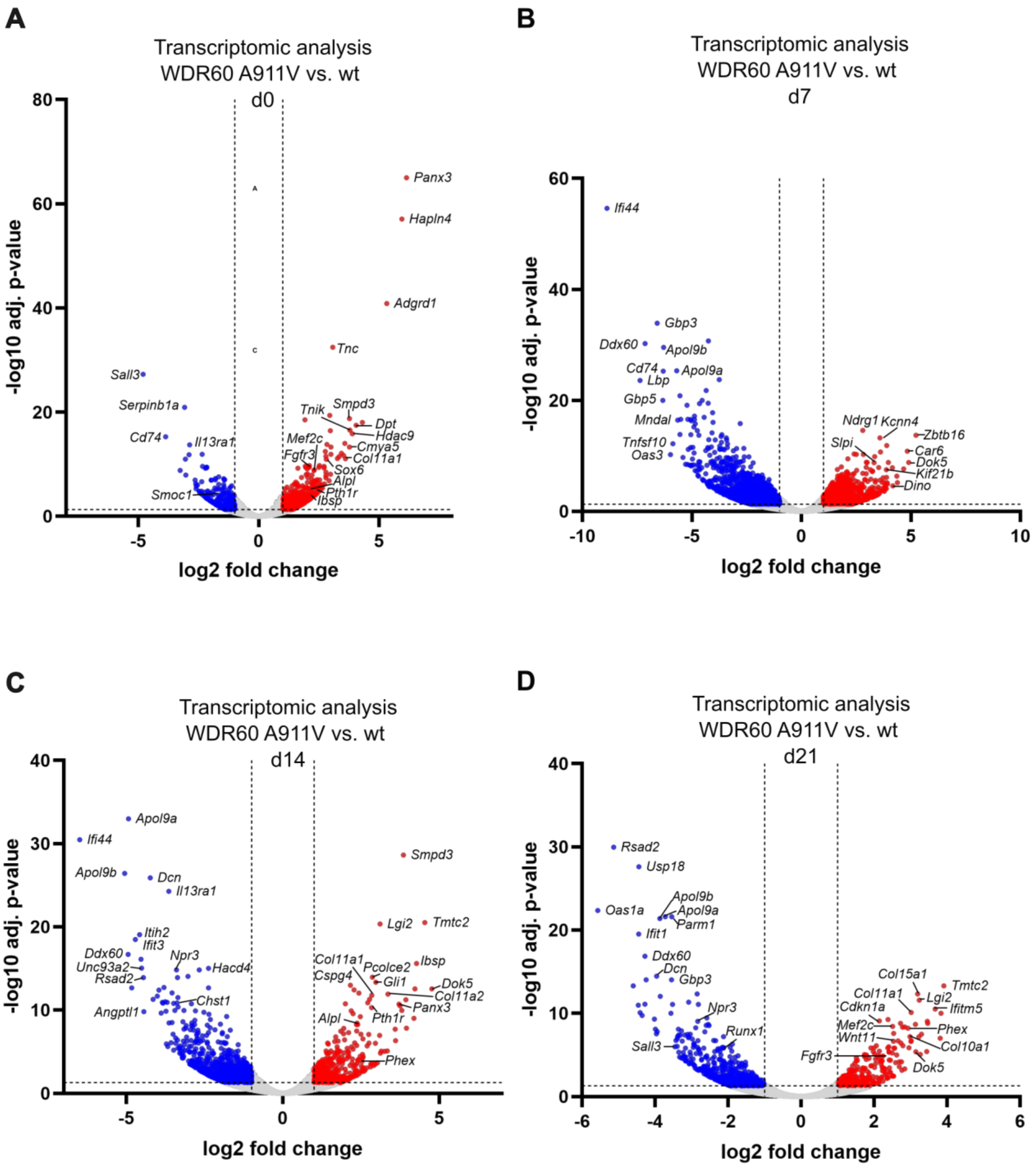


Supplementary Figure 17:

Volcano plots illustrating transcriptomic analysis at day 0 (A), day 7 (B), day 14 (C) and day 21 (D) for WDR60 A911V mutant clones compared to wild type. Genes were color-coded according to log2fold change thresholds of ±1 and adjusted p-value ≤ 0.05. Parts of this figure are shown in Figure 6.

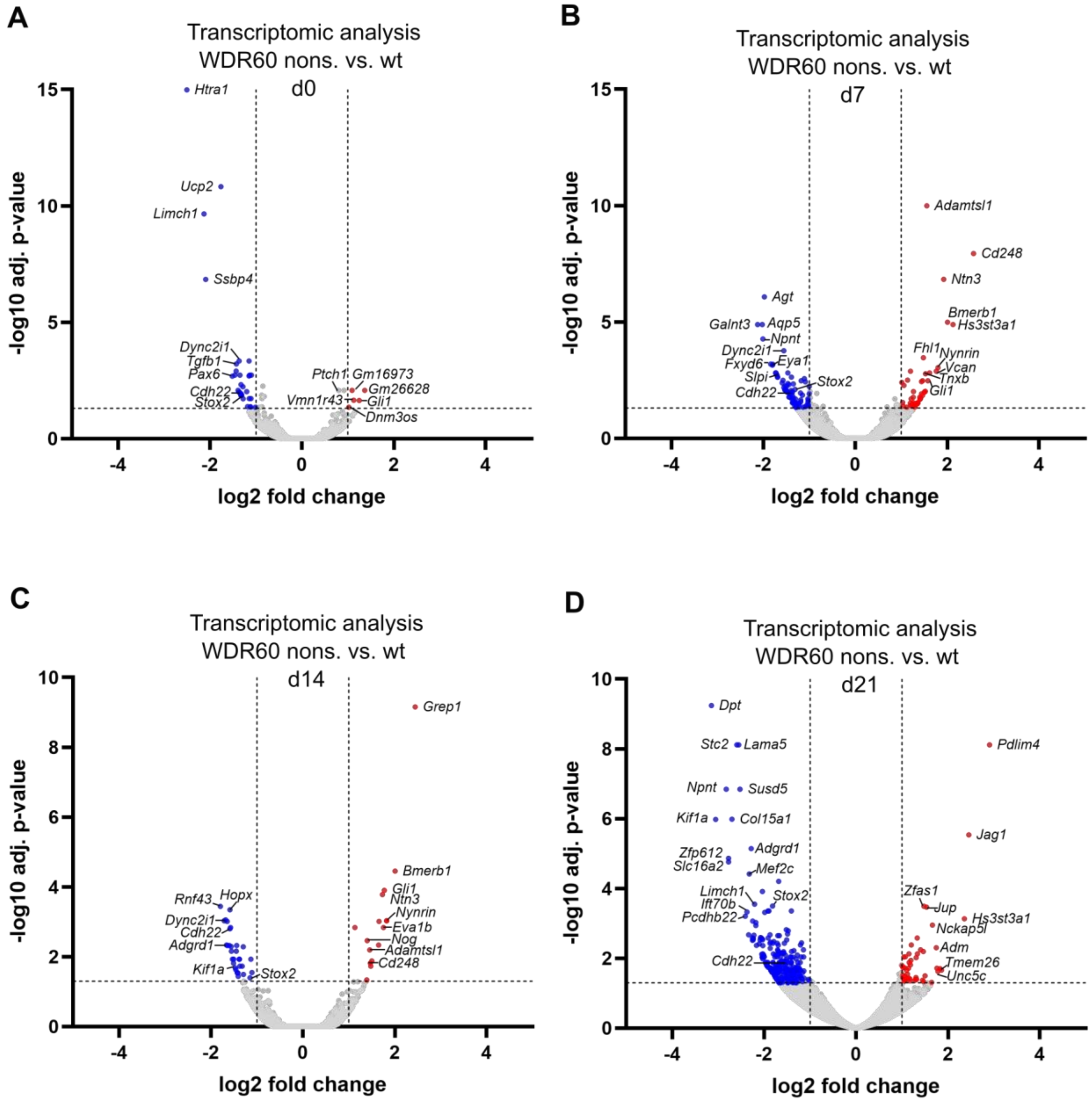


Supplementary Figure 18:

Volcano plots illustrating transcriptomic analysis at day 0 (A), day 7 (B), day 14 (C) and day 21 (D) for WDR60 nonsense mutant clones compared to wild type. Genes were color-coded according to log2fold change thresholds of ±1 and adjusted p-value ≤ 0.05. Parts of this figure are shown in Figure 5.

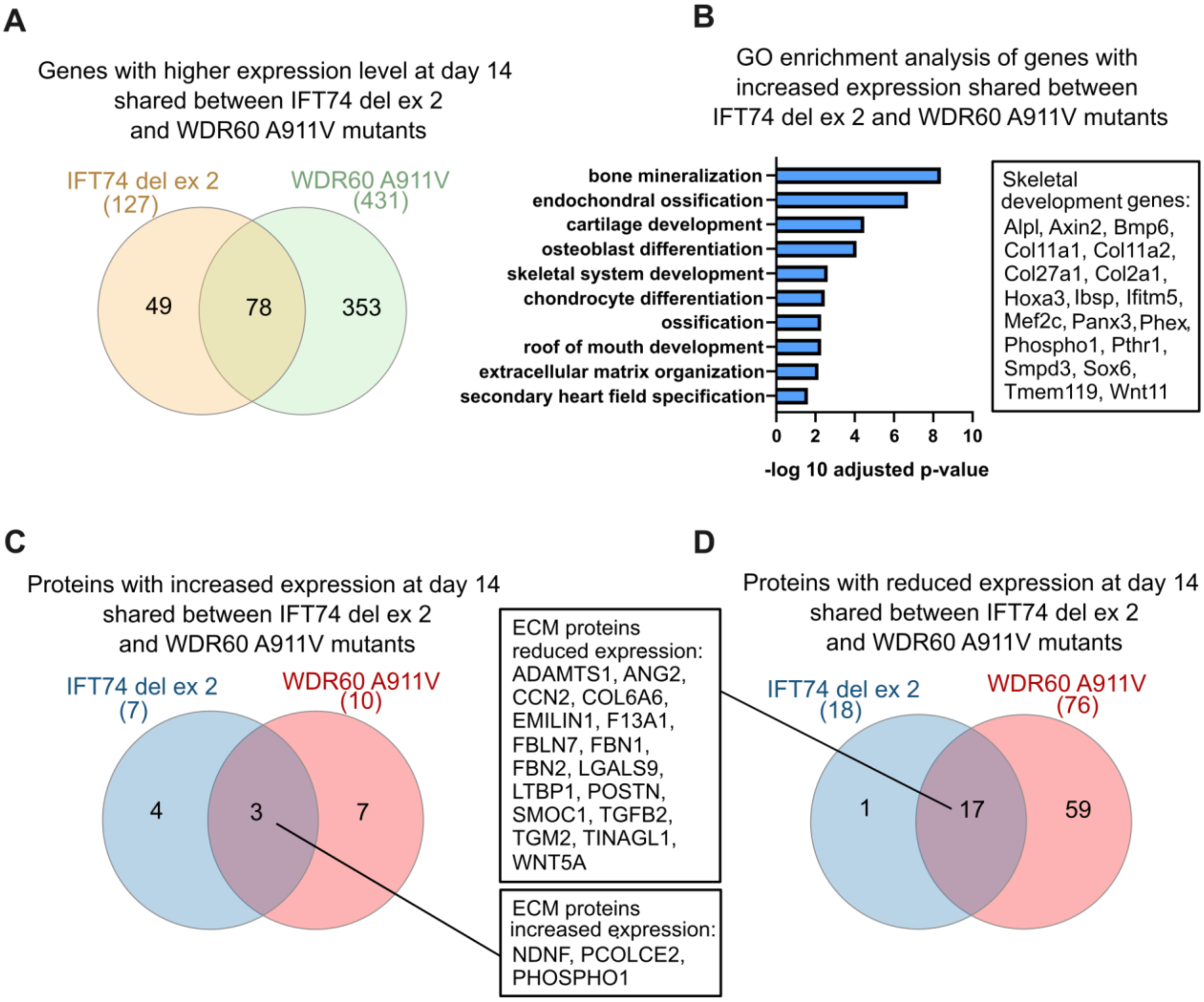


Supplementary Figure 19

(A) Venn diagram illustrating shared genes with increased expression in IFT74 del ex 2 and WDR60 A911V mutants compared to wild type at day 14. (B) GO analysis of genes with increased expression compared to wild type shared between IFT74 del ex 2 and WDR60 A911V mutants. Top 10 most significant GO terms are displayed. The complete list of GO terms and associated genes is provided in Suppl. File 5. Representative skeletal development genes from GO terms GO:0030282~bone mineralization, GO:0001958~endochondral ossification, GO:0051216~cartilage development, GO:0001649~osteoblast differentiation and GO:0001501~skeletal system development are indicated. (C, D) Venn diagrams showing overlap of ECM proteins (GO:0031012) with increased (C) and decreased (D) expression in IFT74 del ex 2 and WDR60 A911V samples compared to wild type at day 14.

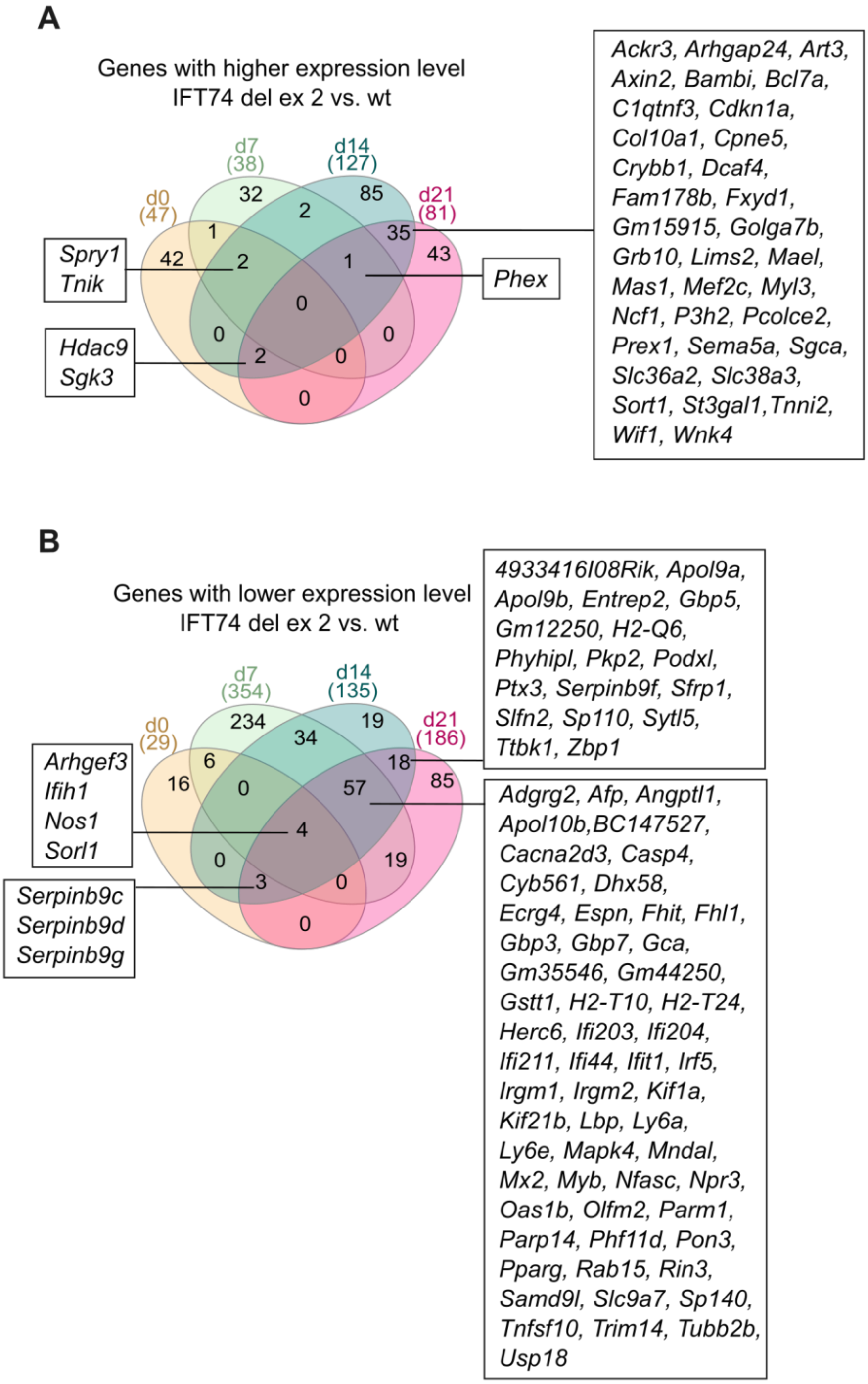


Supplementary Figure 20:

Venn diagrams illustrating overlap of genes with increased (A) or decreased (B) expression levels in IFT74 del ex 2 mutants versus wild type in transcriptomic analyses across different time points. All genes from intersections are listed in Supplementary File 1.

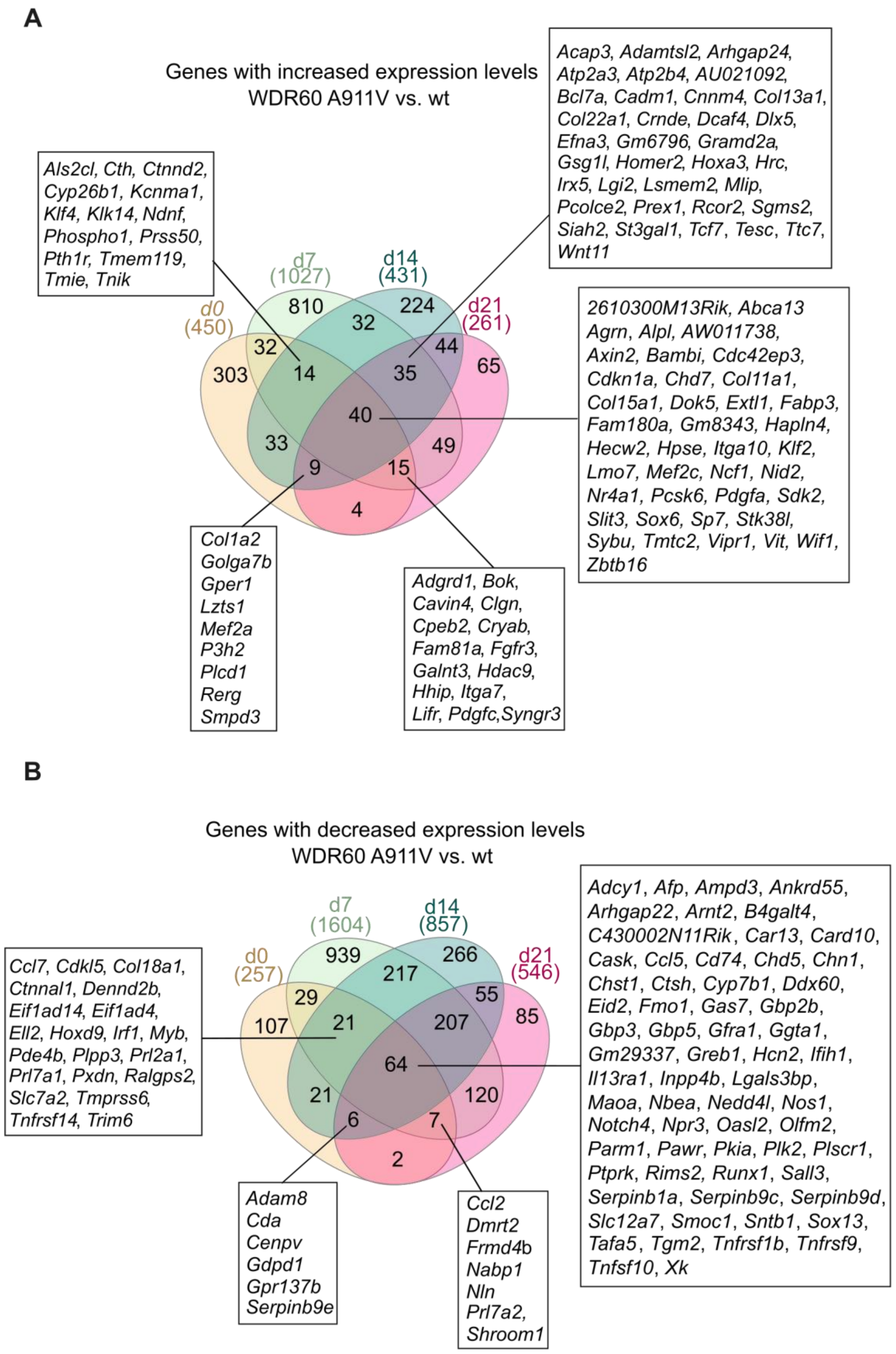


Supplementary Figure 21:

Venn diagrams illustrating overlap of genes with increased (A) or decreased (B) expression levels in WDR60 A911V mutants versus wilt type in transcriptomic analysis across different time points. All genes from intersections are listed in Supplementary File 1.

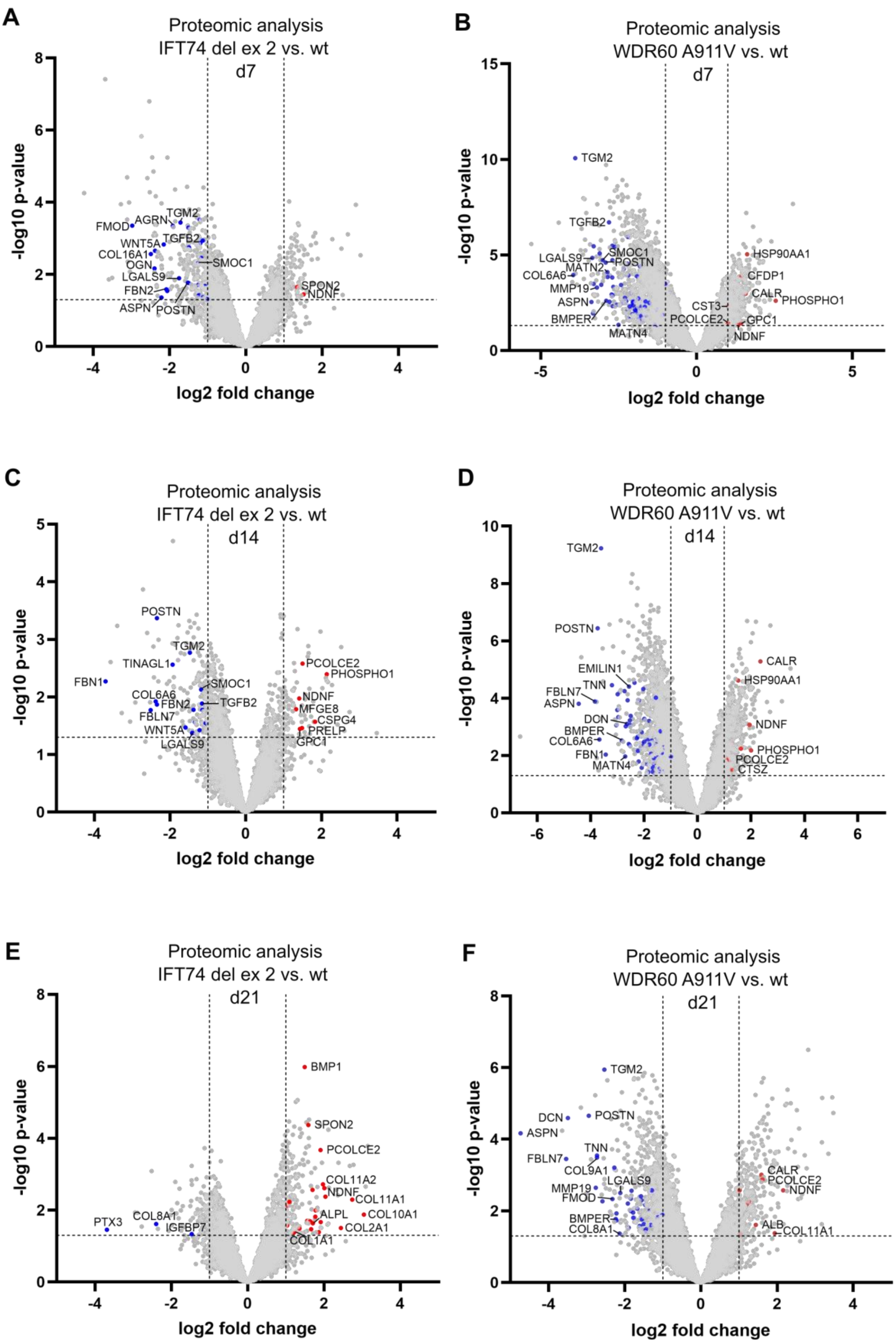
A
Proteomic analysis
IFT74 del ex 2 vs. wt
d7
-log10 p-value
log2 fold change
FMOD
AGRN
TGM2
WNT5A
TGFB2
COL16A1
OGN
SMOC1
LGALS9
FBN2
ASPN
POSTN
SPON2
NDNF
B
Proteomic analysis
WDR60 A911V vs. wt
d7
TGM2
TGFB2
LGALS9
SMOC1
MATN2
POSTN
COL6A6
MMP19
ASPN
BMPER
MATN4
HSP90AA1
CFDP1
CALR
PHOSPHO1
CST3
PCOLCE2
GPC1
NDNF
C
Proteomic analysis
IFT74 del ex 2 vs. wt
d14
POSTN
TGM2
TINAGL1
FBN1
SMOC1
COL6A6
FBN2
TGFB2
FBLN7
WNT5A
LGALS9
PCOLCE2
PHOSPHO1
NDNF
MFGE8
CSPG4
PRELP
GPC1
D
Proteomic analysis
WDR60 A911V vs. wt
d14
TGM2
POSTN
EMILIN1
TNN
FBLN7
ASPN
DCN
BMPER
COL6A6
FBN1
MATN4
CALR
HSP90AA1
NDNF
PHOSPHO1
PCOLCE2
CTSZ
E
Proteomic analysis
IFT74 del ex 2 vs. wt
d21
BMP1
SPON2
PCOLCE2
COL11A2
NDNF
COL11A1
ALPL
COL10A1
COL2A1
COL1A1
PTX3
COL8A1
IGFBP7
F
Proteomic analysis
WDR60 A911V vs. wt
d21
TGM2
DCN
POSTN
ASPN
TNN
FBLN7
COL9A1
LGALS9
MMP19
FMOD
BMPER
COL8A1
CALR
PCOLCE2
NDNF
ALB
COL11A1

Supplementary Figure 22:

Volcano plots illustrating proteomic analyses of IFT74 del ex 2 vs. wt (A, C, E) and WDR60 A911V vs. wt (B, D, F) at different time points during the differentiation process after selecting ECM proteins (GO:0031012 extracellular matrix). Grey dots represent proteins not annotated in (GO:0031012 extracellular matrix). ECM proteins were color-coded according to log2fold change thresholds of ±1 and p value ≤ 0.05. Differentially expressed protein lists are provided in Suppl. File 4.

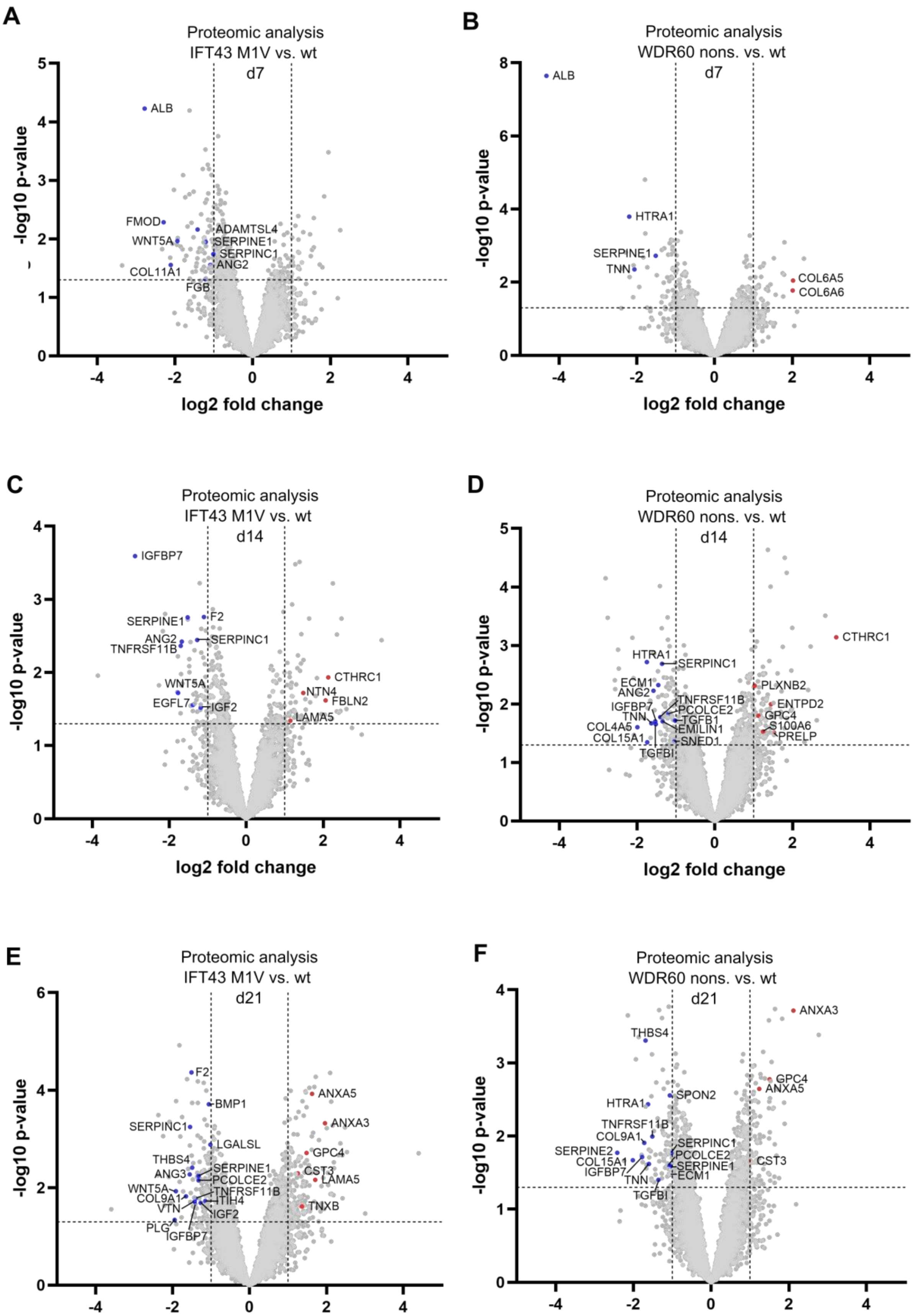
A
Proteomic analysis
IFT43 M1V vs. wt
d7
-log10 p-value
log2 fold change
ALB
FMOD
WNT5A
COL11A1
ADAMTSL4
SERPINE1
SERPINC1
ANG2
FGB
B
Proteomic analysis
WDR60 nons. vs. wt
d7
-log10 p-value
log2 fold change
ALB
HTRA1
SERPINE1
TNN
COL6A5
COL6A6
C
Proteomic analysis
IFT43 M1V vs. wt
d14
-log10 p-value
log2 fold change
IGFBP7
SERPINE1
F2
ANG2
SERPINC1
TNFRSF11B
WNT5A
EGFL7
IGF2
CTHRC1
NTN4
FBLN2
LAMA5
D
Proteomic analysis
WDR60 nons. vs. wt
d14
-log10 p-value
log2 fold change
CTHRC1
HTRA1
SERPINC1
ECM1
ANG2
IGFBP7
TNN
COL4A5
COL15A1
TGFBI
TNFRSF11B
PCOLCE2
TGFB1
EMILIN1
SNED1
PLXNB2
ENTPD2
GPC4
S100A6
PRELP
E
Proteomic analysis
IFT43 M1V vs. wt
d21
-log10 p-value
log2 fold change
F2
BMP1
SERPINC1
LGALSL
THBS4
ANG3
SERPINE1
PCOLCE2
WNT5A
COL9A1
VTN
TNFRSF11B
ITIH4
IGF2
PLG
IGFBP7
ANXA5
ANXA3
GPC4
CST3
LAMA5
TNXB
F
Proteomic analysis
WDR60 nons. vs. wt
d21
-log10 p-value
log2 fold change
ANXA3
THBS4
GPC4
ANXA5
SPON2
HTRA1
TNFRSF11B
COL9A1
SERPINC1
SERPINE2
PCOLCE2
COL15A1
SERPINE1
IGFBP7
ECM1
TNN
TGFBI
CST3

Supplementary Figure 23:

Volcano plots illustrating proteomic analyses of IFT43 M1V vs. wt (A, C, E) and WDR60 nonsense vs. wt (B, D, F) at different time points during the differentiation process after selecting ECM proteins (GO:0031012 extracellular matrix). Grey dots represent proteins not annotated in (GO:0031012 extracellular matrix). ECM proteins were color-coded according to log2fold change thresholds of ±1 and p value ≤ 0.05. Differentially expressed protein lists are provided in Suppl. File 4.

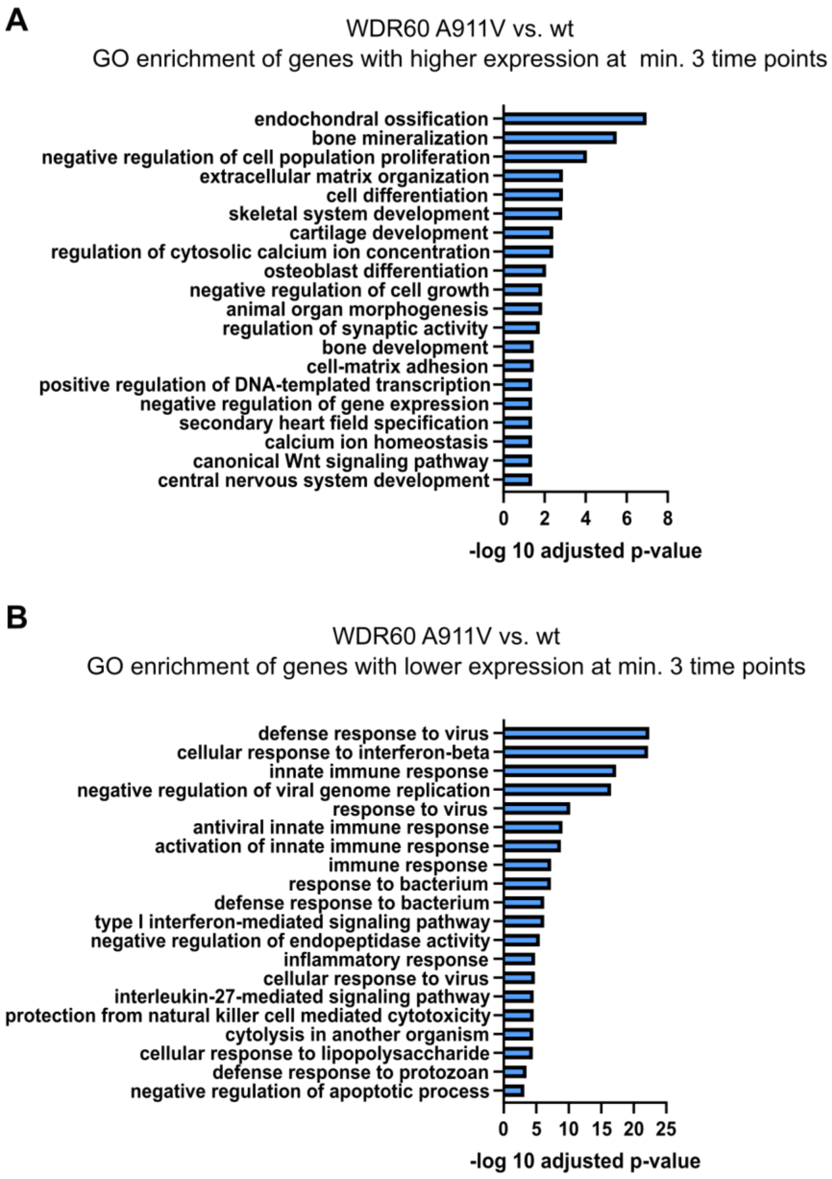


Supplementary Figure 24:

Top 20 significant GO terms from GO analysis with genes that show consistently higher (A) or lower (B) expression levels at ≥ 3 time points in WDR60 A911V vs. wt samples. Complete lists of GO terms and associated genes are provided in Tables S5 and S6.

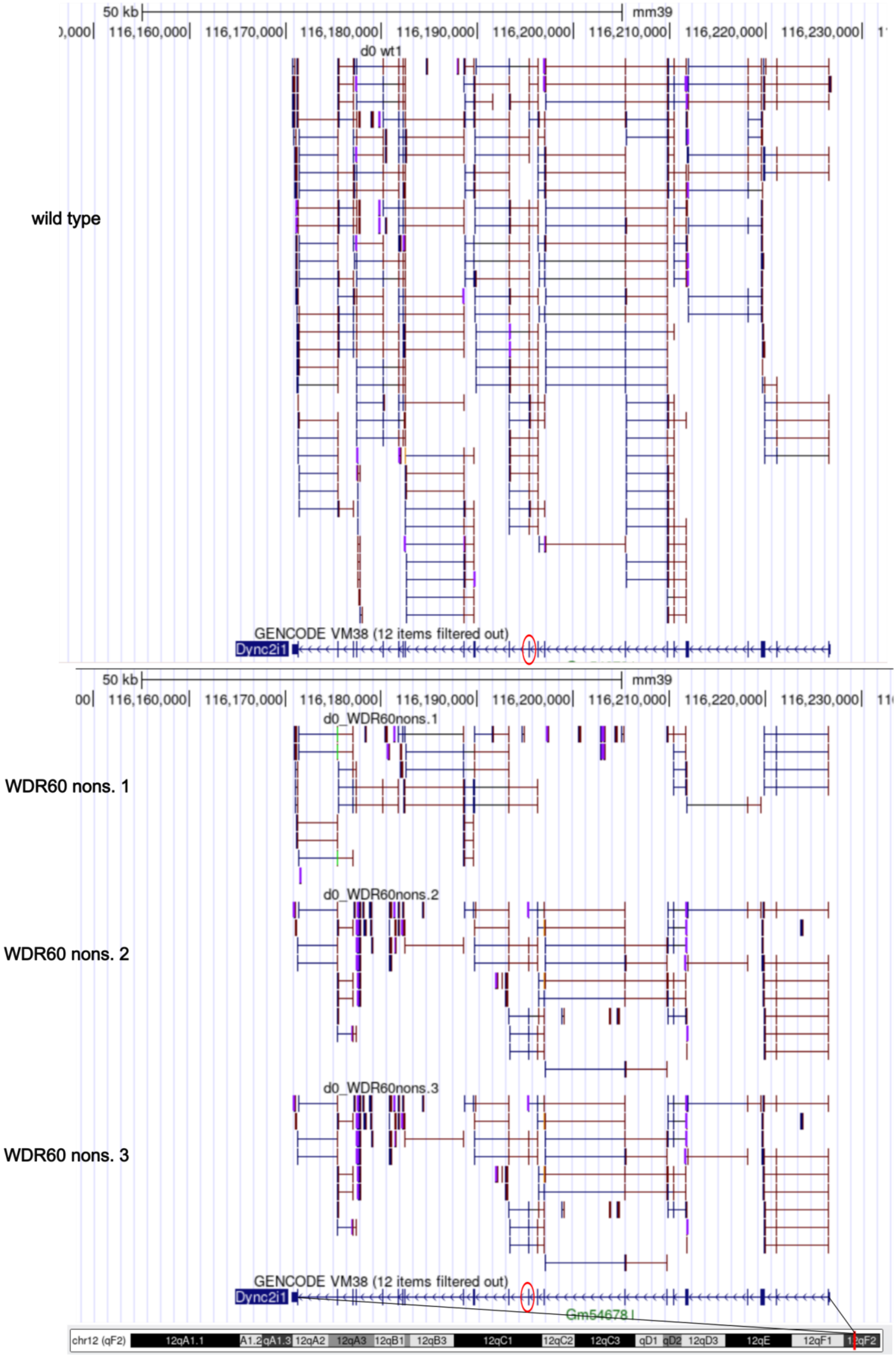

wild type
WDR60 nons. 1
WDR60 nons. 2
WDR60 nons. 3
50 kb
mm39
116,160,000
116,170,000
116,180,000
116,190,000
116,200,000
116,210,000
116,220,000
116,230,000
d0 wt1
d0_WDR60nons.1
d0_WDR60nons.2
d0_WDR60nons.3
GENCODE VM38 (12 items filtered out)
Dync2i1
Gm54678
chr12 (qF2)
12qA1.1
12qA2
12qA3
12qB1
12qB3
12qC1
12qC2
12qC3
12qD3
12qE
12qF1

Supplementary Figure 25:

Visualization of RNA-seq read alignments to the mouse genome (mm39) in the UCSC Genome Browser for one wild-type clone and three WDR60 nonsense clones at day 0. Exon 11, corresponding to the CRISPR gRNA target site, is highlighted by a red circle.

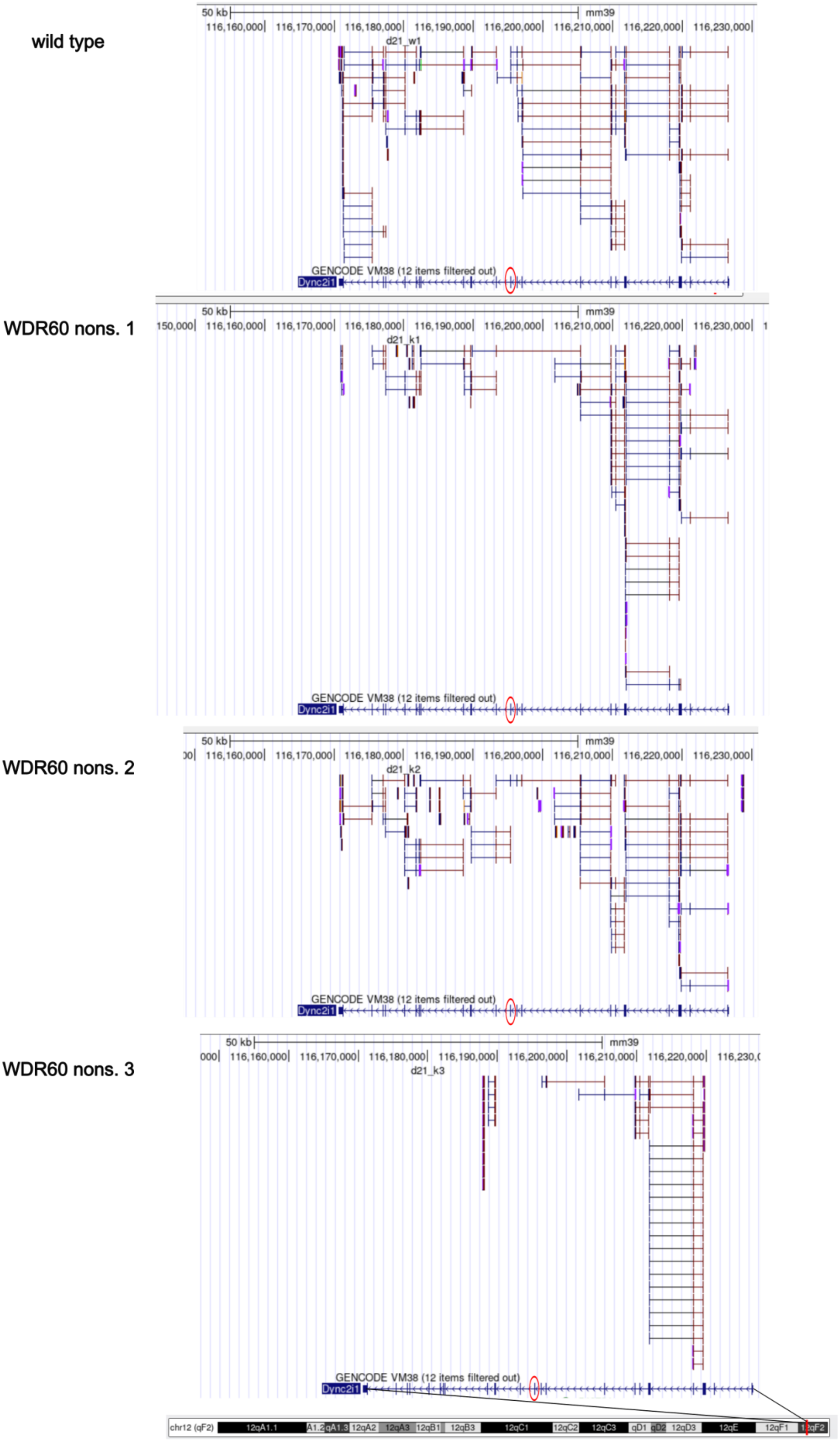
wild type
WDR60 nons. 1
WDR60 nons. 2
WDR60 nons. 3
50 kb
mm39
116,160,000
116,170,000
116,180,000
116,190,000
116,200,000
116,210,000
116,220,000
116,230,000
d21_w1
d21_k1
d21_k2
d21_k3
GENCODE VM38 (12 items filtered out)
Dync2i1
chr12 (qF2)
12qA1.1
12qA2
12qA3
12qB1
12qB3
12qC1
12qC2
12qC3
qD1
qD2
12qD3
12qE
12qF1

Supplementary Figure 26:

Visualization of RNA-seq read alignments to the mouse genome (mm39) in the UCSC Genome Browser for one wild-type clone and three WDR60 nonsense clones at day 21. Exon 11, corresponding to the CRISPR gRNA target site, is highlighted by a red circle.

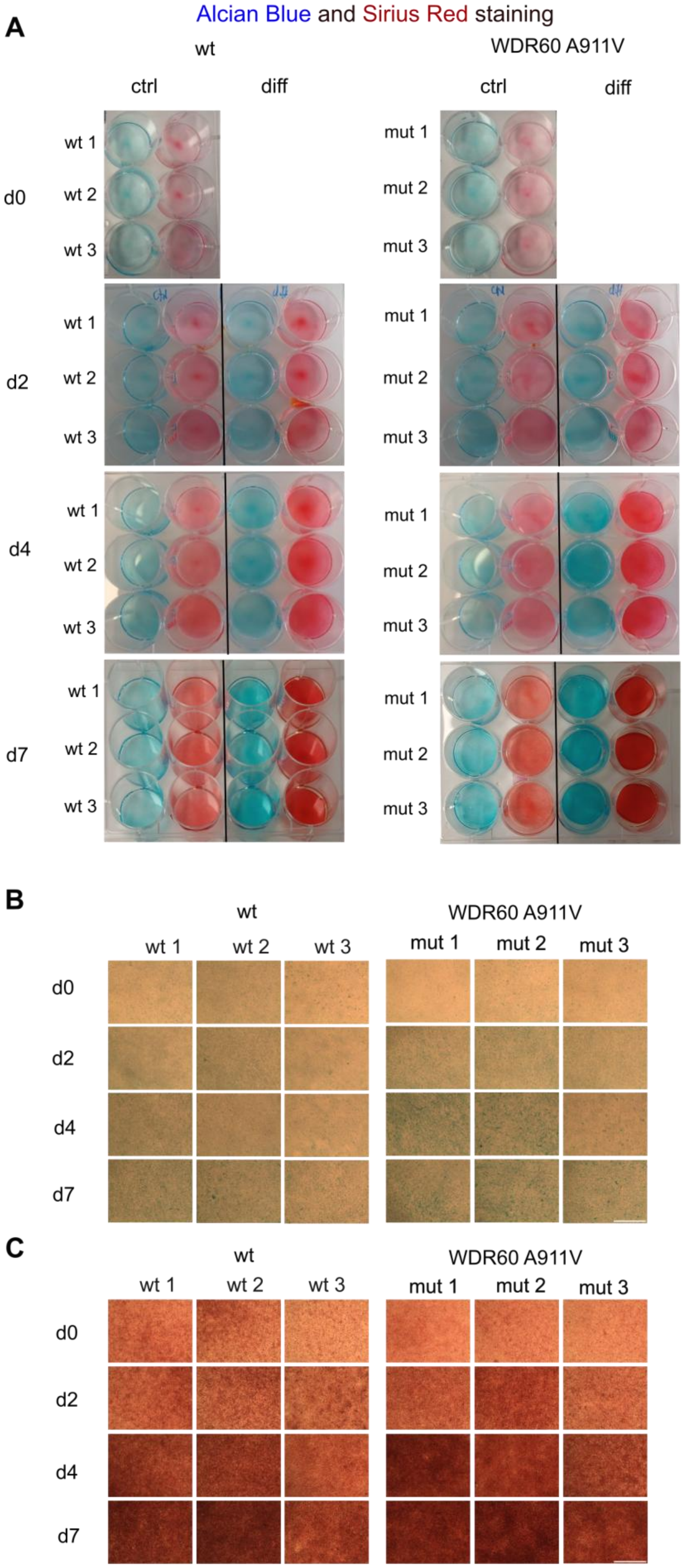
A
Alcian Blue and Sirius Red staining
wt
WDR60 A911V
ctrl
diff
ctrl
diff
d0
d2
d4
d7
wt 1
wt 2
wt 3
mut 1
mut 2
mut 3
B
wt
WDR60 A911V
wt 1
wt 2
wt 3
mut 1
mut 2
mut 3
d0
d2
d4
d7
C
wt
WDR60 A911V
wt 1
wt 2
wt 3
mut 1
mut 2
mut 3
d0
d2
d4
d7

Supplementary Figure 27:

(A) Alcian Blue and Sirius Red staining at day 0, 2, 4 and 7 of three independent clones per genotype. Cells were kept under control conditions (5% FCS medium) and differentiation conditions (differentiation medium) for the respective time period. (B, C) Representative light microscopic images of Alcian Blue- (B) and Sirius Red-stained (C) ATDC5 wild type and WDR60 A911V cultures after differentiation for the indicated time period. Scale bars: 1000 µm.

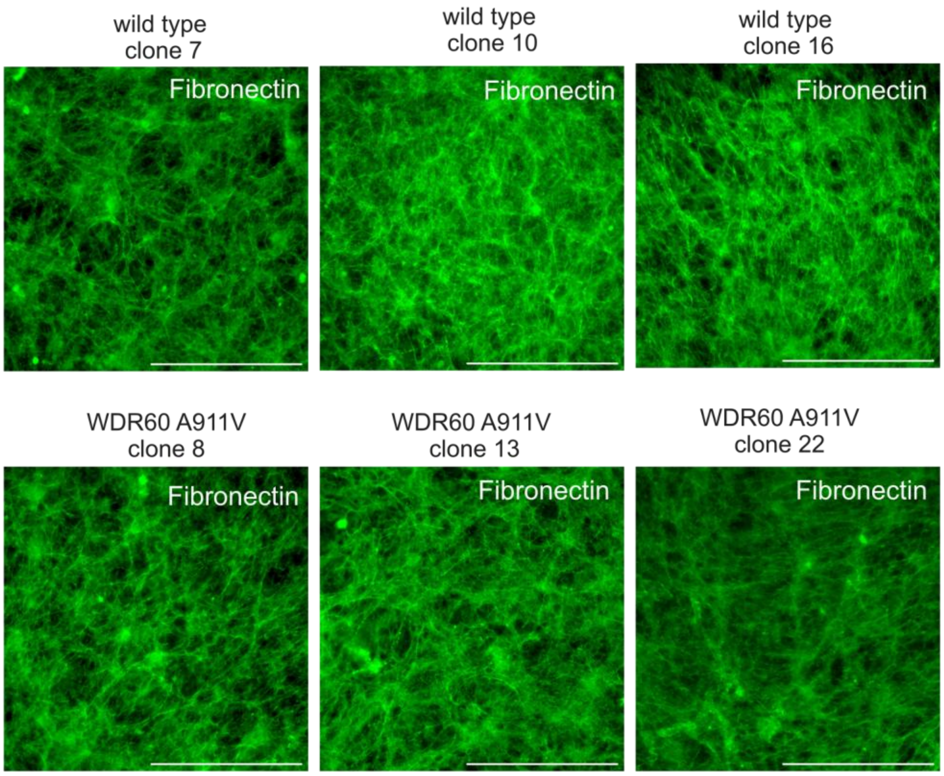


Supplementary Figure 28:

Fibronectin visualisation using immunofluorescence analysis of three different wild type and WDR60 A911V mutant clones after 7 days of differentiation. Cells were stained with rabbit anti fibronectin antibody (1:500, ab2413, abcam). Scale bars: 100 µm. Images were taken using a Leica Thunder microscope. Images were acquired under identical conditions; contrast was uniformly enhanced in ImageJ for display purposes.
