## supplementary methods and tables for "Distinct effects of hypomorphic IFT and dynein-2 skeletal ciliopathy disease alleles on chondrogenic differentiation, ECM composition and wnt signalling in ATDC5 derived cartilage-like organoids"

**Cell culture**

ATDC5 cells were obtained from ECACC. Cells were cultured in medium composed of Dulbecco’s modified eagle medium (DMEM, 32430100, Gibco) and Ham´s F-12 (11765054, Gibco) in a ratio of 1:1, supplemented with 10 % fetal calf serum (FCS, F7524 Sigma), 1% Sodium Pyruvate (S8636, Sigma), 100 U/ml penicillin and 100 µg/ml streptomycin (P-0781, Sigma) at 37°C and 5% CO_2_. The cells were passed every 3-4 days.

For differentiation experiments, ATDC5 cells were plated at 3×10^5^ or 0.75×10^5^ cells/well into 6 or 12-well plates (83.3920/ 83.392, Sarstedt). Upon confluency, the growth mediums FCS concentration was reduced to 5% and was supplemented with 10µg/ml insulin (I9278, Sigma), 30nM sodium selenite (S5261, Sigma), 50µg/ml ascorbic acid (A4403, Sigma) and 10µg/ml transferrin (90190, Sigma). The medium was changed every other day.

**Generation of CRISPR/Cas9 mutant ATDC5 cell lines**

ATDC5 cells were edited using CRISPR or CRISPR base editing. In this study, three cell lines were generated, each carrying a hypomorphic patient-derived mutation in one component of the IFT-A, IFT-B, or dynein-2 complex. Additionally, one WDR60 KO cell line was created. For recreation of WDR60 A911V and IFT43 M1V patient mutations, CRISPR base editing was used to change a single base at the desired position in the genome. For generation of WDR60 KO cells and IFT74 del exon 2 mutants a classic CRISPR approach was used to induce a frameshift mutation. CRISPR Cas9 plasmids, base editors and backbones were obtained from Addgene, if not stated otherwise. All used CRISPR plasmids are listed in Supplementary table 7.

For IFT43 M1V and WDR60 A911V gene editing, BPK1520 (Addgene #65777) plasmids containing gRNA sequences to target the respective mutation sites, were previously cloned in our laboratory (gRNA details in Supplementary table 8). For IFT74 del exon 2 gene editing, gRNA was previously cloned into plasmid PX458 (Addgene #48138). The gRNA containing plasmid for WDR60 KO gene editing was synthesized and purchased from Vectorbuilder (VB240123-1105xhz).

For transfection, cells were seeded into a 6-well plate at a density of 0.35 × 10⁶ cells per well. The next day, cells were transfected with 4µg of the respective CRISPR Cas9 or base editor plasmid and 1µg of BPK1520 containing gRNA information in case of IFT43, WDR60 A911V and WDR60 KO. For the deletion of exon 2 in IFT74, 5 µg of the edited PX458 plasmid, encoding both the gRNA and CRISPR-Cas9, was transfected. The plasmids were mixed with FuGene transfection reagent (E2311, Promega) and dropwise added to the media. After 24 hours the media was changed. 48 hours after the first transfection the cells were once more reseeded and transfected the next day. Successful transfection was ensured by the observation of GFP under the fluorescence microscope. 72 hours after the second transfection, single cell sorting was performed and the single cells collected in 96 well plates. Around three weeks later, colonies obtained from a single cell, were picked, expanded and their genotype was determined. Genotype was determined by Sanger sequencing. Purity of the colonies was further ensured by performing TOPO cloning (45-0030, Life Technologies), according to manufacturers instructions. For each mutation, at least three independent clones were used in experiments to compensate clone-specific effects. Clones that underwent the CRISPR editing procedure but did not carry a mutation at the intended sites were taken as wild type controls.

**Proteoglycan and collagen staining**

Cells were washed twice with PBS (D8537, Sigma) and fixed with ice-cold methanol. At indicated time points, cells were stained for proteoglycans and collagens using 0.1% Alcian Blue (A5268, Sigma) in 0.1M HCl and Sirius Red solution (13422, Morphisto), for 2 hours at room temperature. Following incubation, excess dye was removed. Alcian Blue-stained cells were rinsed with PBS, while Sirius Red-stained cells were washed with 0.01 M HCl to remove unbound dye. Images of the stained cultures were captured using a Leica microscope or phone camera.

**Immunohistochemistry**

For investigation of cilia length, ciliation efficiency and IFT88 accumulation, cells were serum starved for 32 hours with 0% FCS, prior to fixation. Cells were seeded on cover slips or chamber slides and maintained until they reached confluency. First, cells were washed three times with PBS. Then the cells were fixed with ice cold methanol followed by permeabilization with 0.5% Triton in PBS. After blocking with 5% BSA in PBS solution the fixed cells were stained with suitable antibodies.

The fixed cells were stained with rabbit polyclonal IFT88 (13967-1-AP, Proteintech), mouse monoclonal IgG1 gamma tubulin primary antibodies (T6557, Sigma) and mouse monoclonal IgG2b acetylated tubulin (T6793, Sigma) for 2 hours. Cells were then washed three times with PBS and incubated with Alexa fluor 568 goat anti-rabbit (A11011, Life technologies) and Alexa fluor 633 goat anti-mouse IgG1 (A21126, Life technologies) and Alexa flour 488 goat anti-rabbit (A11008, Life technologies) secondary antibodies for one hour. Cells were finally washed three times with PBS and mounted in Vectashield with DAPI. Images for analysis of ciliogenesis, cilia length and IFT88 localization were taken with a Leica THUNDER Imager (Leica, Wetzlar, Germany) with a 63X objective. Representative images shown in Figure 2 A were taken under identical conditions with Zeiss LSM 880 Observer confocal microscope (Zeiss, Jena, Germany). Images were analysed using ImageJ (version 1.54f; National Institutes of Health, Bethesda, MD, USA). To assess cilia length and ciliation efficiency, at least 100 cells per clone were counted and their cilia measured. For IFT88 localization at least 94 cilia per clone were analyzed and assigned to categories.

**Measurement of Hedgehog pathway response**

Cells were seeded in 6-well plates at a density of 0.35 × 10⁶ cells per well. After reaching confluency, the cells were cultured under low-serum conditions (0.5% FCS) for an additional 48 hours. Following 24 hours of serum starvation, Smoothened agonist SAG (73412, Stem cell Technologies) was added at a final concentration of 500nM, or DMSO was used as a control. After 24 hours of treatment, cells were harvested for western blot analysis.

In order to investigate Hedgehog pathway activation, protein samples were investigated for expression of Gli3 full length (Gli3fl) and Gli3 repressor. Following signal detection, bands for Gli3fl, Gli3 repressor and beta actin were quantified. Gli3repressor/Gli3fl ratio fold reduction was calculated for each sample upon SAG treatment. Gli3fl and Gli repressor expression were normalized to beta actin.

**Western blot**

Cells were harvested by scraping the cells into RIPA lysis buffer (ab156034, abcam) containing a protease inhibitor (11836170001, Roche). After 30 minutes incubation on ice, the lysates were centrifuged at 4°C at 13.000 rpm. Subsequently the supernatant was harvested and the total protein concentration was estimated by performing a Bradford assay (23236, Thermo Fisher Scientific). 30µg of protein was then separated in NuPAGE Bis-Tris gel (4-12%, Invitrogen). Followingly, proteins were blotted onto a Nitrocellulose membrane (1704159, Biorad) in a semi-dry transfer. After blocking the membrane with blocking solution (12010020, Biorad), it was incubated over night at 4°C with the following primary antibodies: anti-Gli3 (1:1000, AF3690, R&D systems), anti-beta actin (1:5000, ab8227, abcam) and anti-sox9 (1:1000, ab5535, abcam). After washing, membranes were incubated for 1 h at RT with HRP-conjugated secondary antibodies: goat anti-rabbit (1:5000, 65-6120, Life Technologies), donkey anti-goat HRP (1:10.000, ab205723, abcam). Signal detection was carried out using enhanced chemiluminescence (RPN2236, Cytiva), and membranes were imaged with a Fusion™ Imaging System (Vilber). The intensity of bands was then quantified by ImageJ in a densitometry analysis.

**RNA sequencing**

RNA was extracted from cells grown in 6-well plates at indicated time points using RNeasy Fibrous Tissue Mini Kit (Qiagen, 74704) according to the protocol provided by the manufacturer. RNA concentration was measured with Nano Drop ND-1000 (Thermo Fisher Scientific, Waltham, MA, USA). RNA quality was assured by measurement of RNA integrity value using High Sensitivity RNA Screen Tape Analysis (Tape Station 4150, Part number 5067 5579, Agilent technologies, Santa Clara, CA, USA). RNA sequencing of three different control clones and three different mutant clones per genotype was performed at Novogene, Cambridge, UK. Two individual samples (d21 IFT43 M1V clone 2, d0 WDR60 nonsense clone 2) showed substantially lower mapping efficiency compared with the remaining samples and were therefore re-submitted for library preparation and sequencing in a separate run. For each sample 5 GB raw data were available.

**Transcriptomics bioinformatics**

RNA-seq data analysis was performed using the Galaxy web platform (usegalaxy.eu, Galaxy). Raw sequencing reads, were first quality-checked using FastQC (version 0.74+galaxy1), and adapter trimming was carried out with Trim Galore! (version 0.6.7+galaxy0). Cleaned reads were then mapped to the mouse reference genome (GRCm39) using RNA STAR (version 2.7.11a+galaxy1) as paired end data and “length of the genomic sequence around annotated junctions” set to 149. Read counts per gene were quantified using featureCounts (version 2.0.6+galaxy0) with GENCODE comprehensive gene annotation for mouse (GRCm39, release M35; Ensembl 112). The resulting count matrix was used as input for differential gene expression analysis in DESeq2 (version 2.11.40.8+galaxy0) within Galaxy. Normalized count files were used for the generation of dot plots in R (version 4.4.0) using RStudio (Posit Software, Boston, MA, USA) and ggplot package (version 4.0.0). Heatmaps were created using Prism10 GraphPad Software Inc.). GO enrichment analysis was performed using DAVID (<https://davidbioinformatics.nih.gov/>) functional annotation tool. All RNA sequencing data generated in this study have been deposited in the Gene Expression Omnibus under accession number GSE326420. The data are currently private and will be made publicly available upon publication.

**ECM proteomics**

ATDC5 cells were cultured in differentiation medium until harvest of ECM. Cell bodies were removed by incubating the sample with extraction buffer (20 mM NH₄OH, 0.5% Triton X-100), followed by extensive PBS washing and 30 min DNAseI (15U/ml, 79254, Qiagen) treatment. Lysis of remaining ECM and preparation for mass spectrometry analysis was performed as described previously^1^. Peptides were analyzed with the Evosep One system (Evosep Biosystems) coupled to a timsTOF fleX mass spectrometer (Bruker). 500 ng of peptides were loaded onto Evotips C18 trap columns (Evosep Biosystems) according to the manufacturer’s protocol. Peptides were separated on an EV1137 performance column (15 cm x 150 µm, 1.5 µm, Evosep) using the standard implemented 30 SPD method with a gradient length of 44 min (buffer A: 0.1% v/v formic acid, dissolved in H_2_O; buffer B: 0.1% v/v formic acid, dissolved in acetonitrile). Over the time of the gradient, the concentration of acetonitrile gradually increased from 0 to 90% at a flow rate of 500 nl/min.

The timsTOF fleX mass spectrometer (Bruker, USA) was operated in the DIA-PASEF mode. DIA MS/MS spectra were collected in the range in an m/z range from 100 to 1700. Ion mobility resolution was set to 0.60–1.60 V·s/cm over a ramp time of 100 ms and an accumulation time of 100 ms. The cycle time was set at 1.8s. The collision energy was programmed as a function of ion mobility, following a straight line from 20 eV for 1/K0 of 0.6 to 59 eV for 1/K0 of 1.6. The TIMS elution voltage was linearly calibrated to obtain 1/K0 ratios using three ions from the ESI-L TuningMix (Agilent) (m/z 622, 922, 1222).

Raw data were analyzed with DIA-NN software (v. 1.8.2 beta 22)^2^. A spectral library was predicted using a FASTA file containing the murine protein sequences as of Jan 31st, 2024 (mouse-EBI-reference database, <https://www.ebi.ac.uk/>). The false discovery rate (FDR) was set to 1%. The search was performed allowing one missed cleavage and cysteine carbamidomethylation enabled as a fixed modification. Match between runs was enabled. Quantification was performed using the label-free quantification algorithm MaxLFQ, which calculates the protein quantities as ratios from all peptide intensities.

The data was further processed and analyzed for the statistical analysis using R (v4.3.0) within R studio. Protein intensities were log2-transformed and median normalized. Differential expression analysis was performed using limma^3^. P-values were corrected by multiple testing using the Benjamini–Hochberg method, as applied by limma. Due to high variability across samples, unadjusted p-values were used for downstream visualization and interpretation. Mass spectrometry raw data have been deposited at the ProteomeXchange Consortium (http://proteomecentral.proteomexchange.org) under the accession number PXD076232. Furthermore, all mass spectrometry proteomics datasets used and/or analysed during this study are available online at the MassIVE repository (http://massive.ucsd.edu/; dataset identifier: MSV000101257; Reviewer account details: Username: “MSV000101257_reviewer”, Password: “ST3771_Schmidts"). This study did not generate new code for analysis.

**Statistical analysis**

For each genotype, at least three independently generated clonal cell lines carrying the same mutation were used. For point mutations, clones shared identical nucleotide changes, whereas for frameshift or nonsense mutations, three independent clones harbouring loss-of-function alleles were analysed. All experiments were independently performed at least three times, except for RNA-seq and proteomics experiments where three independent clonal cell lines per genotype were used. GraphPad Prism 10 software was used to graph, analyze, and present the obtained data. Data are presented as mean ± SD. Statistical analyses were performed using appropriate tests as indicated in the figure legends. Two-group comparisons were conducted using unpaired two-tailed Student’s t-test. Two-way ANOVA followed by Sidak’s or Tukey´s multiple comparisons test was used to perform pairwise comparisons of each mutant to the wild-type control, as indicated in the figure legends. P-values < 0.05 were considered statistically significant.

**Proliferation assay**

Cell proliferation was assessed using the Click-iT™ Plus EdU Alexa Fluor™ 555 Imaging Kit (Thermo Fisher Scientific, Cat. No. C10638) following the manufacturer’s instructions. Briefly, cells were seeded on coverslips and incubated with 10 μM EdU (5-ethynyl-2′-deoxyuridine) for 2 hours at the indicated time points. After incubation, cells were fixed with PFA 5% in PBS, permeabilized with Triton 0.5% in PBS, and subjected to the Click-iT™ Plus reaction cocktail for EdU detection. Cover slips were mounted in Vectashield with DAPI. Fluorescent images were acquired at different spots on the cover slip. EdU incorporation was evaluated by analyzing 3 images, but at least 100 nuclei, per time point and clone.

**Fibronectin staining**

Cells were seeded on gelatin-coated cover slips and were maintained until they reached confluency, then medium was changed to differentiation medium. Cells were washed three times with PBS, fixed with 4% PFA solution. After blocking with 5% BSA in PBS solution the fixed cells were stained with rabbit anti fibronectin antibody (1:500, ab2413, abcam) for 2h. Cells were then washed three times with PBS and incubated with Alexa 488 Goat Anti Rabbit (1:500, A11008, Thermo Fisher Scientific). Cells were finally washed three times with PBS and mounted in Vectashield with DAPI. Images were taken with a Leica THUNDER Imager (Leica, Wetzlar, Germany) with a 63X objective.

Supplementary Tables

Table S1: Overview of mutations in *IFT43*, *WDR60* and *IFT74* causing skeletal dysplasia reported in Pubmed (displayed in Figure S1)

Table S2: Predicted proteins in WDR60 nonsense clones

Table S3: GO enrichment of shared differentially expressed genes between WDR60 A911V and IFT74 del ex 2 vs. wt with higher expression levels at day 14

Table S4: GO enrichment of shared differentially expressed genes between WDR60 A911V and IFT74 del ex 2 vs. wt with lower expression levels at day 14 (top 20, see also Suppl. File 6)

Table S5: GO enrichment WDR60 A911V genes with lower expression levels compared to wt at min. 3 time points

Table S6: GO enrichment WDR60 A911V genes with higher expression levels compared to wt at min. 3 time points

Table S7: Information about CRISPR plasmids used in this study

Table S8: Information about gRNA sequences used in this study

| Mutation | Authors | Reference |
| --- | --- | --- |
| IFT43 c.1A>G / p.Met1Val | Arts et al. 2011 | PMID: 21378380 |
| IFT43 c.2T >A / p.Met1Val | Duran et al. 2017 | PMID: 28400947 |
| IFT43 c.100 G>A / p.Glu34Lys | Biswas et al. 2017 | PMID: 28973684 |
| IFT43 c.520T>C / p.Trp174Arg | Duran et al. 2027 | PMID: 28400947 |
| WDR60 c.44delC / p.Ala15Glufs*90 | Kakar et al. 2018 | PMID: 29271569 |
| WDR60 c.1703−3T>A / p.Ser569Aspfs | McInerney-Leo et al. 2013 | PMID: 23910462 |
| WDR60 c.1777C>T/ p.Arg593Trp | Zhang et al. 2019 | PMID: 29068549 |
| WDR60 c.1891C>T / p.Gln631* | McInerney-Leo et al. 2015 | PMID: 25492405 |
| WDR60 c.1924C>T / p.Arg642* | McInerney-Leo et al. 2015 | PMID: 25492405 |
| WDR60 c.2246C>T / p.Thr749Met | McInerney-Leo et al. 2015 | PMID: 25492405 |
| WDR60 c.2284C>T / p.Arg762* | Zhang et al. 2019 | PMID: 29068549 |
| WDR60 c.2305G>A / p.Glu769Lys | Zhang et al. 2019 | PMID: 29068549 |
| WDR60 c.2503_2505dup / p.Arg835dup | Zhang et al. 2019 | PMID: 29068549 |
| WDR60 c.2789C>T / p.Ser930Leu | Zhao et al. 2022 | PMID: 36381051 |
| WDR60 c.2840 A > T / p.Gln947Leu | Cossu et al. 2016 | PMID: 26874042 |
| WDR60 c.2841del / p.Gln947Hisfs* | Zhang et al. 2019 | PMID: 29068549 |
| WDR60 c.2903C>T / p.Ala968Val | Antony et al. 2022 | https://doi.org/10.1101/2022.03.14.483768 |
| IFT74 c.85C>T / p.Arg29* | Luo et al. 2021 | PMID: 33531668 |
| IFT74 c.92delT / p.Leu31Hfs* | Luo et al. 2021 | PMID: 33531668 |
| IFT74 g.26959922_26962977delinsTTATTATA CTC | Bakey et al. 2023 | PMID: 37315079 |
| IFT74 g.26959922_26962969delinsTTA | Hammarsjö et al. 2021 | PMID: 33875766 |
| IFT74 c.256G > A / p.Gly86Ser and p.Leu77_Gly86del | Yu et al. 2024 | https://doi.org/10.1002/pd.6619 |
| IFT74 c.305+1664delG (splice donor) | Bakey et al. 2023 | PMID: 37315079 |
| IFT74 c.306-24A>G / p.Ser103_Arg135del | Luo et al. 2021 | PMID: 33531668 |
| IFT74 c.371_372del / p.Glu124Argfs* | Kleinendorst et al. 2021 | PMID: 32144365 |
| IFT74 c.535C>G / p.Gln179Glu | Luo et al. 2021 | PMID: 33531668 |
| IFT74 c.789+2T>G (splice donor) | Bakey et al. 2023 | PMID: 37315079 |
| IFT74 c.853G>T / p.Glu285* | Zhongling et al. 2021 | PMID: 34539760 |
| IFT74 c.909dup; p.(Glu304ArgfsTer7) | Yu et al. 2024 | https://doi.org/10.1002/pd.6619 |
| IFT74 c.974+4A>G (splice donor) | Bakey et al. 2023 | PMID: 37315079 |
| IFT74 c.1685-1G>T (splice defect) | Mardy et al. 2021 | PMID: 33748949 |
| IFT74 deletion of exon 14-19 | Lindstrand et al. 2016 | PMID: 27486776 |

Table S1

| genotype | Predicted protein |
| --- | --- |
| *Dync2i1* wild type | MEPGKRRTKDDTWKADDLRKHLKVQSGSPKEEKKLREKKAHKDSESAAPEYREHKSRDPDREARHKEKTAERDLYTSTEHPRGERDRERHKERRKDAKDREKDKLKERHRDQEAEKAHSRGKDREREKDRRARKEEIRQSMAYHDLLSRDMRGRQMAEKVEKKASKIRTEERERRDEDSERIDEDRERRYRERKLQYGDSKEHPLSYWLYKEDGEKKHRKAKDADREKRLREKSSMREKRERHAREKGSSLSDREVEDRHREKRHKEGLHYDDERRRSHADKKERSSKEEHKKRELKELEKEDNDLEATGPDEYLPNLEDDFVDYEDDFEVCDGDDDSNNEHEAREKAEELPLAQKREIQEIQKAISAENERVGELSLKMFQKQGWTEYTKEPWTDANDSPSRTPVCGIFVDFATASHRQKSRSQALKQKTRSSKLLRLIDLDFSFTFSLLDLPPVNEYDMYIRNFGKKNTKQAYVQYNEDNVERDIQTEDIETREVWTQHPGEGTAVSGGSEEKDFSDVTVVPKIDTPRLANFLRAACQVVAVLLEEDRLAAGPSWIPRAQDKALNISDSSSQLNTSLPFLQSRKVSCLHASRVQRQTVVSVHDLPEKAFAPSLDSRHLLCVWDIWQPSGPQKVLICESKVTCCCFSPLKAFLLFAGTVHGSVVVWDLREDSRIHHYVRLSNCFWAFRTPTFSTDGILTSVNHRSPLQAIEPVATSAYKKQSFVLSPFSTQEEMAGLSFHIASLDETGVLNVWVVVELPKADISGSMSDLGLIPGGRIKLVHSTVIQLGNSLSHKDSELWGSTQTLSVKFLPSDPNHFVVGTDMGLISHSTRQDWRVSPRVFKPEQHGVRPIKVNVIDFSPFEETVFLAGCSDGSIRLHQLTSERPIMQWDNSTSGHAVTSLQWSPTRPAVFLVQDDASRIYVWDLLENDLGPVAQQPISPDKLVAMTIVGEPEKTSGSFVALVLARTSGTVDVQNLKRRWTTPAVDEHSQLRLLLQK |
| WDR60 nons. clone 7: *Dync2i1* c.1550_1621del; p.Phe517Cysfs*6 | MEPGKRRTKDDTWKADDLRKHLKVQSGSPKEEKKLREKKAHKDSESAAPEYREHKSRDPDREARHKEKTAERDLYTSTEHPRGERDRERHKERRKDAKDREKDKLKERHRDQEAEKAHSRGKDREREKDRRARKEEIRQSMAYHDLLSRDMRGRQMAEKVEKKASKIRTEERERRDEDSERIDEDRERRYRERKLQYGDSKEHPLSYWLYKEDGEKKHRKAKDADREKRLREKSSMREKRERHAREKGSSLSDREVEDRHREKRHKEGLHYDDERRRSHADKKERSSKEEHKKRELKELEKEDNDLEATGPDEYLPNLEDDFVDYEDDFEVCDGDDDSNNEHEAREKAEELPLAQKREIQEIQKAISAENERVGELSLKMFQKQGWTEYTKEPWTDANDSPSRTPVCGIFVDFATASHRQKSRSQALKQKTRSSKLLRLIDLDFSFTFSLLDLPPVNEYDMYIRNFGKKNTKQAYVQYNEDNVERDIQTEDIETREVWTQHPGEGTAVSGGSEEKDCGCCIA* |
| WDR60 nons. clone 14: *Dync2i1* c.1582_1583del; pThr528Serfs*4 | MEPGKRRTKDDTWKADDLRKHLKVQSGSPKEEKKLREKKAHKDSESAAPEYREHKSRDPDREARHKEKTAERDLYTSTEHPRGERDRERHKERRKDAKDREKDKLKERHRDQEAEKAHSRGKDREREKDRRARKEEIRQSMAYHDLLSRDMRGRQMAEKVEKKASKIRTEERERRDEDSERIDEDRERRYRERKLQYGDSKEHPLSYWLYKEDGEKKHRKAKDADREKRLREKSSMREKRERHAREKGSSLSDREVEDRHREKRHKEGLHYDDERRRSHADKKERSSKEEHKKRELKELEKEDNDLEATGPDEYLPNLEDDFVDYEDDFEVCDGDDDSNNEHEAREKAEELPLAQKREIQEIQKAISAENERVGELSLKMFQKQGWTEYTKEPWTDANDSPSRTPVCGIFVDFATASHRQKSRSQALKQKTRSSKLLRLIDLDFSFTFSLLDLPPVNEYDMYIRNFGKKNTKQAYVQYNEDNVERDIQTEDIETREVWTQHPGEGTAVSGGSEEKDFSDVTVVPKIDSEVG* |
| WDR60 nons. clone 20: *Dync2i1* c.1583del; pThr528Ilefs*47 | MEPGKRRTKDDTWKADDLRKHLKVQSGSPKEEKKLREKKAHKDSESAAPEYREHKSRDPDREARHKEKTAERDLYTSTEHPRGERDRERHKERRKDAKDREKDKLKERHRDQEAEKAHSRGKDREREKDRRARKEEIRQSMAYHDLLSRDMRGRQMAEKVEKKASKIRTEERERRDEDSERIDEDRERRYRERKLQYGDSKEHPLSYWLYKEDGEKKHRKAKDADREKRLREKSSMREKRERHAREKGSSLSDREVEDRHREKRHKEGLHYDDERRRSHADKKERSSKEEHKKRELKELEKEDNDLEATGPDEYLPNLEDDFVDYEDDFEVCDGDDDSNNEHEAREKAEELPLAQKREIQEIQKAISAENERVGELSLKMFQKQGWTEYTKEPWTDANDSPSRTPVCGIFVDFATASHRQKSRSQALKQKTRSSKLLRLIDLDFSFTFSLLDLPPVNEYDMYIRNFGKKNTKQAYVQYNEDNVERDIQTEDIETREVWTQHPGEGTAVSGGSEEKDFSDVTVVPKIDIRGWLTFSGQRARWLLYCLKRTVWQLDPAGFPGLKTKLSTSVTAHLN* |

Table S2

| Term | Genes | FDR |
| --- | --- | --- |
| GO:0030282~bone mineralization | SMPD3, PHOSPHO1, WNT11, IBSP, ALPL, AXIN2, PTH1R, PHEX, IFITM5 | 3,99E-09 |
| GO:0001958~endochondral ossification | SMPD3, PHOSPHO1, MEF2C, COL2A1, TMEM119, ALPL, BMP6 | 1,91E-07 |
| GO:0051216~cartilage development | SMPD3, COL2A1, COL11A1, COL11A2, HOXA3, SOX6, BMP6 | 3,24E-05 |
| GO:0001649~osteoblast differentiation | MEF2C, WNT11, COL11A2, TMEM119, AXIN2, BMP6, PANX3 | 7,88E-05 |
| GO:0001501~skeletal system development | SMPD3, COL2A1, COL27A1, COL11A2, PTH1R, IFITM5 | 2,32E-03 |
| GO:0002062~chondrocyte differentiation | MEF2C, COL2A1, COL11A2, PTH1R, SOX6 | 3,27E-03 |
| GO:0001503~ossification | SMPD3, COL2A1, IBSP, COL11A1, PTH1R, BMP6 | 5,02E-03 |
| GO:0060021~roof of mouth development | MEF2C, COL2A1, WNT11, COL11A2, TBX2 | 5,02E-03 |
| GO:0030198~extracellular matrix organization | VIT, COL2A1, COL27A1, IBSP, COL11A1, NDNF | 6,93E-03 |
| GO:0003139~secondary heart field specification | MEF2C, WNT11, AXIN2 | 2,39E-02 |
| GO:0010629~negative regulation of gene expression | C1QTNF3, MEF2C, CDKN1A, ATP2B4, MAEL, SLIT3, KLF4 | 2,85E-02 |
| GO:0010467~gene expression | MEF2C, HOXA3, MAEL, FLVCR2, SOX6, KLF4, SGCG | 3,35E-02 |
| GO:1901653~cellular response to peptide | SMPD3, KLF4, KLF2 | 4,50E-02 |

Table S3

| Term | Genes | FDR |
| --- | --- | --- |
| GO:0045087~innate immune response | APOL9B, UBA7, IFI211, OAS1B, IFIT1, IFIT3, IFIH1, C7, CASP4, DHX58, TNFRSF14, LBP, GBP2, 9930111J21RIK2, 9930111J21RIK1, GBP3, ZBP1, GBP5, GBP2B, GBP7, IFI204, DTX3L, SP110, MX2, ANG2, IRGM2, IRGM1, IFI202B, PARP9, PARP14, GM12250, IGTP, IRF7, IFIT3B, TRIM14, IRF5 | 8,07E-29 |
| GO:0035458~cellular response to interferon-beta | GBP2B, GBP7, IFI204, IFI205, IFI203, IFI211, IRGM2, IRGM1, OAS1B, IFI202B, IFIT1, IFIT3, GM12250, MNDAL, IGTP, GBP2, 9930111J21RIK2, 9930111J21RIK1, GBP3 | 4,62E-26 |
| GO:0002376~immune system process | H2-T24, SLFN2, H2-Q6, H2-Q4, IFI211, OAS1B, IFIT1, IFIT3, IFIH1, C7, CASP4, DHX58, TNFRSF14, LBP, GBP2, GBP3, ZBP1, GBP5, GBP2B, GBP7, IFI204, DTX3L, SP110, MX2, IRGM2, IRGM1, IFI202B, PARP9, PARP14, GM12250, IGTP, IRF7, TRIM14, IRF5, IRF9 | 1,01E-22 |
| GO:0051607~defense response to virus | ZBP1, RTP4, GBP5, GBP2B, GBP7, DTX3L, MX2, OAS1B, IFIT1, PARP9, IFIT3, GM12250, IFIH1, CXCL10, DHX58, IRF7, IFIT3B, IRF5, GBP2 | 6,40E-16 |
| GO:0002218~activation of innate immune response | ZBP1, GBP5, GBP2B, GBP7, IFI204, IFI205, IFI203, IFI211, IFI202B, GBP2, MNDAL, GBP3 | 1,24E-14 |
| GO:0009617~response to bacterium | GBP5, SLFN2, IFI204, IFI205, SP110, MX2, IFI211, IRGM2, IRGM1, IFI44, IFIT1, USP18, IFIT3, GM12250, CXCL10, DHX58, LY6A, GBP2 | 7,95E-14 |
| GO:0051715~cytolysis in another organism | GBP5, GBP2B, GBP7, IGTP, GBP2, GBP3 | 9,38E-09 |
| GO:0042832~defense response to protozoan | GBP2B, GBP7, IGTP, IRGM2, GBP2, IRF9, GM12250, GBP3 | 2,85E-08 |
| GO:0042742~defense response to bacterium | GBP5, GBP2B, GBP7, IRGM2, IRGM1, OAS1B, GM12250, CASP4, IGTP, TNFRSF14, LBP, GBP2, GBP3 | 3,13E-08 |
| GO:0140374~antiviral innate immune response | IFIH1, ZBP1, CXCL10, DHX58, IFIT3B, OAS1B, USP18, IFIT3 | 3,05E-06 |
| GO:0160075~non-canonical inflammasome complex assembly | GBP5, GBP2B, CASP4, GBP2, GM12250 | 3,63E-06 |
| GO:0140639~positive regulation of pyroptotic inflammatory response | GBP5, GBP2B, GBP2, GM12250, GBP3 | 3,63E-06 |
| GO:0031347~regulation of defense response | GBP5, GBP2B, GBP7, GBP2, GBP3 | 3,63E-06 |
| GO:0098586~cellular response to virus | IFIH1, CXCL10, IGTP, IRGM2, IRGM1, IFI44, IRF5 | 4,65E-06 |
| GO:0051707~response to other organism | IFIT3B, IFI44, IFIT1, USP18, IFIT3 | 4,50E-05 |
| GO:0035457~cellular response to interferon-alpha | IFI204, IFI211, IFIT3B, OAS1B, IFIT1 | 1,09E-04 |
| GO:0071346~cellular response to type II interferon | GBP5, GBP2B, GBP7, IGTP, IRGM1, GBP2, GBP3 | 1,12E-04 |
| GO:0060335~positive regulation of type II interferon-mediated signaling pathway | IGTP, IRGM2, IRGM1, PARP9 | 1,23E-04 |
| GO:0044406~adhesion of symbiont to host | GBP2B, GBP7, GBP2, GBP3 | 1,74E-04 |
| GO:0009615~response to virus | IFIH1, MX2, DHX58, IFIT3B, OAS1B, IFIT1, IFIT3 | 1,91E-04 |

Table S4

| term | genes | FDR |
| --- | --- | --- |
| GO:0051607~defense response to virus | IFITM3, RTP4, TRIM12C, OAS1A, OAS1B, DDX60, IFIT1, IFIT3, OAS1G, MNDAL, IFIH1, DHX58, GBP2, OASL1, IFITM6, OASL2, IL33, GBP5, GBP2B, IFI207, DTX3L, MX2, IFI203, BST2, CXCL10, PLSCR1, OAS3, IRF1, IRF7, IFIT3B, IRF5, TRIM34B, TRIM34A | 3,78E-23 |
| GO:0035458~cellular response to interferon-beta | TGTP2, GBP2B, IFI207, IFI204, IFI205, IFI203, IFI211, IRGM2, IRGM1, OAS1A, IFI47, OAS1B, IFIT1, IFIT3, MNDAL, OAS1G, TRIM6, IRF1, IGTP, GBP2, 9930111J21RIK2, GBP3 | 5,78E-23 |
| GO:0045087~innate immune response | TRIM12C, TRIM30A, UBA7, OAS1A, OAS1B, IFIT1, IFIT3, OAS1G, IFIH1, TRIM6, CASP4, DHX58, C1RL, TNFRSF14, LBP, OASL1, 9930111J21RIK2, OASL2, TGTP2, IFI204, DTX3L, SP110, MX2, ANG2, IRGM2, IRGM1, IFI47, PARP14, BST2, VNN1, OAS3, IGTP, TRIM14, IRF5, TRIM34B, TMEM106A, TRIM34A | 4,76E-18 |
| GO:0045071~negative regulation of viral genome replication | IFITM3, IFI207, MX2, IFI203, OAS1A, OAS1B, MNDAL, OAS1G, IFIH1, BST2, TRIM6, PLSCR1, OAS3, CCL5, OASL1, IFITM6, OASL2 | 2,93E-17 |
| GO:0009615~response to virus | IFITM3, MX2, OAS1A, OAS1B, DDX60, IFIT1, IFIT3, OAS1G, IFIH1, BST2, OAS3, DHX58, IFIT3B, IFI27L2A, OASL1, OASL2 | 5,61E-11 |
| GO:0140374~antiviral innate immune response | OAS1A, OAS1B, USP18, IFIT3, OAS1G, IFIH1, CXCL10, TRIM6, OAS3, DHX58, IFIT3B, OASL1, OASL2 | 8,70E-10 |
| GO:0002218~activation of innate immune response | TRIM12C, GBP5, GBP2B, TRIM30A, IFI207, IFI204, IFI205, IFI203, IFI211, GBP2, MNDAL, GBP3 | 1,55E-09 |
| GO:0006955~immune response | H2-T24, CD74, IL15, P2RY14, H2-Q4, IFI44, SERPINB9, AFP, TNFRSF1B, CX3CL1, CXCL10, PLSCR1, RAET1E, CXCL12, CCL7, SERPINB9D, SERPINB9E, PLSCR2, SERPINB9G, TNFSF10, CCL2, CTSH, SERPINB9B, SERPINB9C | 4,80E-08 |
| GO:0009617~response to bacterium | GBP5, TRIM30A, IFI204, IFI205, SP110, MX2, IRGM2, IRGM1, IFI44, VEGFD, IFIT1, USP18, IFIT3, NAALADL2, CXCL10, DHX58, CCL2, LY6A, GBP2 | 5,11E-08 |
| GO:0042742~defense response to bacterium | GBP5, GBP2B, LYZ2, IRGM2, STAB2, IRGM1, OAS1A, SERPINB9, OAS1B, OAS1G, PLAC8, OAS3, CASP4, IGTP, TNFRSF14, GBP2, GBP3 | 5,54E-07 |
| GO:0060337~type I interferon-mediated signaling pathway | IFIH1, IFITM3, OAS3, IRF7, OAS1A, OAS1B, OASL1, OAS1G, IFITM6, OASL2 | 5,54E-07 |
| GO:0010951~negative regulation of endopeptidase activity | SERPINB1A, SERPINB9D, SERPINB6B, SERPINB9E, SERPINB9G, SERPINB9B, SERPINB9, SERPINB9C | 2,65E-06 |
| GO:0006954~inflammatory response | GBP5, VCAM1, PLA2G2E, F2R, TNFRSF1B, KNG2, CX3CL1, CXCL10, SERPINB1A, IL1RL1, VNN1, CCL7, NAGLU, CCL5, CHST1, CCL2, TRIM14, ADAM8, IRF5 | 1,36E-05 |
| GO:0098586~cellular response to virus | IFIH1, CXCL10, TRIM6, CCL5, IGTP, IRGM2, IRGM1, IFI44, IRF5 | 1,47E-05 |
| GO:0070106~interleukin-27-mediated signaling pathway | OAS3, OAS1A, OAS1B, OASL1, OAS1G, OASL2 | 2,27E-05 |
| GO:0042270~protection from natural killer cell mediated cytotoxicity | SERPINB9D, SERPINB9E, SERPINB9G, SERPINB9B, SERPINB9, SERPINB9C | 2,27E-05 |
| GO:0051715~cytolysis in another organism | GBP5, GBP2B, IGTP, GBP2, GBP3 | 2,79E-05 |
| GO:0071222~cellular response to lipopolysaccharide | GBP5, GBP2B, IRGM2, IRGM1, TNFRSF1B, CXCL10, PLSCR1, PLSCR2, IGTP, PDE4B, ANKRD1, CCL2, CMPK2, LBP, GBP2, GBP3 | 3,09E-05 |
| GO:0042832~defense response to protozoan | TGTP2, GBP2B, IGTP, IRGM2, IL4RA, GBP2, GBP3 | 2,94E-04 |
| GO:0043066~negative regulation of apoptotic process | ANGPT4, CD74, ARNT2, PLK2, CTNS, SERPINB9, IGF1, IFIT3, CX3CL1, PLAC8, CD59B, CD59A, GRK5, SERPINB9D, SERPINB9E, SERPINB9G, IFIT3B, CTSH, SERPINB9B, SERPINB9C, NOS1 | 6,56E-04 |

Table S5

| Term | Genes | FDR |
| --- | --- | --- |
| GO:0001958~endochondral ossification | SMPD3, PHOSPHO1, MEF2C, COL13A1, GALNT3, DLX5, TMEM119, ALPL | 1,06E-07 |
| GO:0030282~bone mineralization | SMPD3, PHOSPHO1, COL1A2, WNT11, ALPL, AXIN2, PTH1R, FGFR3 | 3,06E-06 |
| GO:0008285~negative regulation of cell population proliferation | CDKN1A, GPER1, TESC, ZBTB16, CTH, P3H2, SLIT3, AXIN2, PTH1R, KLF4, FGFR3, RERG | 8,63E-05 |
| GO:0030198~extracellular matrix organization | VIT, COL15A1, COL1A2, COL13A1, COL22A1, COL11A1, ADAMTSL2, NDNF | 1,28E-03 |
| GO:0030154~cell differentiation | MEF2A, MEF2C, CADM1, COL13A1, DLX5, TESC, ARHGAP24, CLGN, NR4A1, GPER1, SOX6, AGRN, FGFR3 | 1,28E-03 |
| GO:0001501~skeletal system development | SMPD3, COL1A2, COL13A1, ZBTB16, CHD7, PTH1R, HAPLN4 | 1,38E-03 |
| GO:0051216~cartilage development | SMPD3, ZBTB16, COL11A1, HOXA3, SOX6, FGFR3 | 3,79E-03 |
| GO:0051480~regulation of cytosolic calcium ion concentration | GPER1, HRC, ATP2B4, PLCD1, BOK | 3,79E-03 |
| GO:0001649~osteoblast differentiation | MEF2C, WNT11, DLX5, TMEM119, AXIN2, SP7 | 8,39E-03 |
| GO:0030308~negative regulation of cell growth | CDKN1A, WNT11, CTH, SLIT3, CRYAB, RERG | 1,30E-02 |
| GO:0009887~animal organ morphogenesis | WNT11, HHIP, PDGFC, HOXA3, PDGFA, SLIT3 | 1,30E-02 |
| GO:0060025~regulation of synaptic activity | MEF2C, AGRN, SYBU | 1,70E-02 |
| GO:0060348~bone development | SMPD3, CADM1, PDGFC, PDGFA, FGFR3 | 3,36E-02 |
| GO:0007160~cell-matrix adhesion | COL13A1, ITGA10, ITGA7, HPSE, NID2 | 3,36E-02 |
| GO:0045893~positive regulation of DNA-templated transcription | MEF2A, NR4A1, MEF2C, WNT11, DLX5, BAMBI, TESC, ZBTB16, SOX6, KLF4, KLF2 | 4,05E-02 |
| GO:0010629~negative regulation of gene expression | MEF2C, CDKN1A, GPER1, ATP2B4, SLIT3, KLF4, FGFR3, CRYAB | 4,05E-02 |
| GO:0003139~secondary heart field specification | MEF2C, WNT11, AXIN2 | 4,05E-02 |
| GO:0055074~calcium ion homeostasis | HRC, TMTC2, ALPL, FGFR3 | 4,05E-02 |
| GO:0060070~canonical Wnt signaling pathway | WNT11, SIAH2, TCF7, AXIN2, KLF4 | 4,05E-02 |
| GO:0007417~central nervous system development | CYP26B1, ZBTB16, CHD7, SOX6, HAPLN4 | 4,05E-02 |

Table S6

| **IFT or dynein component** | **Mutation** | **Base changes** | **CRISPR plasmid** | **source** |
| --- | --- | --- | --- | --- |
| IFT-A | IFT43 M1V | A 🡪 G | pCMV_ABEmax_P2A_GFP | Addgene Plasmid #112101 |
| IFT-B | IFT74 del exon 2 | Frameshift | pSpCas9(BB)-2A-GFP (PX458) | Addgene Plasmid #48138 |
| Dynein-2 | WDR60 A911V | C 🡪 T | pCMV_BE4max_P2A_GFP | Addgene Plasmid #112099 |
| Dynein-2 | WDR60 KO | Frameshift | Cas9 expression vector pRP[Exp]-mCherry/  Hygro-CBh>hCas9 | Vectorbuilder VB010000-9378bvk |

Table S7

| **IFT/dynein component** | **Mutation** | **gRNA sequence 5´🡪3´** | **PAM** | **complementary strand** |
| --- | --- | --- | --- | --- |
| IFT-A | IFT43 M1V | GGCGATGGACGATTTAC | TGG | reverse |
| IFT-B | IFT74 del exon 2 | GGCTGAAGATTTGTGATTAC | CCA | forward |
| Dynein-2 | WDR60 A911V | CCTGCAGTGTTCCTGGTCC | AGG | reverse |
| Dynein-2 | WDR60 KO | TACCGAAGATCGACACTCCG | AGG | reverse |

Table S8
